# Integrative analysis of angiogenic signaling in obesity: capillary features and VEGF binding kinetics

**DOI:** 10.1101/2024.12.23.630107

**Authors:** Yunjeong Lee, Keith Lionel Tukei, Yingye Fang, Shobhan Kuila, Xinming Liu, Princess I. Imoukhuede

## Abstract

Obesity is a global health crisis, with its prevalence particularly severe in the United States, where over 42% of adults are classified as obese. Obesity is driven by complex molecular and tissue-level mechanisms that remain poorly understood. Among these, angiogenesis—primarily mediated by vascular endothelial growth factor (VEGF-A)—is critical for adipose tissue expansion but presents unique challenges for therapeutic targeting due to its intricate regulation. Systems biology approaches have advanced our understanding of VEGF-A signaling in vascular diseases, but their application to obesity is limited by scattered and sometimes contradictory data. To address this gap, we performed a comprehensive analysis of the existing literature to synthesize key findings, standardize data, and provide a holistic perspective on the adipose vascular microenvironment. The data mining revealed five key findings: (1) obesity increases adipocyte size by 78%; (2) vessel density in adipose tissue decreases by 51% in obese mice, with vessels being 47–58% smaller and 4–9 times denser in comparison with tumor vessels; (3) capillary basement membrane thickness remains similar regardless of obesity; (4) VEGF-A shows the strongest binding affinity for VEGFR1, with four times stronger affinity for VEGFR2 than for NRP1; and (5) binding affinities measured by radioligand binding assay and surface plasmon resonance (SPR) are significantly different. These consolidated findings provide essential parameters for systems biology modeling, new insights into obesity-induced changes in adipose tissue, and a foundation for developing angiogenesis-targeting therapies for obesity.

## Introduction

Angiogenesis, primarily driven by vascular endothelial growth factor-A (VEGF-A, i.e., VEGF-A165), is essential for healthy adipose tissue expansion and remodeling^1^. Insufficient vascularization causes unhealthy white adipose tissue expansion in obesity, leading to fibrosis, inflammation, and systemic insulin resistance in preclinical models^2–5^. Therapeutic modulation of angiogenesis remains controversial because targeting the primary molecular drivers of vascularization, the VEGF family, does not fully control obesity^6,7^. This is despite the numerous studies establishing the relationship between VEGF signaling and adipose tissue development, which have found that (1) VEGF-A mRNA expression is upregulated during adipocyte differentiation in vitro^8^, (2) overexpression of VEGF-A in adipose tissue not only promotes angiogenesis but also improves the metabolic system^9^, and (3) anti-VEGF suppresses both angiogenesis and the formation of differentiating adipocytes^10^. Despite these discoveries, how to control adipogenesis via VEGF-A signaling is still unclear. This knowledge gap can be addressed by using systems biology to describe the fundamental processes governing vascularization in obesity with mathematical models and experimental data, simulating them with computational approaches, and analyzing the system to identify the best approaches to addressing insufficient vascularization in obesity.

Although several mathematical models have been developed to explain VEGF-A signaling in cancer or peripheral arterial disease, they are hard to employ in developing obesity-specific models because of the following distinctive features of adipose tissue: (1) an extremely high proportion of adipocytes in adipose tissue volume^11^ that is not present in the established muscle^12^ or tumor models^13–15^ (Figure 1A), (2) phenotypically different capillaries in adipose tissue compared with unorganized tumor capillaries, and (3) different levels of the molecular drivers of angiogenesis, VEGFRs, in adipose tissue (Figure 1B)^13,16^. Thus, there is a need to develop adipose tissue–specific computational models, and these require adipose tissue–specific data and mathematical representations.

**Figure 1.**
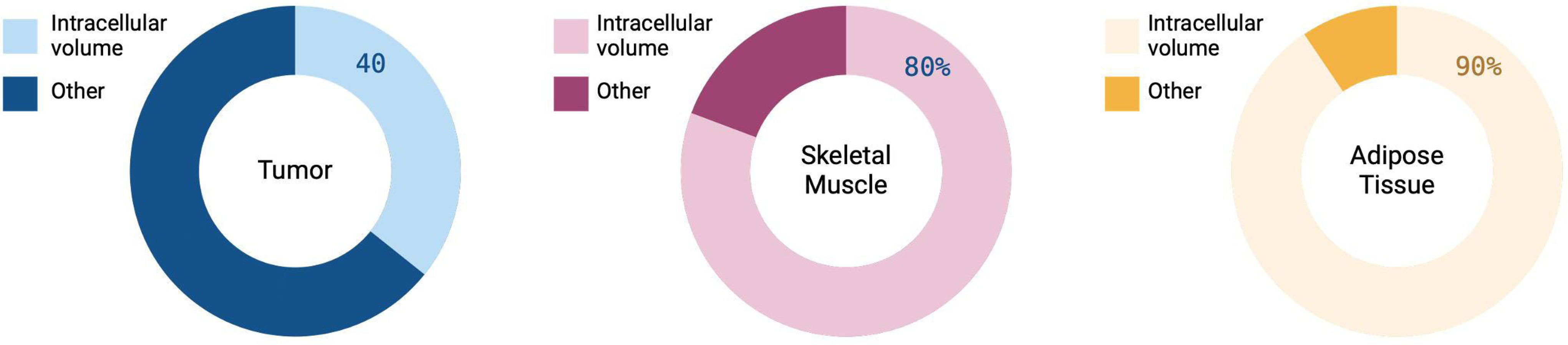
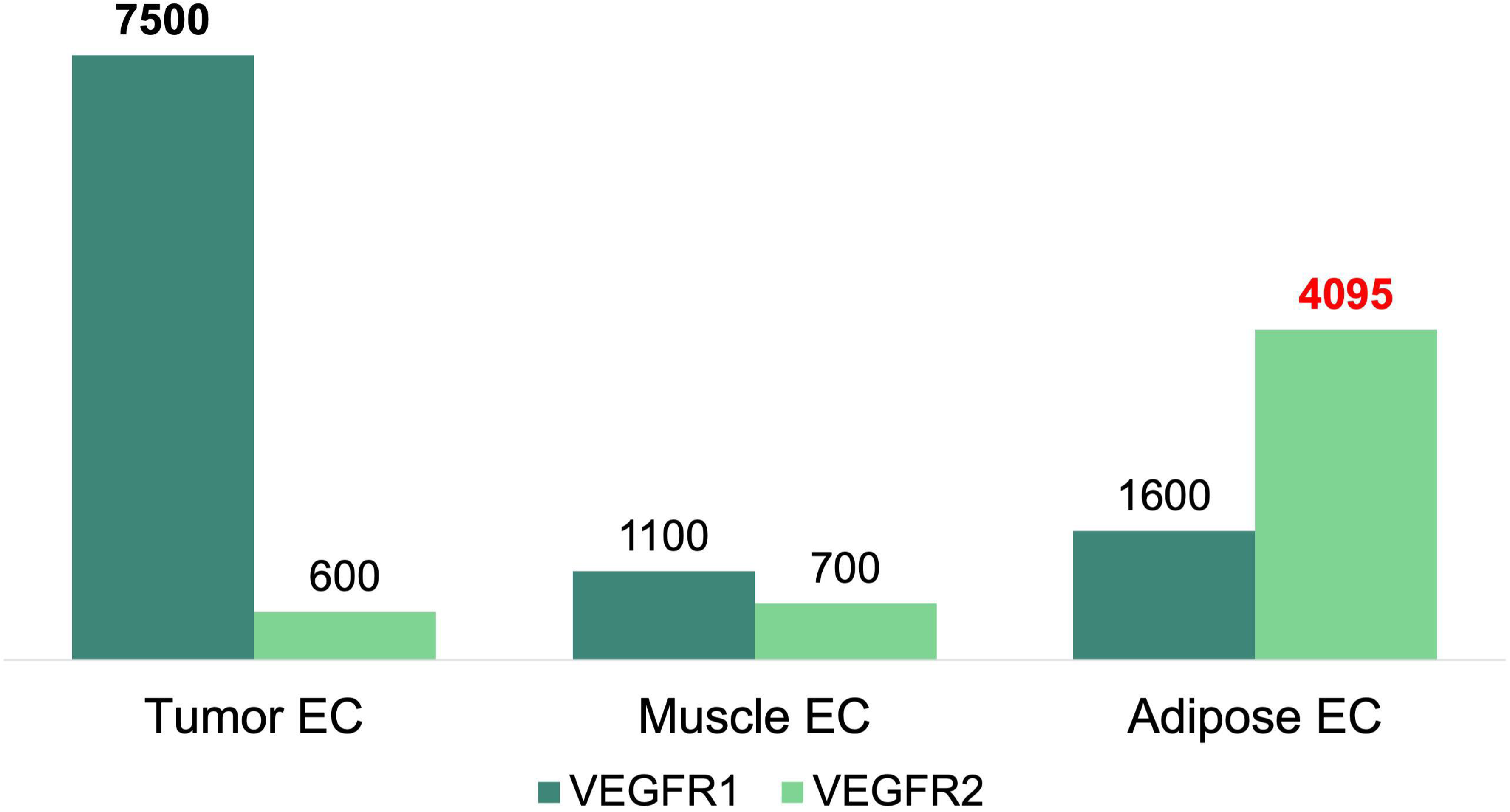
Distinctive characteristics of adipose tissue compared with tumor and muscle. (A) Proportions of tissue volume that are intracellular vs. extracellular in tumor, muscle, and adipose tissue (ref. from Arner & Rydén^11^, Yen et al.^12^, Finley et al.^13^, Hao et al.^14^, and Del Monte^15^). The adipose tissue has a significantly large proportion of intracellular volume compared with other tissues. (B) The number of receptors on endothelial cells (EC) measured in tumor, muscle, and adipose tissue (ref. from Finley et al.^13^ and Fang et al.^16^). Tumor has a high level of VEGFR1, muscle has similar levels of VEGFR1 and VEGFR2, and adipose tissue has a high level of VEGFR2.

To address these needs for adipose tissue–specific data on vascular and adipocyte properties, we performed an in-depth analysis of the experimental literature, identifying data governing adipocyte and vascular morphology and data that standardize VEGFR binding kinetics. These data offer new insights into the adipose-tissue vascular microenvironment, reflecting the differential changes occurring as a result of obesity. They will accelerate the use of systems biology to mathematically represent, model, and predict approaches for treating obesity.

## Methods

### Search Strategy

We mined the literature using PubMed, Google Scholar, and the University of Washington Libraries. Search keywords included adipocyte size, high-fat diet mice, gonadal adipose tissue, vessel size, vessel diameter, capillary diameter, vessel density, tumor, retina capillary basement membrane thickness, muscle capillary basement membrane thickness, brain capillary basement membrane thickness, thickness of renal glomerular capillary basement membrane, VEGF, VEGF-A, VEGF165, VEGF-A165, VEGFR1, Flt-1, VEGFR2, KDR, NRP1, surface plasmon resonance, SPR, radioligand, radioactivity, 123I, 125I, binding affinity, and K_d_.

### Study selection and eligibility criteria

We included *ex vivo* studies that measured adipocyte diameter or the cross-sectional area of an adipocyte, the cross-sectional area of vessels (vessel size), and vessel densities per mm^2^ in gonadal adipose tissue of lean and obese mice. Since most computational models for VEGF signaling have been developed in tumors, identifying potential differences between tumors and adipose tissue is an important goal. Thus, data for vessel size and vessel density in mouse tumors were included in our analysis. We also included *ex vivo* studies reporting the capillary basement membrane thickness in retina, muscle, heart, brain, and kidney from mice and rats. To analyze the binding kinetics of VEGF-A to its receptors, we included *in vitro* studies that measured binding affinities (K_d_), association rate constants (k_on_), and dissociation rate constants (k_off_) using radioligand assays and surface plasmon resonance. We excluded papers if they (1) did not provide enough information to extract average values and standard errors (e.g., the standard deviation was reported without sample size or only the average value was reported) or (2) reported very large values considered as outliers (e.g., the vessel size in tumors reported by Koyama *et al.*^17^ is 10 times larger than values in other studies). We did not include non-English papers.

We did not analyze VEGF-B and VEGF-C, although they influence lipid metabolism and lymphangiogenesis, which are important mechanisms in obesity, because of a lack of information about their binding affinities. Instead, we gathered and reported a list of binding data for VEGF-B and VEGF-C to VEGFRs regardless of measurement method.

### Data extraction

The data were extracted from the full text or supplementary materials of the selected papers. We used minor data extraction techniques in the following cases. (1) When information about average values and standard errors was provided by an image, we used ImageJ V1.53k (https://imagej.net/) to extract that information^18^. (2) When papers provided a graph showing association and dissociation phases of VEGF-A binding without providing values, we estimated kinetic data by fitting 1:1 Langmuir equations to the provided graph. This data extraction process was performed by using an in-house code written in Python Programming Language V3.11.0 (http://www.python.org/). The detailed approaches for data extraction, data fitting, and ImageJ analysis are provided in the *Supplementary information*.

All data sets include the following information: the first author’s name, year published, data collection techniques, measurements (i.e., means), and standard errors. For geometric data, we additionally extracted the following: species, strain, sample size, sex, age, diet or status (e.g., healthy or diabetic mice), duration of diet, body weight, and location of tissue. For tumor data, we included tumor cell lines, the location in the mouse body where the tumors were injected, and antibodies used for vessel staining. For binding affinity data, we extracted the ligands used, the receptors, and their sources. The data extraction was done independently by two authors (Yunjeong Lee and Keith Lionel Tukei), and any disagreement was resolved by discussion among three authors (Yunjeong Lee, Keith Lionel Tukei, and Shobhan Kuila).

### Data analysis

The weighted average and standard deviation of each group were calculated with a random-effects model. We assumed that the analyzed data followed normal distributions. The weight of the *i*^th^ study was defined by *1/(SE_i_^2^ + T^2^)*, where *SE_i_*is the standard error of the *i*^th^ study and *T^2^*is the between-study variance. Cochran’s Q-test, the most common way to assess the presence of heterogeneity between studies, was performed. Two statistics evaluated the level of heterogeneity: (1) I^2^, which represents the proportion of variation between studies among the total variation, and (2) prediction intervals, which show the amount of dispersion of the observed measurements^19^.

After calculating the weighted average and standard deviations of groups, we used Welch’s t-test to compare a pair of two groups because (1) the groups are unpaired, (2) the groups have unequal variances, and (3) the analyzed data were assumed to follow the normal distribution. When we performed a one-tailed test to identify inequality between more than two groups, we used a one-tailed Welch’s t-test with the Benjamini-Hochberg method for multiple testing. The Benjamini-Hochberg method was chosen since it is balanced between Type I and Type II errors and its statistical power is stronger than that of the Bonferroni method, a commonly used multiple-testing correction. The Bonferroni method is more conservative because it divides the p-value by the number of tests, and it is recommended when the cost of Type I errors is more expensive than Type II errors^20^. Since the cost of Type I errors in our analysis is similar to that of Type II errors, we chose the Benjamini-Hochberg method. When we compared more than two groups, the homogeneity of variances was assessed by Bartlett’s test since it (1) is appropriate when the sample sizes are unequal, and (2) is generally powerful for various variance ratios between groups. After testing the homogeneity of variance, Welch’s ANOVA followed by Dunnett’s T3 test was used to compare means of multiple groups. Dunnett’s T3 test was chosen because it is appropriate for a small sample dataset. All statistical analysis was performed using the “metafor”, “weights”, “stats”, and “PMCMRplus” libraries in R Project for Statistical Computing, RStudio V4.3.0 (http://www.r-project.org/). The full codes and Excel files are available on the Open Science Framework (doi: 10.17605/OSF.IO/N9CEY).

### Surface Plasmon Resonance (In Vitro)

Surface plasmon resonance (SPR) experiments were performed at 25°C using the Reichert 4SPR system (Reichert, Inc., USA) with PEG-coated gold sensor chips containing 10% COOH (Reichert, Inc., USA #13206061). The chip was divided into four flow cells: growth factors were immobilized in channels 1 or 3, leaving channels 2 or 4 blank as references. The running buffer was 1x HBS-EP pH 7.4 (10 mM HEPES, 3 mM EDTA, 150 mM NaCl, 0.005% Tween-20). The ligands NRP1 (Cat. #3870-N1-025/CF, R&D Systems) and VEGFR2 (Cat. #357-KD-050/CF, R&D Systems) were immobilized using an amine coupling method. EDC (1-ethyl-3-[3-dimethylaminopropyl]carbodiimide, 40 mg/mL), and NHS (N-hydroxysuccinimide, 10 mg/mL) were dissolved in water, mixed, and injected at 10 μL/min for 7 minutes to activate the surface. Proteins were diluted to 30 μg/mL in 10 mM acetate buffer (pH 4.0) and injected at 10 μL/min until the immobilization level reached ≥2000 RU, based on an Rmax target of less than 200 RU to minimize mass transfer effects. The surface was deactivated by injecting 1M ethanolamine hydrochloride-NaOH (pH 8.5) for 7 minutes at 10 μL/min.

For kinetic analysis, analyte VEGF-A (Cat. #293-VE-010, R&D Systems) was injected at concentrations of 50, 25, 12.5, 6.25, and 3.12 nM, and the association and dissociation curves were fitted using a 1:1 Langmuir binding model in TraceDrawer ver.1.8.1. Sensorgrams were visually inspected, and the fitting was validated by the χ2-to-Rmax ratio (<0.10), ensuring a reliable 1:1 interaction model. Raw sensorgrams (3.12–50 nM) were aligned, and nonspecific binding was subtracted using reference channel sensorgrams. Global fitting, considered more accurate than single-curve fitting, was applied using nonlinear least-squares analysis in TraceDrawer to determine association (k_on_) and dissociation (k_off_) rates across multiple response curves. The results are presented as mean ± standard error.

## Results

### Study selection

A total of 76 studies were analyzed (Figure 2; *n* = the number of studies): *n* = 10 for adipocyte size; *n* = 11 for vessel size; *n* = 12 for vessel density; *n* = 34 for capillary basement membrane thickness; *n* = 8 for VEGF-A binding affinity to VEGFR1; *n* = 12 for VEGF-A binding affinity to VEGFR2; *n* = 10 for VEGF-A binding affinity NRP1; *n* = 3 for VEGF-A to VEGFR1 association and disassociation rates; *n* = 6 for VEGF-A to VEGFR2 association and disassociation rates; and *n* = 3 for VEGF-A to NRP1 association and dissociation rates. After assessing full-text articles, we included only studies that provided information enabling the extraction of average values and standard errors, since those values were essential in the analysis. A total of 21 studies were excluded from the final selection. The rationales for their exclusion were as follows:

1. *Lack of data to extract mean ± standard error (15 studies)*: Lang *et al.*^21^, Sletta *et al.*^22^, Cai *et al.*^23^, Bloodworth *et al.*^24^, Welt *et al.*^25^, Colotti *et al.*^26^, Geretti *et al.*^27^, Guerrin *et al.*^28^, Mamluk *et al.*^29^, Papo *et al.*^30^, Shibuya *et al.*^31^, Soker *et al.*^32^, Tillo *et al.*^33^, Pan *et al.*^34^, and Vintonenko *et al.*^35^ were excluded because the mean or standard error could not be extracted.
2. *Outliers (3 studies)*: Koyama *et al.*^17^ was excluded because the reported vessel size was 10 times larger than those in other studies. Fuh *et al.*^36^ and Teran and Nugent^37^ were also excluded because their surface plasmon resonance analysis reported too large VEGF-A binding affinities (about 30 times and 700 times larger than other studies, respectively). In Teran and Nugent, especially, the rationale for this discrepancy was not reasonable. They claimed two possibilities as the reason for this discrepancy: (1) experimental settings different from those in the cell-based binding assay, and (2) the properties of the Fc-receptor chimeras used for surface plasmon resonance. However, we could not find such a noticeable difference in binding affinities across other studies using surface plasmon resonance and cell-based assays (please refer to the supplementary tables). Also, studies using surface plasmon resonance (e.g., Mamer *et al.*^38^and Papadopoulos *et al.*^39^) used Fc-receptors from the same source (R&D systems) as Teran and Nugent and yielded binding affinities similar to those reported in other studies. The discrepancy might be caused by the use of fibroblast growth factor receptors for nonspecific binding.
3. *Non-murine geometrical data (1 study)*: Belligoli *et al.*^40^ was excluded because they reported human capillary basement membrane thickness and we focused on murine tissues.
4. *Lack of tissue data (2 studies)*: Fraselle-Jacobs *et al*.^41^, which measured capillary basement membrane thickness in adipose tissue, was excluded from the analysis because we found only one paper that provided data for capillary basement membrane thickness in adipose tissue. Smith *et al.*^42^, which measured capillary basement membrane thickness in the inner ear, was excluded for the same reason.

**Figure 2.**
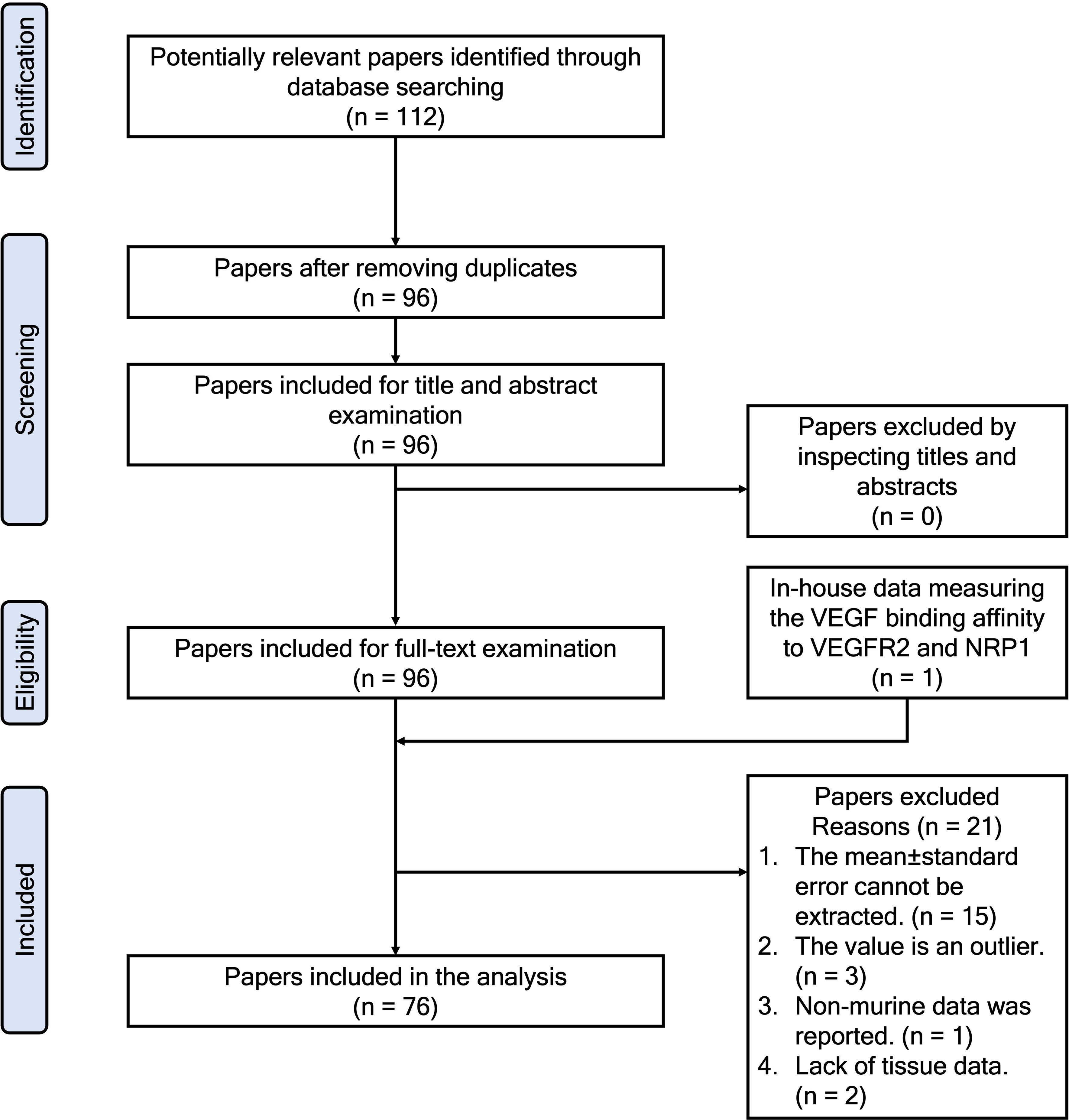
Flow chart of the analysis.

### Adipocyte diameter is larger in the adipose tissue of diet-induced obese mice than in lean mice

After selecting papers on the basis of eligibility criteria and extracting data, we first investigated adipocyte size in the most well-studied adipose tissue, the gonadal adipose tissue of lean and diet-induced obese mice. We analyzed this data because of the exceptionally large occupation of intracellular space in adipose tissue compared with tumors. Indeed, while tumor cells occupy about 40% of tumor^13–15^ volume, the adipocytes’ volume is about 90% of the total adipose tissue volume^11^ (Figure 1A). The large diameter of adipocytes contributes to their large volume percentage and the smaller interstitial space available for VEGF. This is a distinctive feature of adipose tissue compared with tumors. We analyzed 22 measurements from 10 studies (Table S1 and Figure 3). The main result is that adipocytes of obese mice were significantly larger than those of lean mice: the adipocytes of diet-induced obese mice were about 78% larger than those of lean mice (Table 1; 71 ± 5.3 µm vs. 40 ± 4.3 µm, p < 0.001).

**Figure 3.**
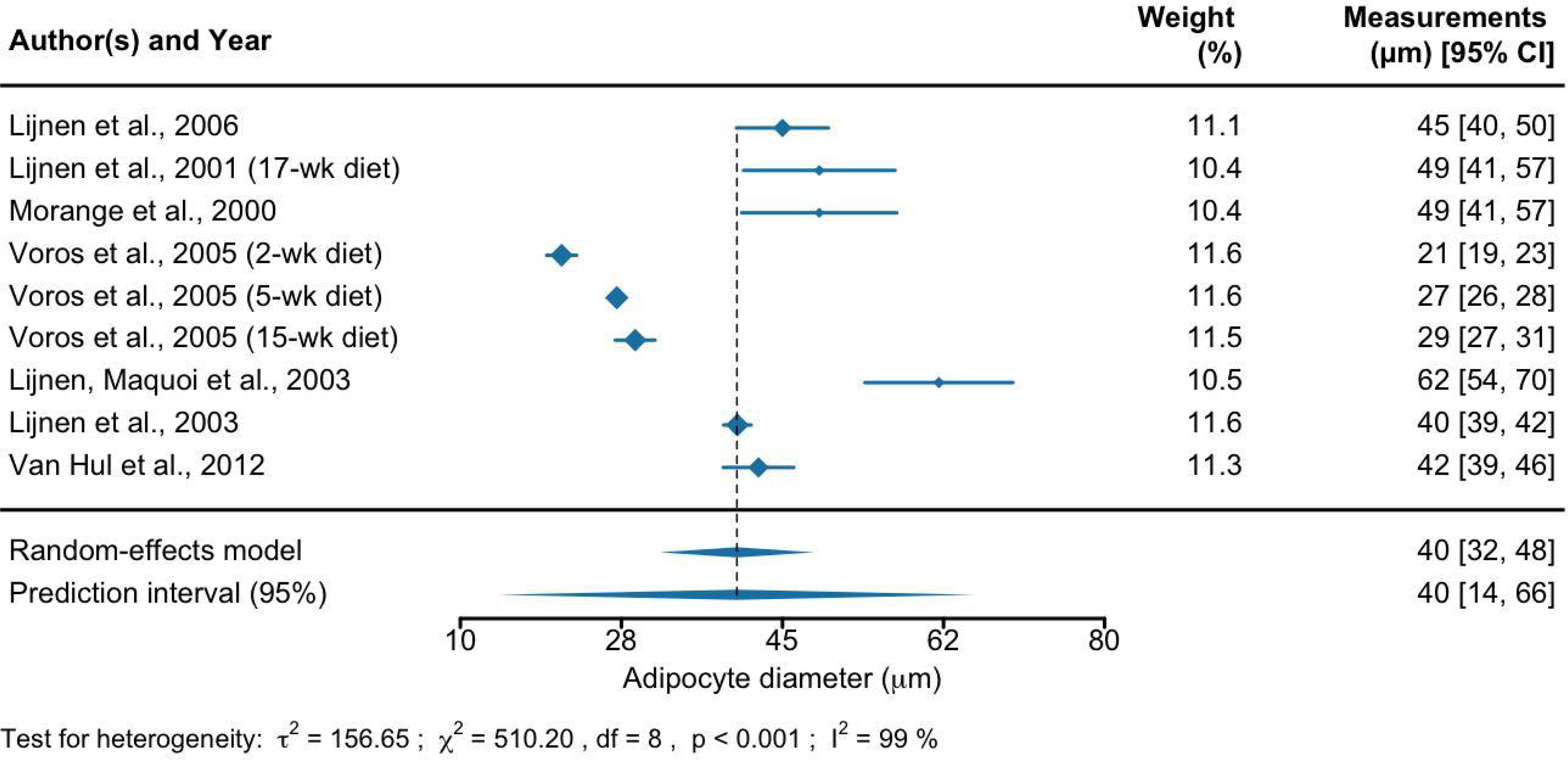
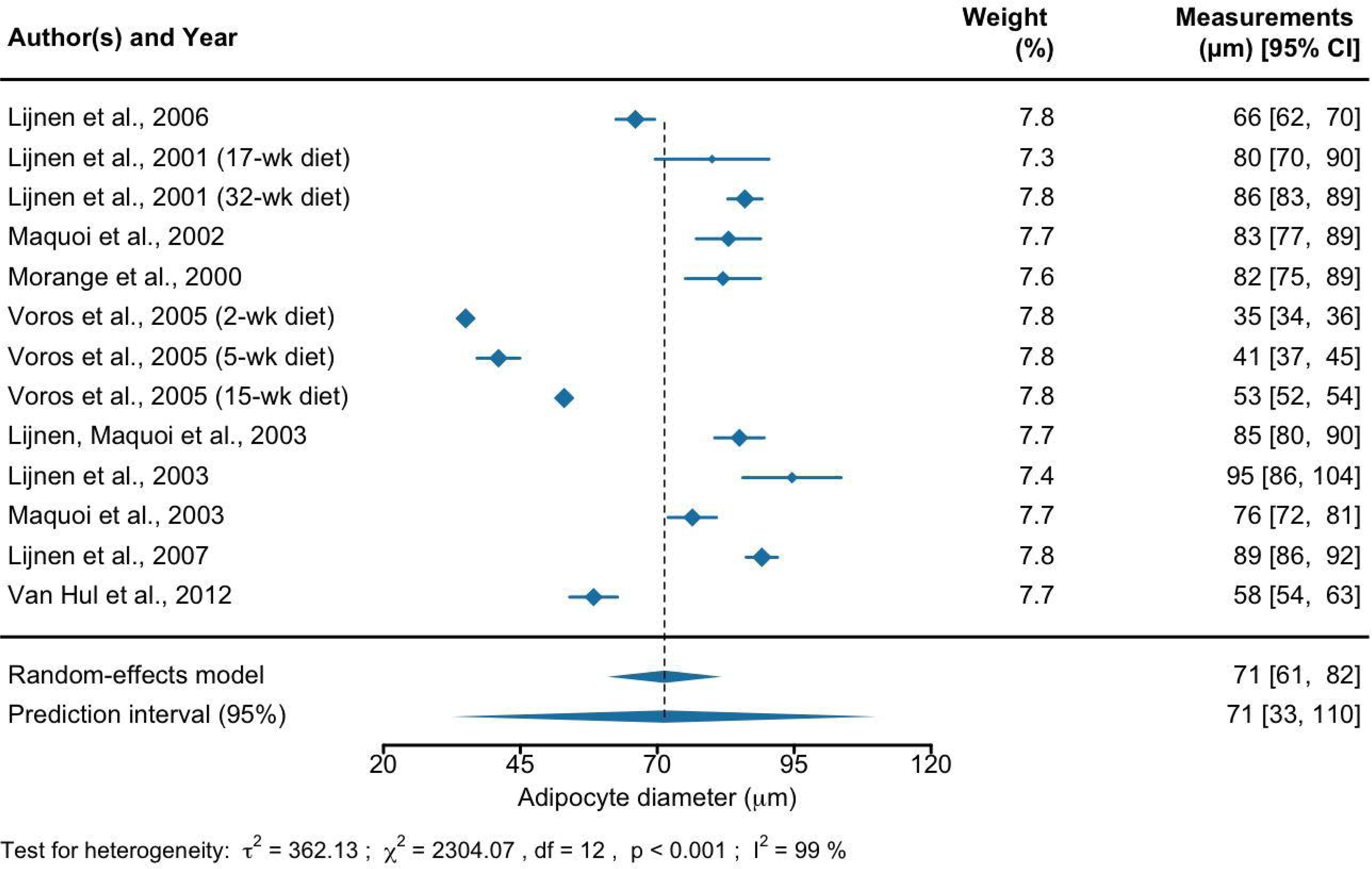
Adipocyte diameter (µm) in gonadal adipose tissue in lean and diet-induced obese mice. Forest plots of the analysis of adipocyte diameter (µm) measured in adipose tissue of lean mice (A) and obese mice (B) across studies are shown. The first, third, and last columns represent the list of references, their weights in the analysis, and measurements with a 95% confidence interval, respectively. The size of the blue diamond for each study is the visualization of its statistical weight in the analysis. The two diamonds below the table and the black dashed line show the combined measurements across studies. The upper diamond represents the mean and 95% confidence interval, and the lower diamond represents the mean and 95% prediction interval. The results of the heterogeneity test are represented by the estimate of between-studies variance (*τ*^2^), Cochran’s *Q*-test statistic for heterogeneity (*χ*^2^), degrees of freedom (df), p-value (p), and proportion of heterogeneity-induced variability among the total variability (I^2^). In the scatter plot (C), the error bar shows weighted mean ± standard error of each group. A one-tailed Welch’s t-test was used for the comparison. Abbreviation: week (wk).

**Table 1.**
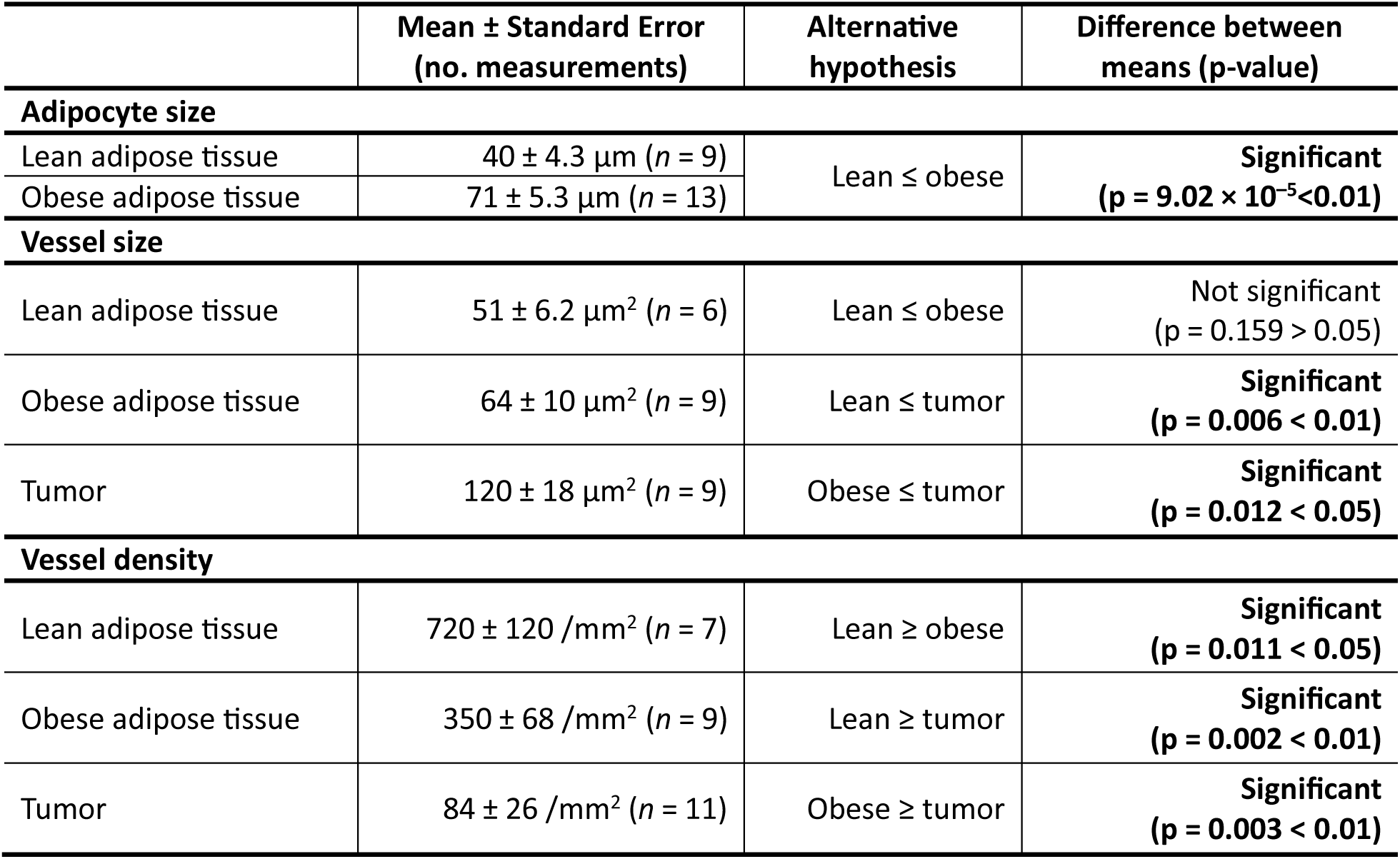
Analysis of adipocyte size, vessel size, and vessel densities in adipose tissues and tumors.

### Smaller vessel size and higher vessel density in adipose tissue compared with tumors

Adipose tissue is known to be highly vascularized^1^. As vessel size and vessel density are useful measures of vascular morphology, we compared vessel size and vessel density between obese and lean mice (Table 1, Figure 4, Tables S2, S4, Figures S1, S2, S4, and S5). We found that obese mice, in comparison with lean mice, had half the vessel density but similar vessel size (vessel density: 350 ± 68 /mm^2^ vs. 720 ± 120 /mm^2^, p = 0.011 < 0.05; vessel size: 64 ± 10 µm^2^ vs. 51 ± 6.2 µm^2^, p = 0.159 > 0.05).

**Figure 4.**
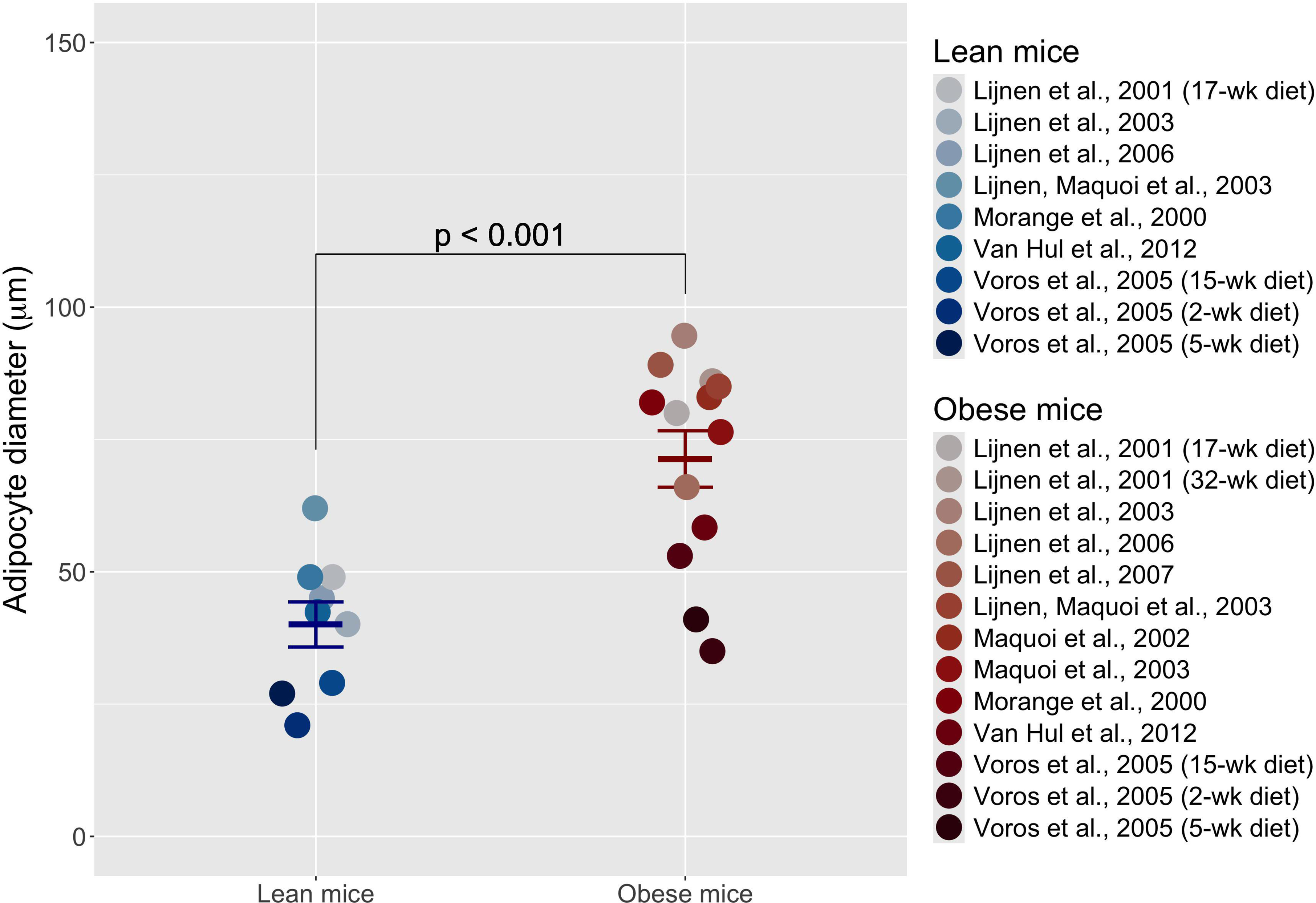
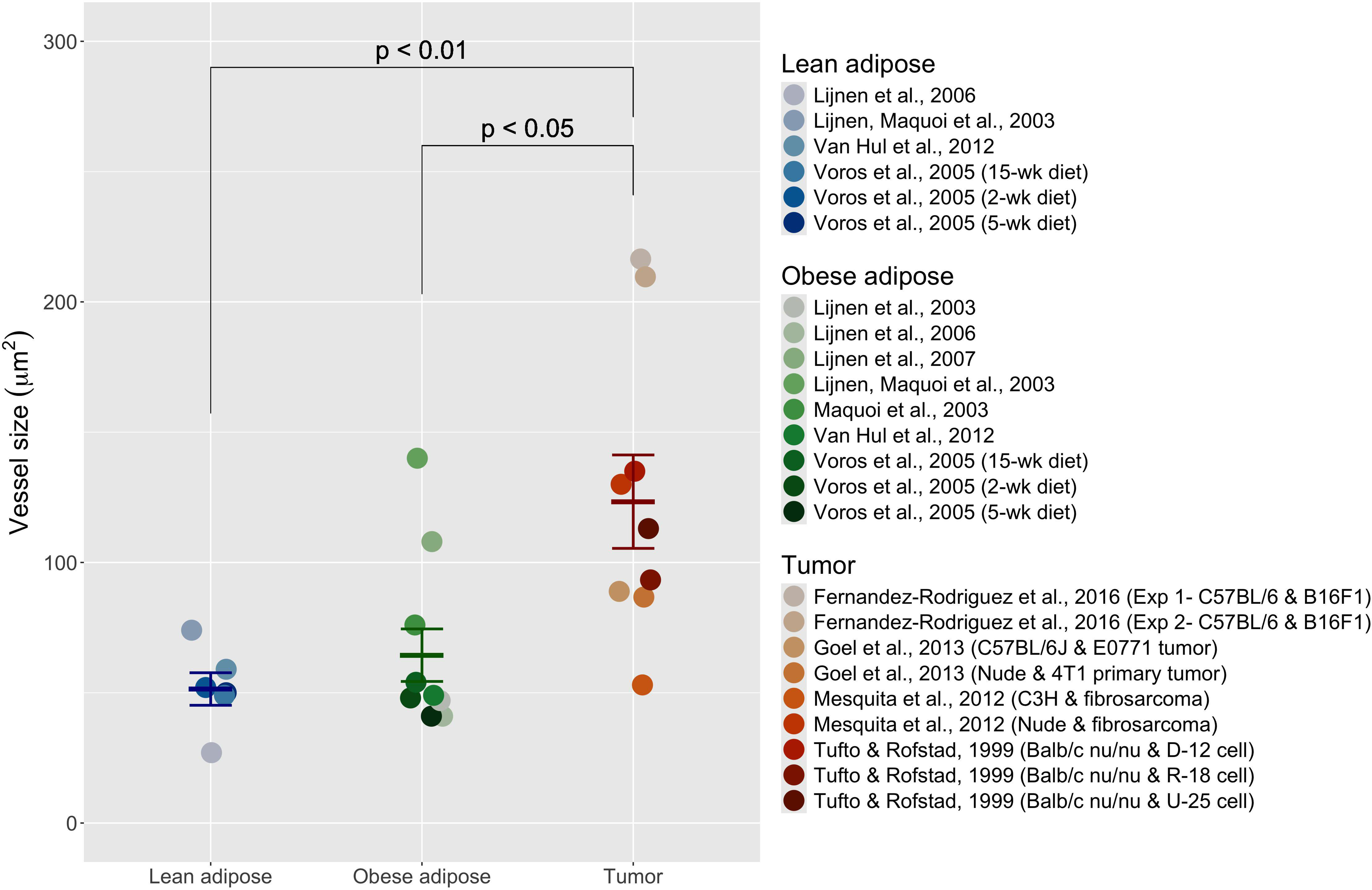
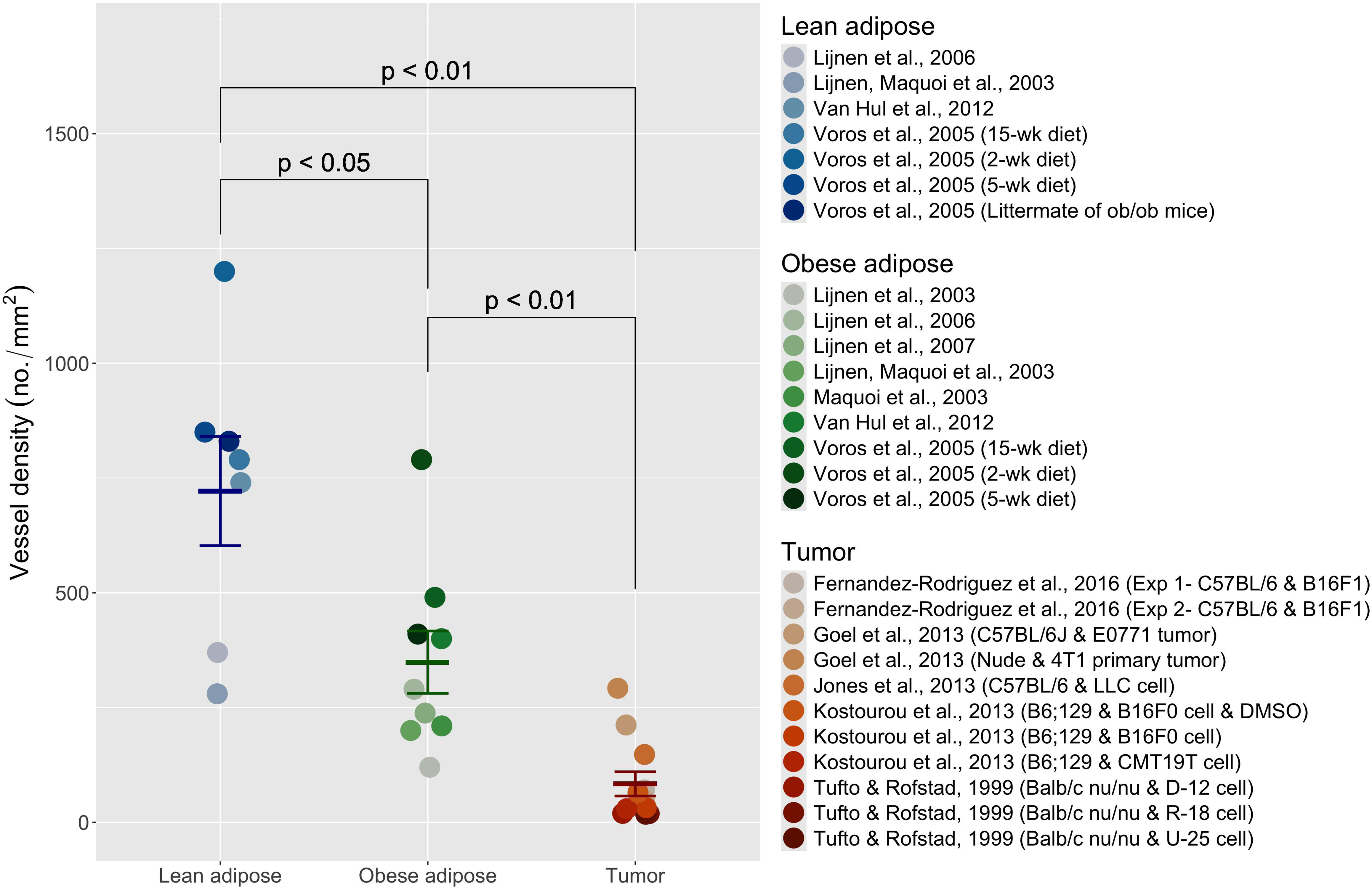
Characteristics of vessels in gonadal adipose tissue in lean and diet-induced obese mice and mouse tumors. The vessel size (A) and the vessel density (B) in adipose tissues and tumors are plotted by study. The error bar represents a weighted mean ± standard error of each group. A one-tailed Welch’s t-test (p-value < 0.05) with multiple testing correction was used to compare two tissues. The reference names in tumor data include mouse strain and tumor lines. Abbreviation: week (wk), experiment (exp), and dimethyl sulfoxide (DMSO).

Tumor tissue is also known for its high vascularity^43^, and tumor vessels in lean mice were two times larger than in lean and obese adipose tissue (Figures 4A and S3, and Table S3; lean adipose vs. tumor: 51 ± 6.2 µm^2^ vs. 120 ± 18 µm^2^, p = 0.006 < 0.01; obese adipose vs. tumor: 64 ± 10 µm^2^ vs. 120 ± 18 µm^2^, p = 0.012 < 0.05). However, the tumor was four and nine times less vascularized than were lean and obese adipose tissue, respectively (Figures 4B and S6, and Table S5; lean adipose vs. tumor: 720 ± 120 /mm^2^ vs. 84 ± 26/mm^2^, p = 0.002 < 0.01; obese adipose vs. tumor: 350 ± 68 /mm^2^ vs. 84 ± 26 /mm^2^, p = 0.003 < 0.01). Overall, our data show that obesity reduces vascular density but not vessel size in adipose tissue and that adipose tissue has higher vascular density but smaller vessel size than tumors.

### Effect of obesity on capillary basement membrane thickness

The capillary basement membrane is a component of the extracellular matrix in tissue. Since the thicker capillary basement membrane occupies a larger volume fraction in adipose tissue, its thickness affects the interstitial space volume in the tissue. Additionally, it is known that capillary basement membrane thickening is associated with diabetes, and diabetes is one of the common comorbidities of obesity. Thus, to determine the relationship between capillary basement membrane thickness and obesity, we analyzed 34 studies that measured capillary basement membrane thickness in lean and fat mice and rats (Table 2, Figure 5, Table S6, and Figures S7–S8). The measurements from the retina, muscle, and heart were examined in both categories of mice and rats since measurements from other tissues (e.g., brain and kidney) were not found for obese mice and rats. Interestingly, capillary basement membrane thickness in tissues of obese mice and rats was similar to that of lean mice and rats, with no significant difference (obese vs. lean: 104 ± 14 nm vs. 94 ± 5 nm, p = 0.536 > 0.05). Our result indicates that obesity does not affect capillary basement membrane thickness.

**Figure 5.**
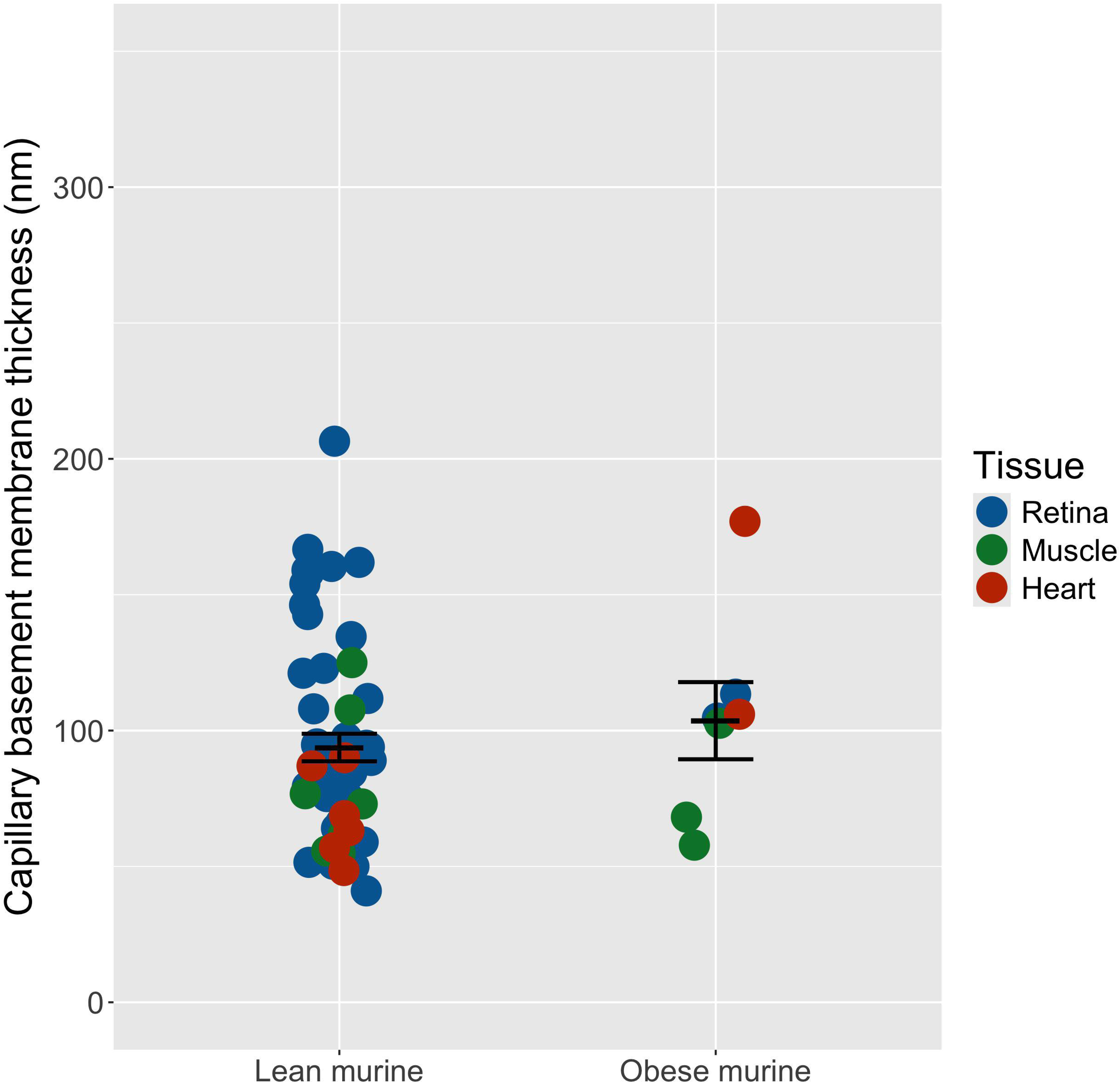
Capillary basement membrane thickness (nm) in lean and obese mouse and rat tissues. The capillary basement membranes in retina, muscle, and heart were compared between lean and obese mice and rats. A two-tailed Welch’s t-test (p-value < 0.05) was used to compare CBM thicknesses.

**Table 2.**
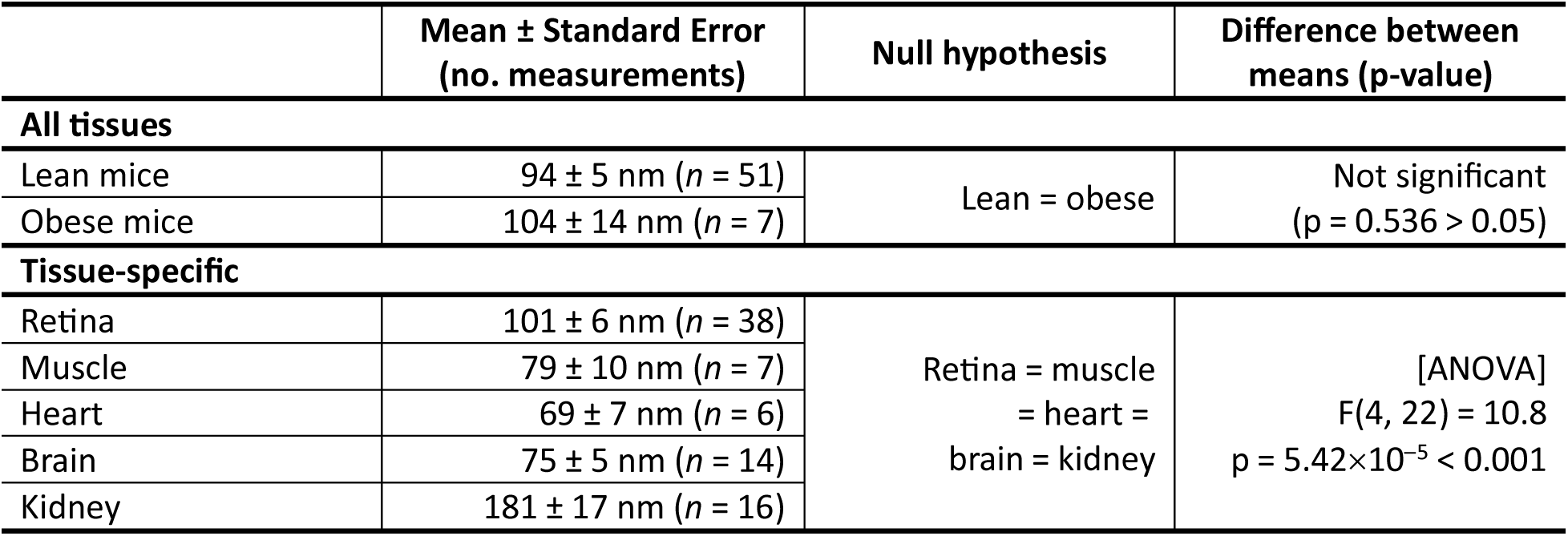
Capillary basement membrane thickness in mouse and rat tissues.

In order to examine if capillary basement membrane thickness varies across tissues, we examined capillary basement membrane measurements from multiple tissues: retina, muscle, heart, brain, and kidney (Table 2, Table S6, Figures S9–S14). These tissues were chosen because they are the most well-studied tissues in the field in regard to capillary basement membrane thickness. The obese mouse and rat data were excluded from the retina, muscle, and heart datasets for fair comparison, since they did not include brain and kidney measurements. The test for homogeneity of variance showed significantly different variances across tissues (p = 1.30 × 10^-5^ <0.001). From the following Welch’s ANOVA and Dunnett’s T3 test, the retina had similar capillary basement membrane thickness as muscle (101 ± 6 nm vs. 79 ± 10 nm, p = 0.555 > 0.05; Table S10), but the retina had significantly different thickness compared with heart and brain (retina: 101 ± 6 nm vs. heart: 69 ± 7 nm, p = 0.031; retina vs. brain: 101 ± 6 nm vs. 75 ± 5 nm, p = 0.019). Capillary basement membrane thickness in muscle was similar to that in heart and brain (muscle vs. heart: 79 ± 10 nm vs. 69 ± 7 nm, p = 0.990 > 0.05; muscle vs. brain: 79 ± 10 nm vs. 75 ± 5 nm, p = 1.000 > 0.05). We did not analyze this in adipose tissue, because of the single datum (109 ± 11 nm from Fraselle-Jacobs *et al.*^41^). Capillary basement membrane thickness was greatest in the kidney (181 ± 17 nm), and it was about 2–3 times thicker than in other tissues (retina vs. kidney: p < 0.01; muscle vs. kidney: p < 0.001; heart vs. kidney: p < 0.001; brain vs. kidney: p < 0.001). Our result suggests that capillary basement membrane thickness varies across tissues.

### Binding data for VEGF-A:VEGFR2 and VEGF-A:NRP1 measured by surface plasmon resonance

We measured the binding rates and affinities for VEGF-A with NRP1 (Figure S15A), as kinetics and affinity studies were lacking. The VEGF-A:VEGFR2 binding rates were also measured as a positive control. The VEGF-A:NRP1 binding affinity (K_d_) was measured as 6.36 ± 1.07 nM, with an association rate (k_on_) of 7.96 ± 2.15 × 10^5^ M^−1^ s^−1^ and dissociation rate (k_off_) of 1.56 ± 0.55 × 10^-3^ s^-1^. The kinetics rate for VEGF-A and VEGFR2 was determined with binding affinity K_d_ = 520 ± 250 pM, which is in a pM range similar to those reported in other studies (Figure S15B)^44–46^. The association rate (k_on_) for VEGF-A:VEGFR2 was calculated to be 6.24 ± 0.46 × 10^5^ M^-1^s^-1^, with the dissociation rate (k_off_) of 3.18 ± 1.98 × 10^-4^ s^-1^. These measurements were included in our analysis. We measured a lower binding affinity and slower dissociation rate in VEGF-A’s binding to NRP1 in comparison with its binding to VEGFR2.

### Stronger binding of VEGF-A to VEGFR1 than to VEGFR2 and NRP1

VEGF receptors have distinct functions in angiogenesis; thus, comparing their binding affinities offers receptor-level insights into differential VEGFR signaling in obesity. We analyzed 21 studies that measured VEGF-A binding affinity to VEGFR1, VEGFR2, and the co-receptor NRP1, using either radioligand assays or surface plasmon resonance (Table 3, Figure 6A, Tables S7–S9, Figures S16–S18). As expected, the binding affinity of VEGF-A to VEGFR1 was significantly stronger than to VEGFR2, being about six times stronger (34 ± 12 pM vs. 210 ± 60 pM, p = 0.017 < 0.05). On the other hand, the binding affinity of VEGF-A to NRP1 was not significantly weaker than the VEGF-A:VEGFR1 affinity, although it was about 24 times weaker (34 ± 12 pM vs. 820 ± 350 pM, p = 0.065 > 0.05). Similarly, VEGF-A binding affinity to NRP1 was not significantly weaker than to VEGFR2, although it was four times weaker (210 ± 60 pM vs. 820 ± 350 pM, p = 0.088 > 0.05). Overall, our study shows the strongest binding affinity of VEGF-A to VEGFR1 and its weakest binding affinity to NRP1.

**Figure 6.**
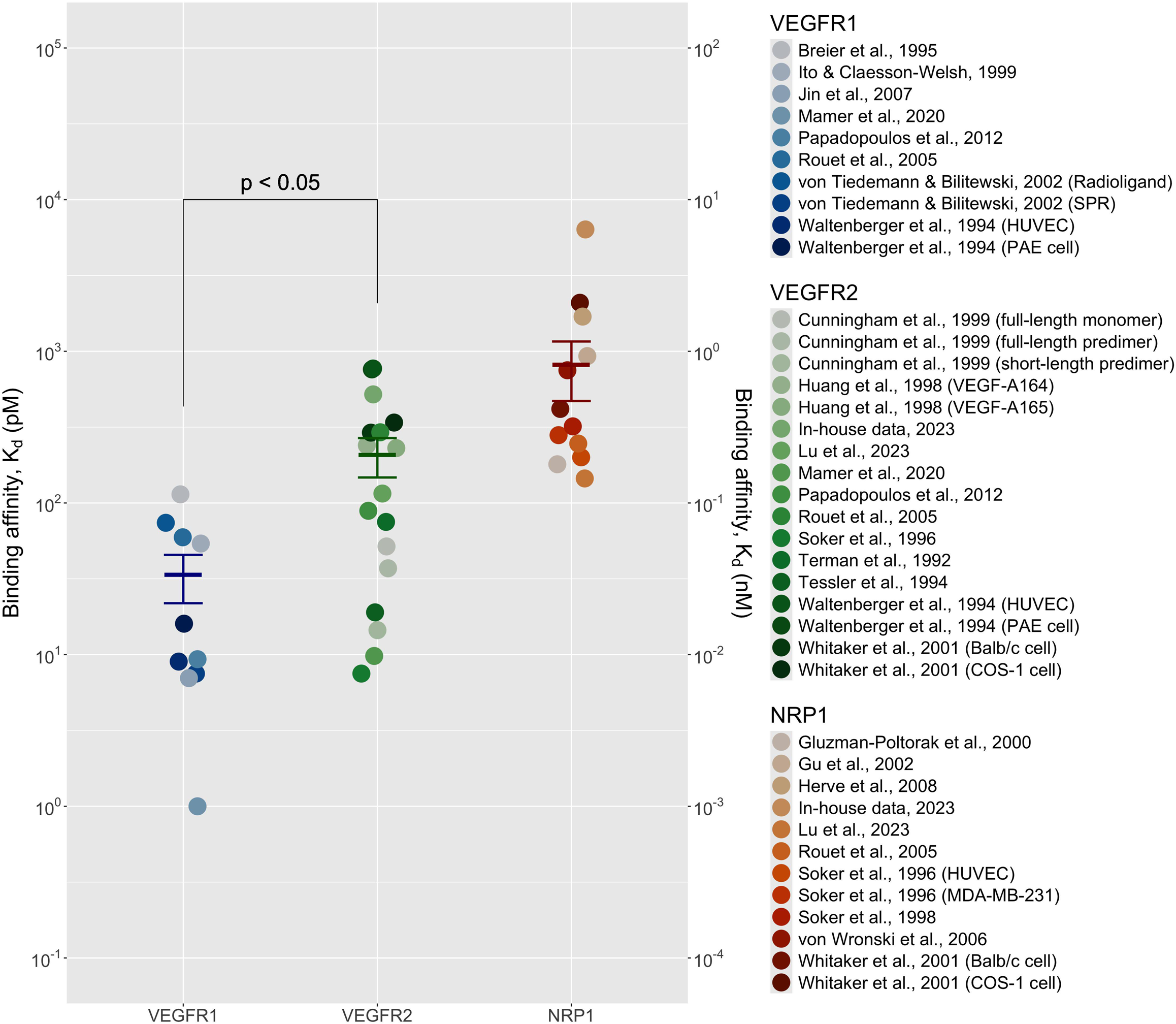
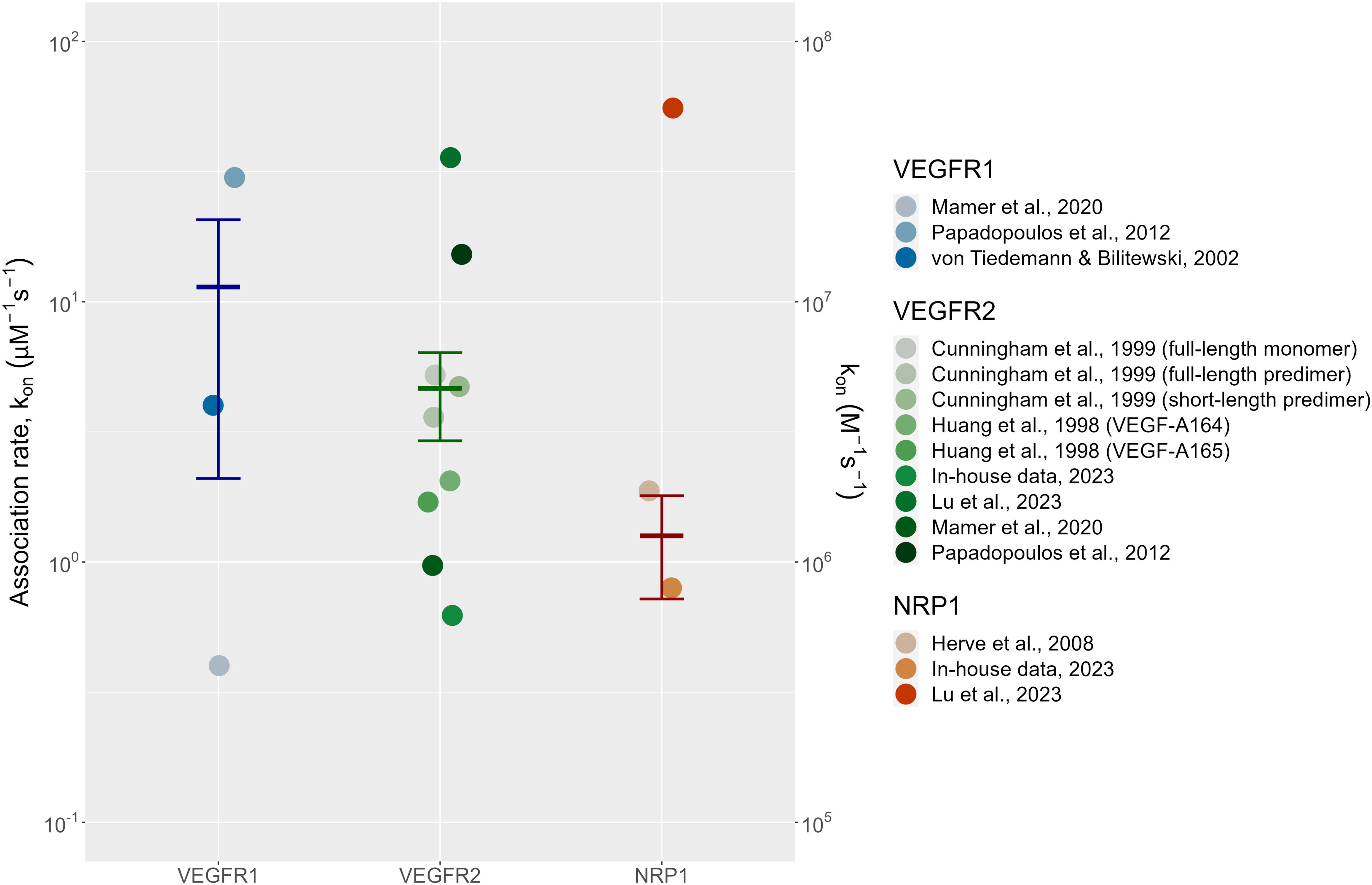
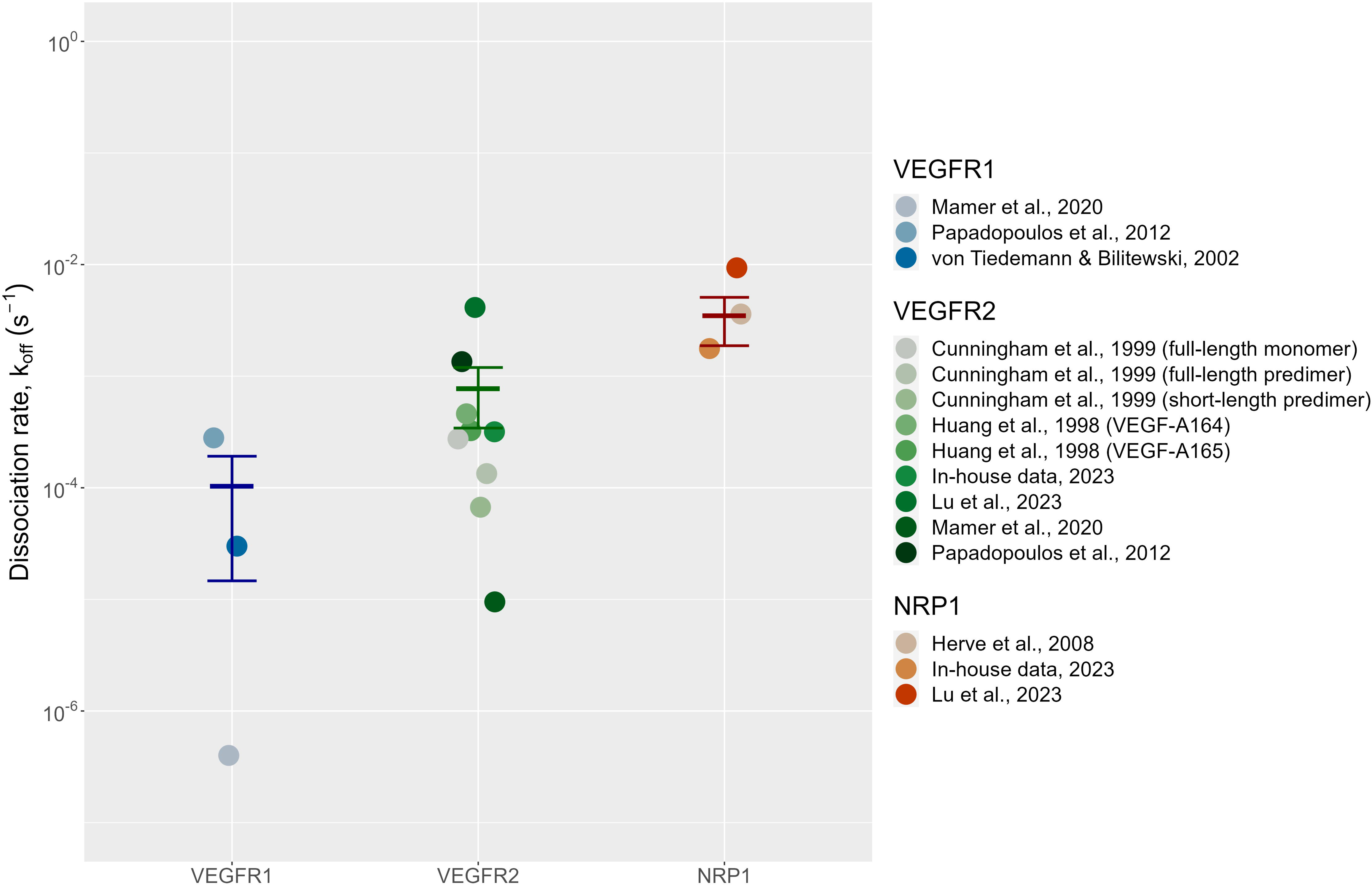
Binding rates of VEGF to VEGFR1, VEGFR2, and NRP1. The respective binding affinities (A; K_d_), association rate constants (B; k_on_, M^−1^ s^−1^), and dissociation rate constants (C; k_off_, s^−1^) of VEGF for VEGFR1, VEGFR2, and NRP1 are plotted. The error bar represents the weighted mean ± standard error of the rate for each receptor. A one-tailed Welch’s t-test with multiple testing corrections (p-value < 0.05) was used to compare a pair of groups. Abbreviation: surface plasmon resonance (SPR), human umbilical vein endothelial cell (HUVEC), and porcine aortic endothelial (PAE).

**Table 3.**
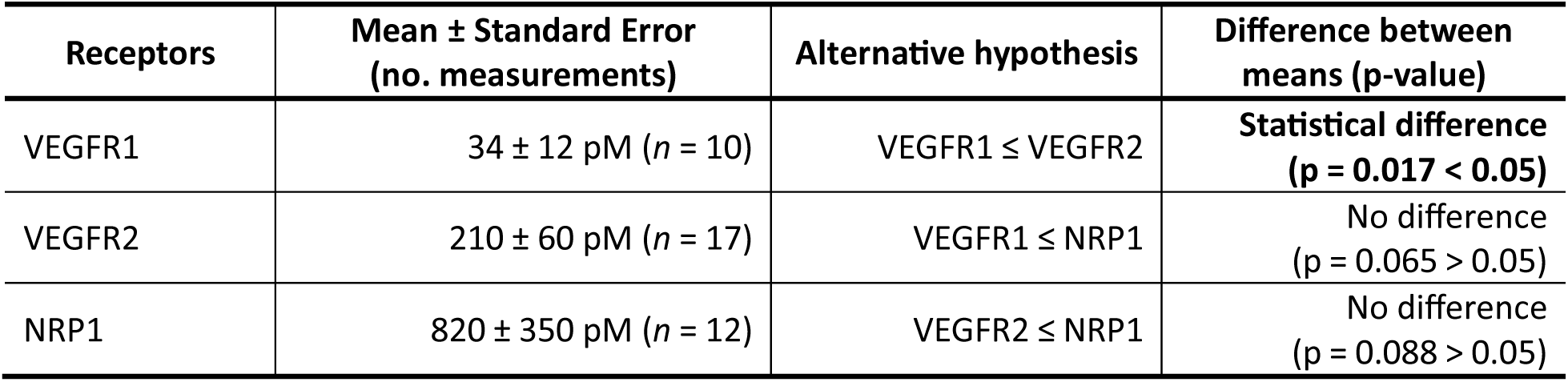
Analytic K_d_ values for VEGF-A binding to its receptors.

Association (k_on_) and dissociation (k_off_) rate constants allow us to predict not only the equilibrium state of a ligand-receptor interaction, but also how fast the system responds to changes in the concentration of the ligand or to another competitor. We analyzed data for VEGFR1, VEGFR2, and NRP1 binding kinetics with VEGF-A. The association rates of VEGF-A and these receptors were not significantly different, although the association rate of VEGFR1 was larger than that of VEGFR2, as expected (Table 4, Figure 6B, Table S7–S9, Figure S19–S21; 11 ± 9.3 × 10^6^ M^-1^s^-1^ vs. 4.6 ± 1.7 × 10^6^ M^-1^s^-1^, p = 0.274 > 0.05). The association rate of NRP1 (1.3 ± 0.54 × 10^6^ M^-1^s^-1^) was four to eight times smaller than those of VEGFR1 and VEGFR2 (p = 0.274 > 0.05 and p = 0.238 > 0.05, respectively). The dissociation rate constants for all receptors were also similar, although the mean for VEGFR1 was eight times smaller than the mean for VEGFR2 (Table 4, Figure 6C, Table S7–S9, Figure S22–S24; 1.0 ± 0.89 × 10^-4^ s^-1^ vs. 7.7 ± 4.3 × 10^-4^ s^-1^), and that of NRP1 was larger by an order of magnitude (3.5 ± 1.6 × 10^-3^ s^-1^). Overall, our data suggest that association and dissociation rates of VEGF-A with these receptors do not differ significantly, while its binding affinities for VEGFR1 and VEGFR2 do differ significantly.

**Table 4.**
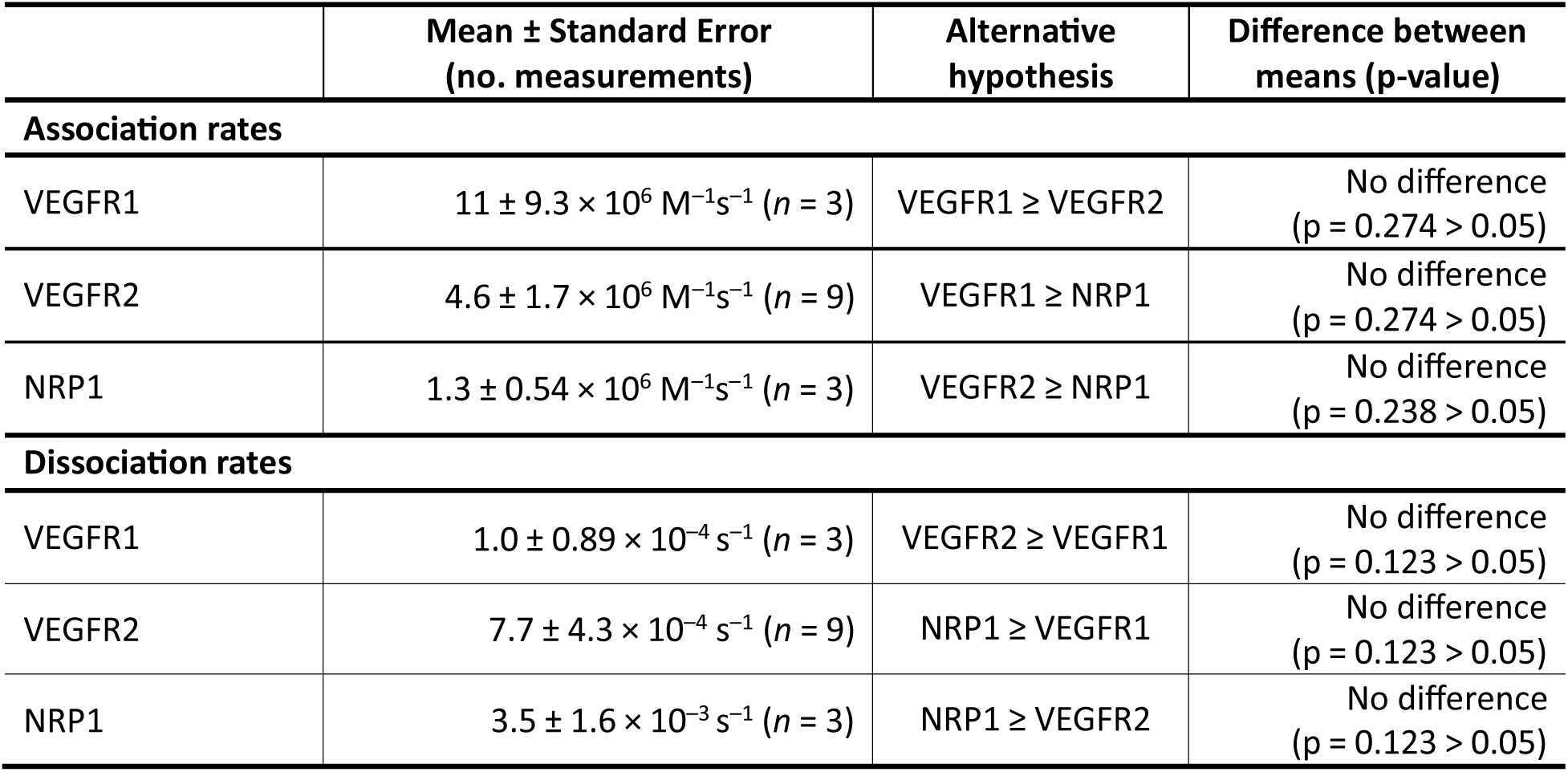
Analytic association and dissociation rate values of VEGF-A binding to VEGFR1, VEGFR2, and NRP1.

We also examined the binding affinities of VEGF-B and VEGF-C for VEGF receptors (Table 5). VEGF-B binds to VEGFR1 and NRP1, while VEGF-C binds to VEGFR2 and VEGFR3. VEGF-B includes two isoforms, VEGF-B167 and VEGF-B186, with VEGF-B167 predominantly expressed in most normal tissues^47^. VEGF-B186 must be proteolytically processed to bind to NRP1, while VEGF-B167 binds to NRP1 without cleavage^48^. From the data mining, we found that VEGF-B binds to VEGFR1 and the b1 domain of NRP1 with binding affinities of 114 pM and 36 µM, respectively. VEGF-B peptides bind to the b1 domain of NRP1 with binding affinities of 0.39–9.55 µM. Most studies show that the mature VEGF-C and its mutants bind to VEGFR2 and VEGFR3 with binding affinities in the nM range, while Joukov *et al*.^49^ reported a pM range of K_d_ for VEGF-C:VEGFR2 and VEGFR3. These different binding affinities are possibly due to different measurement methods or cells used to produce recombinant VEGF-C. Due to the lack of data, we did not analyze them using a random-effects model.

**Table 5.**
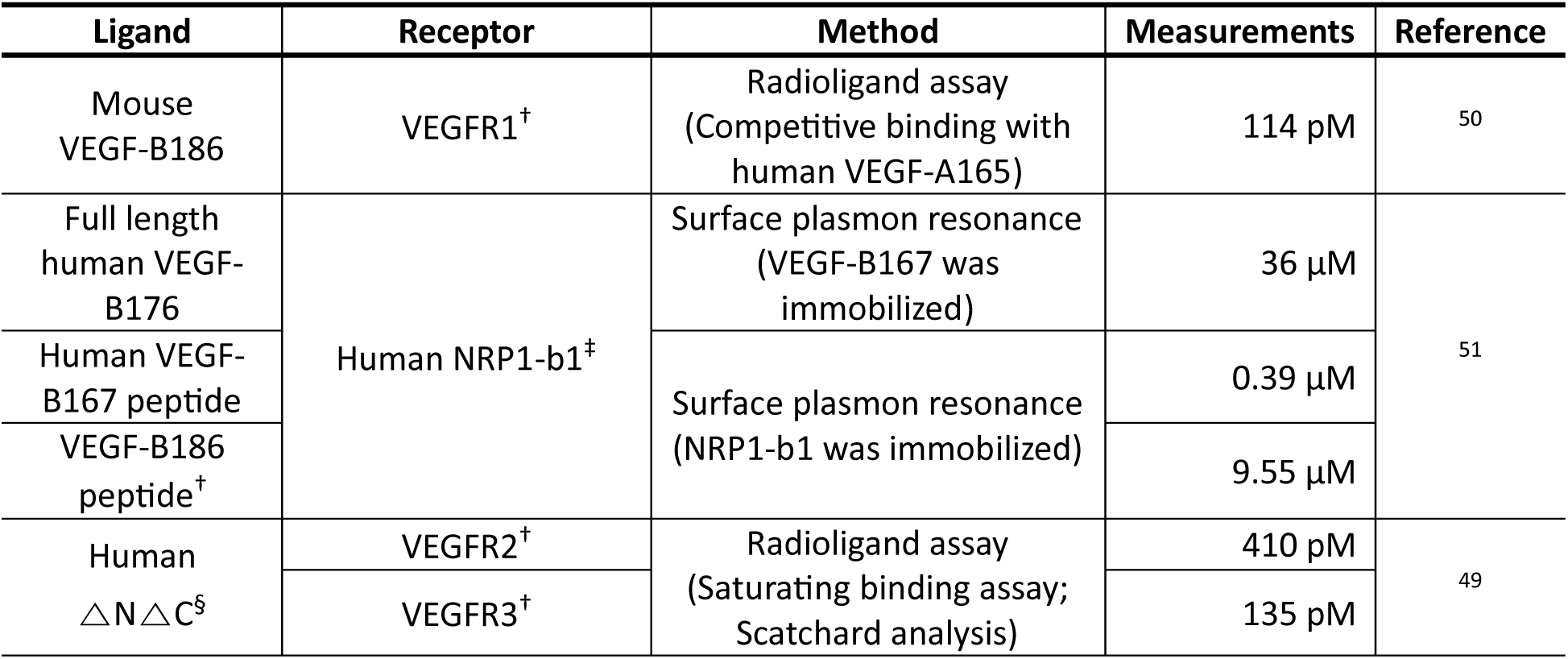
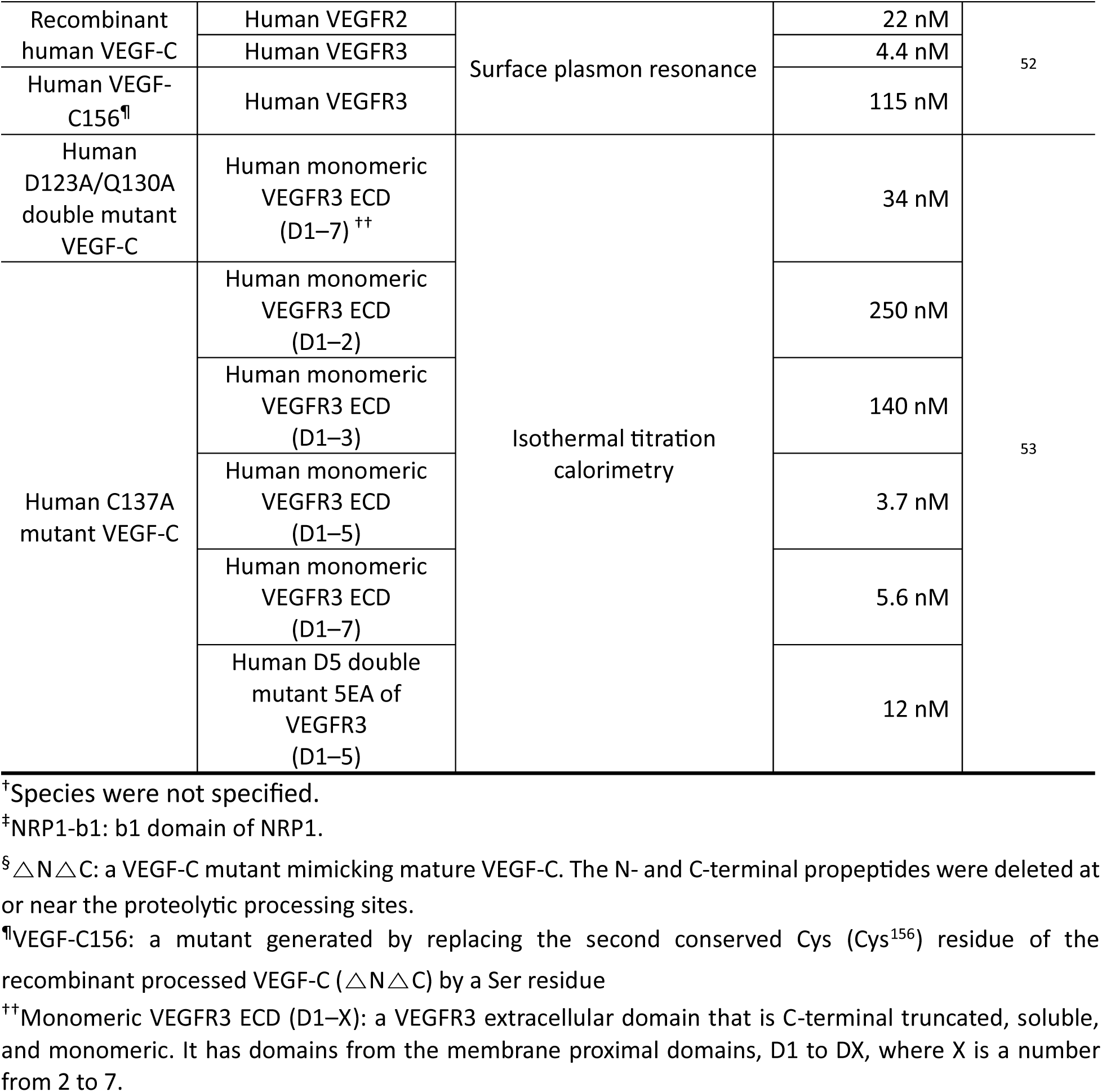
Binding affinities of VEGF-B and VEGF-C to VEGFRs and NRP1.

### Significant difference in binding affinities between radioligand and SPR assays, except for VEGF-A:NRP1

The binding affinity is a specific type of equilibrium constant, calculated by dividing the dissociation rate “constant” by the association “constant”. Thus, theoretically, the value should be consistent under similar observation conditions. Nevertheless, different techniques to detect ligand–receptor binding may yield variations in binding kinetic measurements due to different experimental settings. In order to assess variability in VEGF-A binding affinity measurements, we analyzed data from radioligand assays and SPR. The analysis of 21 studies showed that the VEGF-A binding affinity for VEGFR1 measured by radioligand assay was about eight times weaker than that measured by SPR: 46 ± 15 pM vs. 5.9 ± 2.6 pM, respectively (Table 6, Figure S25–S27). Also, the binding affinity of VEGF-A to VEGFR2 measured by radioligand assay was about six times weaker than that measured by SPR (Figure S28 and S29; radioligand assay vs. SPR; 310 ± 110 pM vs. 51 ± 15 pM) while radioligand assay showed seven-times stronger binding affinities for

**Table 6.**
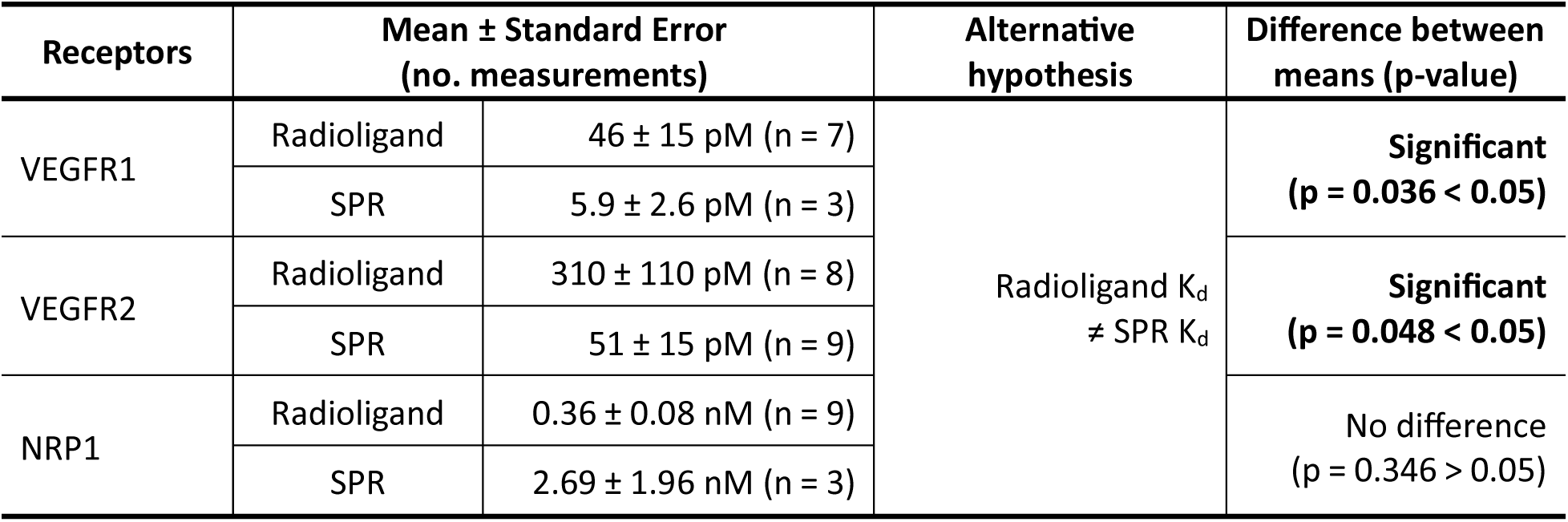
Analytic K_d_ values for VEGF-A binding to its receptors measured by radioligand and SPR (surface plasmon resonance) assays.

NRP1 (Figure S30 and S31; 0.36 ± 0.08 nM vs. 2.69 ± 1.96 nM). We applied Welch’s t-test to compare K_d_ values measured by radioligand assay and SPR for each receptor. All results of t-tests for VEGF receptors except NRP1 were significant (p = 0.036 < 0.05 for VEGFR1; p = 0.048 < 0.05 for VEGFR2; p = 0.346 > 0.05 for NRP1). Our results indicate that radioligand assay and SPR report significantly different VEGFR binding affinities.

## Discussion

The prevalence of obesity in the United States reached 42% in 2021^54^, and obesity causes severe cardiovascular diseases. Adipose tissue expansion in obesity requires angiogenesis, but how to control angiogenesis in obesity is still unknown. Systems biology can provide new insight into this problem by incorporating biological data and computational models. Several studies have constructed computational models to understand pathological angiogenesis in cancer or peripheral artery diseases^13,55^. However, applying the same models to adipose tissue requires adaptation to its unique microenvironment, specifically by incorporating adipose tissue-specific data. Thus, prior to developing obesity models, it is important to identify and analyze data affecting angiogenic signaling in the targeted tissue. The work in this paper yielded the following five key findings: (1) obese mice have 78% larger adipocytes than lean mice; (2) obesity reduces vessel density but does not affect vessel size, and adipose tissue has smaller vessel size but higher vessel density than tumors; (3) obesity does not affect capillary basement membrane thickness, as opposed to what has been reported in diabetes; (4) by standardizing the binding rates, we confirmed that VEGF-A’s strongest binding is to VEGFR1 rather than to VEGFR2 or NRP1; and (5) the binding affinities measured by radioligand assay and SPR are significantly different. From these key findings, our study will enable the development of adipose tissue-specific computational models.

### Obesity increases adipocyte size and reduces vessel density but not vessel size

Our analysis determined that in obese mice, adipocyte diameter is increased by 78% and vessel density is reduced by 51%, consistent with findings from a previous meta-analysis of human adipose tissue^56^. However, vessel size may not be affected by obesity, as suggested by our analysis. While we did not identify studies statistically comparing vessel size between lean and obese human subjects, representative histological images with stained vessels appear to show that the vessels in their adipose tissues are of similar size^57^. In obesogenic conditions, adipocytes undergo hypertrophy (cell enlargement), which reduces vascular density and creates hypoxic conditions within the tissue, making these processes crucial for inclusion in a computational model of obesity. Overall, our study confirms the association of obesity with adipocyte size and vessel density and suggests its non-association with vessel size.

### Contribution of our study to the successful development of adipose-tissue models; difference between adipose tissue and tumor vasculature

Because both tumors and obesity are affected by hypoxia, one may hypothesize that vessels in the two conditions may be similar. Hence, our work aimed to identify (1) any similarities that would enable computational tumor models to be applied to obesity, or (2) any differences that would justify novel obesity-specific computational models. To address this, we examined the differences in vascular morphologies between adipose tissue and tumors, demonstrating that adipose tissues exhibit a higher vessel density but a smaller vessel size than tumors. Specifically, the vessel size in adipose tissue is approximately half that in tumors, while vessel density is four to nine times higher. Consequently, the surface area of vessels per unit volume in adipose tissue may be 3–6 times greater than in tumors.

These differences that we observed between tumors and obesity may be understood through their differing mechanisms of hypoxia and angiogenesis. In obesity, hypoxia is induced as vessel densities are reduced by the enlargement of adipocytes, while in tumors hypoxia results from poor oxygen diffusion from abnormal vasculature^58,59^. Tumors exhibit chaotic vessel formation due to their continuous pro-angiogenic state and rapid neovascularization. This rapid angiogenesis limits the development of mature vessel structures and results in larger, irregular vessels often concentrated at the periphery rather than throughout the tissue^60,61^. In contrast, adipose tissue has unique features. The stromal vascular fraction in the adipose tissue supports a well-organized, hierarchical vasculature where each adipocyte is adjacent to at least one vessel, creating an evenly distributed network across the tissue^62,63^.

If a tumor model is adopted for obesity, this significant difference would lead to incorrect predictions of VEGF-A distributions in adipose tissue (e.g., concentrations of free VEGF-A, interstitial matrix-bound VEGF-A, and receptor-bound VEGF-A) and potentially affect predictions of anti-VEGF drug efficacy. Instead, the development of adipose-specific models, with parameters defined here, would enable researchers to identify the most effective obesity treatment.

### Why does obesity not affect capillary basement membrane thickness?

We sought to understand whether obesity alters capillary basement membrane (CBM) thickness in adipose tissue, but could not because there were no data available. We instead examined three organs from obese mice and rats—retina, muscle, and heart—and compared them with their correlates in lean mice and rats. This analysis showed that CBM thickness is not altered by obesity in non-adipose tissue. Thus, in computational modeling, the CBM thickness could be assumed to have the same value in lean and obese conditions. Experimental studies should clarify whether obese and lean adipose tissue also have similar CBM.

The insensitivity of CBM thickness to obesity was unexpected, given that 80% of the studies we analyzed used prediabetic and diabetic mice and rats (40% for each group; only 20% used obese, non-diabetic mice) and that diabetes, a common comorbidity of obesity, has been associated with CBM thickening in diabetic models, particularly in the retina, muscles, brain, and kidneys^64–69^. In fact, CBM thickening has been recognized as a hallmark of diabetic retinopathy, nephropathy, and cardiomyopathy, and in the brain, it is also associated with Alzheimer’s disease^70^. This raises the question: what factors might account for the lack of a significant effect of obesity on CBM thickness in our study? We suggest two possible explanations for our findings:

1. **Duration of diabetes**: The duration of diabetes rather than diabetes itself may determine the CBM thickness^71^. For example, 6–7-month-old Zucker diabetic fatty rats develop diabetes after 3–5 months of age^72^, suggesting that CBM thickening may not have progressed enough to reach significance. Similarly, C57BL/6J mice develop obesity and diabetes after 4 months on a diet high in fat and simple carbohydrate^73^. However, the mice in Williams *et al.* had undergone only a 4-month high-fat diet and exhibited CBM thickness comparable to that of lean mice^74^.
2. **Obese rodent strains**: The strain of obese rats included in our analysis may also explain our findings. Studies by Lash *et al.* and Dosso *et al.* on obese Zucker rats^75,76^, which are a model for prediabetes^77^, support this possibility. Considering that humans with prediabetes exhibit CBM thickness similar to that of lean, healthy humans (103 nm vs. 117 nm)^40^, it is reasonable to expect that CBM thickness will not differ significantly between the prediabetic Zucker rats and lean rats.

We sought to determine if CBM thickness varies across tissues, in order to decide whether computational models should consider this aspect of the targeted tissue. Importantly, tissues have phenotypically different capillaries depending on their function. For example, kidneys have continuous fenestrated capillaries, with continuous basement membrane and fenestrated endothelium that allows large molecules to pass across the wall^78^. On the other hand, heart and muscle have continuous basement membranes and non-fenestrated endothelium, which allow only small molecules to pass^78,79^. The liver has discontinuous capillaries with a fragmented basement membrane, allowing the movement of large molecules for liver metabolism^80^. All tissues we examined in this study have continuous CBMs, so we aimed to identify any differences in CBM thickness between these tissues of lean mice and rats. The retina and kidney have higher CBM thicknesses than other tissues, possibly due to different distributions of molecular components (collagen, nidogen, etc.), cell types, or physical factors (e.g., hydrostatic pressure)^81,82^. For example, the kidney CBM is composed of two basement membranes: one from the endothelium and another from the epithelium^83^.

### Standardized VEGF-A binding rates accelerate the development of a more feasible computational model

Our analysis confirms that VEGF-A binding to VEGFR1 is the strongest and establishes that VEGF-A binding to NRP1 is the weakest of its binding to receptors. This strong VEGF:VEGFR1 binding is attributed to its highest association rate (k_on_) and lowest dissociation rate (k_off_), whereas VEGF:NRP1 binding displayed the opposite kinetic profile. The binding affinity (K_d_) is calculated by K_d_ = k_off_/k_on_. Our standardized values of VEGF-A binding affinities to VEGFR1 and VEGFR2 are consistent with values used in previous computational studies, while VEGF-A binding affinity to NRP1 from our study was three times weaker than the previously used values^13,84,85^. On the other hand, our association and dissociation rate constants differed by two to four times and 1.3 to 3.5 times from the previously used values, respectively. This difference might have affected the simulation outcomes of the previous studies^13,84,85^ such as the proportion of VEGF-A among other ligands bound to the same VEGF receptors, because the rate constants are key factors determining how fast VEGF-A binds to its receptors in competing with other ligands. To our best knowledge, ours is the first analysis to standardize the VEGF-A binding rates, because only a few studies have gathered and analyzed VEGF-A binding data^86^. Thus, our analysis enhances the understanding of complex VEGF-A signaling mechanisms and enables the development of more feasible computational models by providing standardized binding affinities.

### Variation in measurements from different ligand-binding assays: computational modelers should be aware of this when choosing kinetic values

Our analysis revealed that measurement techniques can significantly impact binding affinity values. Specifically, binding affinities are commonly measured using two techniques: the radioligand assay and surface plasmon resonance (SPR). When comparing data from these methods, we found that radioligand assays yield higher binding affinity measurements for VEGF-A:VEGFR1 and VEGF-A:VEGFR2, while no difference was observed for VEGF-A:NRP1. Several factors may explain these notable differences. First, heparin has been shown to reduce the VEGF-A:VEGFR1 binding affinity in porcine aortic endothelial cells^87^. Specifically, K_d_ was 54 pM in the absence of heparin but increased to 77–118 pM when heparin was present. This suggests that heparan sulfate proteoglycan (HSPG) on the cell membrane likely influences the binding affinity observed in the radioligand assay, which is a cell-based assay. In contrast, SPR is a chip-based assay and may not fully replicate the membrane-associated effects. Second, the different structures of receptors used in the radioligand assay and SPR may also affect the binding affinity. While the radioligand assays that we cited in our analysis used full-length receptors on cells, in SPR reconstituted receptors with no transmembrane and intracellular domains are bound to chips. The absence of these domains and associated membrane stabilization may contribute to differences in VEGF-A binding. For example, previous studies showed that mutation of the transmembrane domain of NRP1 significantly reduced VEGF-A binding^88^. Thus, computational modelers should consider these variations in the measurements of ligand-binding assays to more consciously choose kinetic values and investigate their impact on model outcomes through a sensitivity analysis across the pM to nM range that we identified across these assays.

### Study limitations

This study has three limitations: (1) **Small sample sizes in datasets**: The small sample size may overestimate the confidence interval or prediction interval and result in questionable standardized values. The overestimated confidence interval also affects the results of statistical tests in which we compare group means. Indeed, one of our datasets, VEGF-A:NRP1 binding affinities measured by SPR, included only three studies, which yielded wide prediction intervals and insignificant differences in measurements between SPR and the radioligand assay. Additional related studies should be accumulated to reduce the possible overestimation of pooled variance. (2) **Lack of moderator analysis**: Moderator analysis is usually performed to identify the moderators that affect heterogeneity in datasets. Our analysis showed high heterogeneity between studies for each dataset. For example, the lower bound of the 95% prediction intervals for some datasets was smaller than 0 (e.g., −4.7 to 10.0 nM for binding affinities of VEGF-A:NRP1 measured by SPR). Most values of I^2^, which indicates the proportion of total variability attributed to between-study variability, exceeded 90%, categorizing the standardized values as having high heterogeneity (I^2^ >75%)^89^. However, we lacked the data to perform moderator analysis. We expect that the high heterogeneity between studies in datasets would be caused by different mouse models, cell lines, different ligand–receptor interaction conditions, etc. (3) **Not meta-analysis**: We performed an extensive literature search (*n* = 75 studies) using free-text search-phrase approaches to find targeting measurements. However, we acknowledge that this is not a meta-analysis. A meta-analysis requires an exhaustive literature search using both controlled vocabulary (e.g., MeSH) and text word search. Another best practice for meta-analysis is registration in a database such as PROSPERO^90^. Future studies should build upon the principles that we have established here to elevate the analysis to the level of a meta-analysis.

## Conclusions

Our findings will enable advances in three important areas: (1) **Adipose tissue microenvironment and obesity**: Our results show that obesity does not affect capillary basement membrane thickness even though there is increased capillary basement thickness associated with diabetes. Also, obesity decreases vessel density in adipose tissue while vessel size remains the same. These findings help to identify which structures within the vascular microenvironment are affected by obesity. Further, our results align with previous studies reporting that obese mice have larger adipocytes than lean mice^91,92^. (2) **Understanding of vascular dysfunction**: Our observation that adipose tissue has smaller vessels and higher vessel density compared with tumors underlies the importance of investigating tissue-specific vascular phenotypes. These different phenotypes indicate that tumor and adipose tissue angiogenesis may be differently regulated—despite theories to the contrary, which focus on the hypoxia in both environments. Further, by identifying vessel morphology and considering it alongside VEGF-A kinetic parameters, researchers can examine crosstalk between adipose tissue and tumors via VEGF-A signaling. Indeed, there is evidence that adipocyte-induced VEGF–mTOR signaling increases tumor cell growth and that obesity upregulates the signal^93^. Thus, the enhanced knowledge provided by our analysis should be helpful for the development of angiogenesis-targeting treatment in both tumor and obesity research. 3) **Computational modeling**: With the consolidated data that we provide on adipose tissue and adipose vascular characteristics, systems biology researchers can develop much-needed computational models of obesity. **Therefore, this work has broad potential impact on** biological and biomedical research.

## Acknowledgments

We thank Dr. Vincenza Cifarelli for providing invaluable support through discussions. This material is based upon work supported by the National Science Foundation under Grant No. 1923151. Research reported in this publication was also supported by the National Institute of Heart, Lung, and Blood of the National Institutes of Health under Award Number 7R01HL159946-02. Any opinions, findings, and conclusions or recommendations expressed in this material are those of the author(s) and do not necessarily reflect the views of the National Science Foundation or the National Institutes of Health.

## Data availability statement

The data that support the findings of this study are available in the methods and/or supplementary material of this article.

## Author contribution

Yunjeong Lee, Yingye Fang, and Princess I Imoukhuede designed the research. Yunjeong Lee and Keith Lionel Tukei acquired, analyzed, and interpreted the data. Shobhan Kuila acquired and interpreted the SPR data. Xinming Liu acquired the capillary basement membrane thickness and VEGF-B binding affinity data. Yunjeong Lee, Keith Lionel Tukei, Shobhan Kuila, Yingye Fang, and Princess I Imoukhuede were involved in drafting and revising the manuscript.

## Figure legends

**Figure S1.**
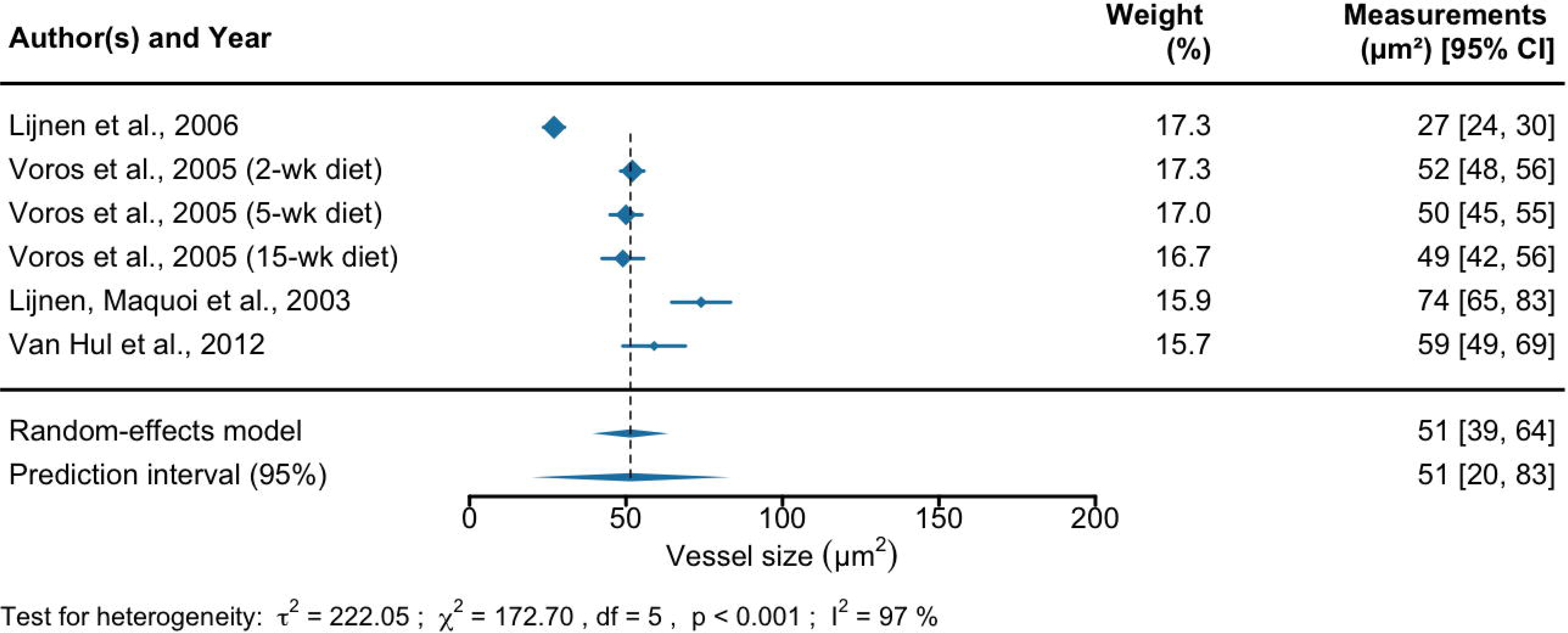
Forest plots of the analysis of vessel size (µm^2^) measured in adipose tissue of lean mice across studies. The first, third, and last columns represent the list of references, their weights in the analysis, and measurements with a 95% confidence interval, respectively. The size of the blue diamond for each study represents its statistical weight in the analysis. The two diamonds below the table and the black dashed line show the combined measurements across studies. The upper diamond represents the mean and 95% confidence interval, and the lower diamond represents the mean and 95% prediction interval. The results of the heterogeneity test are represented by the estimate of between-studies variance (*τ*^2^), Cochran’s *Q*-test statistic for heterogeneity (*χ*^2^), degrees of freedom (df), p-value (p), and proportion of heterogeneity-induced variability among the total variability (I^2^). Abbreviation: week (wk).

**Figure S2.**
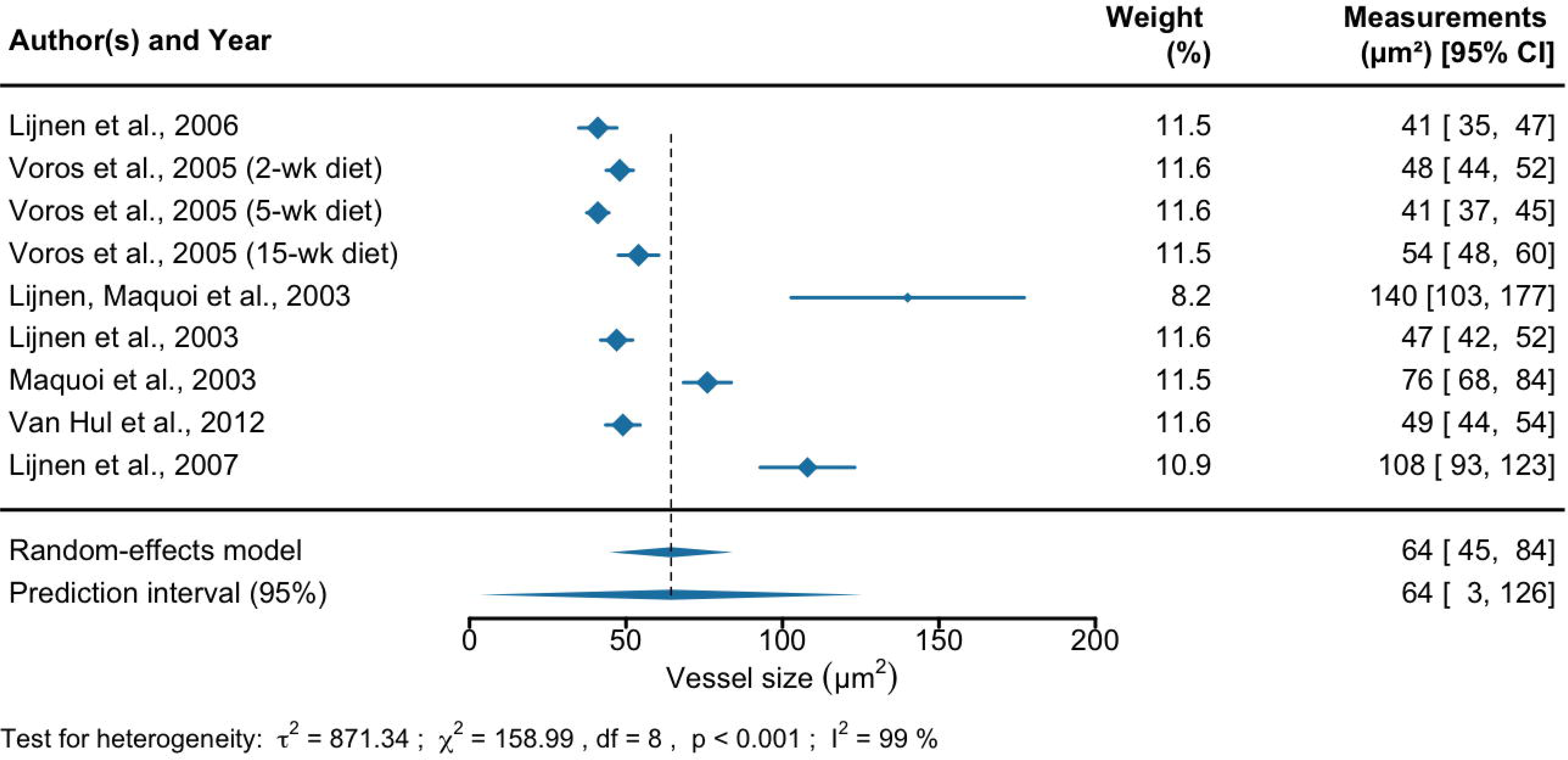
Forest plots of the analysis of vessel size (µm^2^) measured in adipose tissue of diet-induced obese mice across studies. The first, third, and last columns represent the list of references, their weights in the analysis, and measurements with a 95% confidence interval, respectively. The size of the blue diamond for each study represents its statistical weight in the analysis. The two diamonds below the table and the black dashed line show the combined measurements across studies. The upper diamond represents the mean and 95% confidence interval, and the lower diamond represents the mean and 95% prediction interval. The results of the heterogeneity test are represented by the estimate of between-studies variance (*τ*^2^), Cochran’s *Q*-test statistic for heterogeneity (*χ*^2^), degrees of freedom (df), p-value (p), and proportion of heterogeneity-induced variability among the total variability (I^2^). Abbreviation: week (wk).

**Figure S3.**
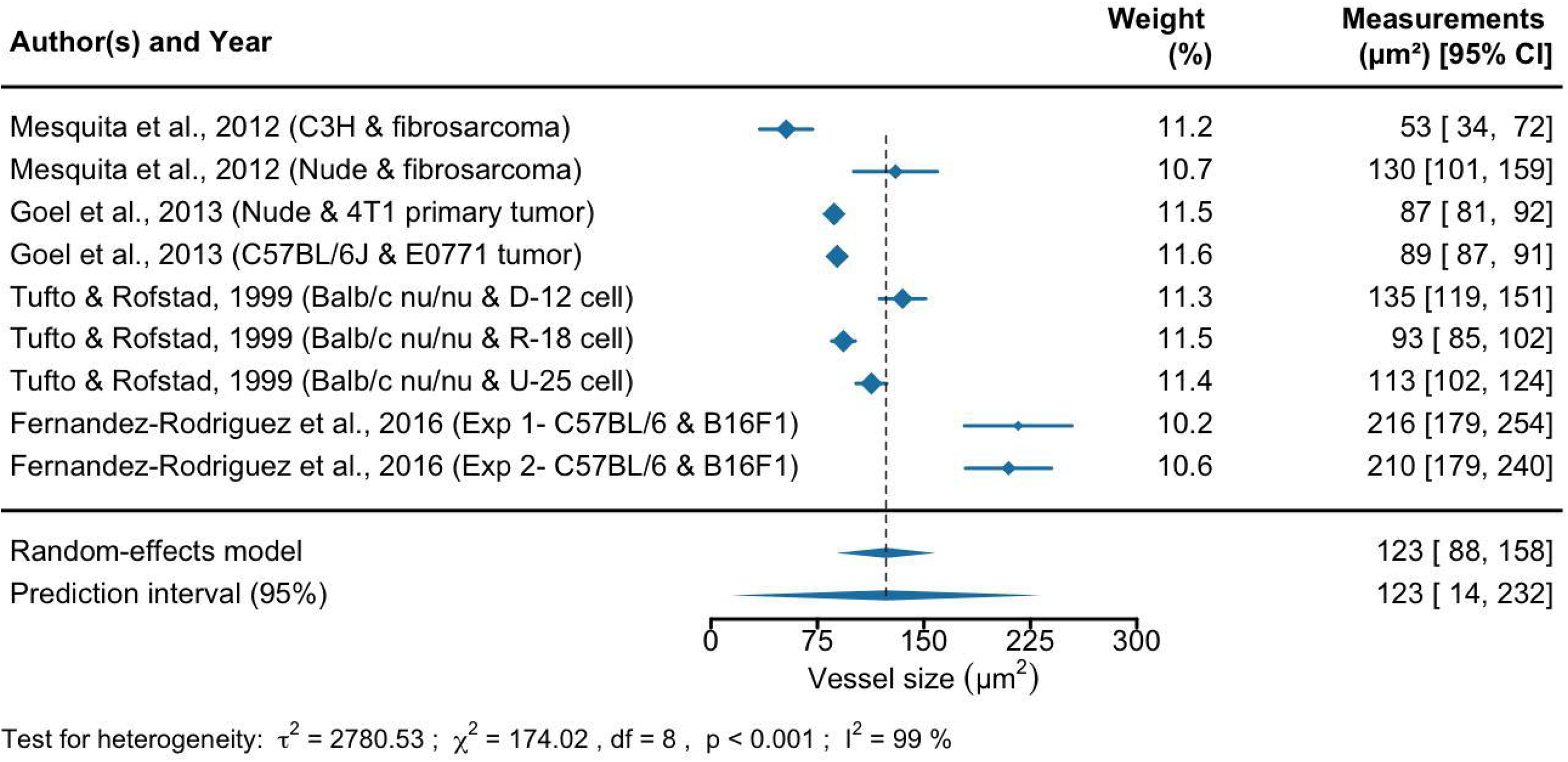
Forest plots of the analysis of vessel size (µm^2^) measured in mouse tumors across studies. The first, third, and last columns represent the list of references, their weights in the analysis, and measurements with a 95% confidence interval, respectively. The size of the blue diamond for each study represents its statistical weight in the analysis. The two diamonds below the table and the black dashed line show the combined measurements across studies. The upper diamond represents the mean and 95% confidence interval, and the lower diamond represents the mean and 95% prediction interval. The results of the heterogeneity test are represented by the estimate of between-studies variance (*τ*^2^), Cochran’s *Q*-test statistic for heterogeneity (*χ*^2^), degrees of freedom (df), p-value (p), and proportion of heterogeneity-induced variability among the total variability (I^2^). The reference names include mouse strain and tumor lines. Abbreviation: experiment (exp).

**Figure S4.**
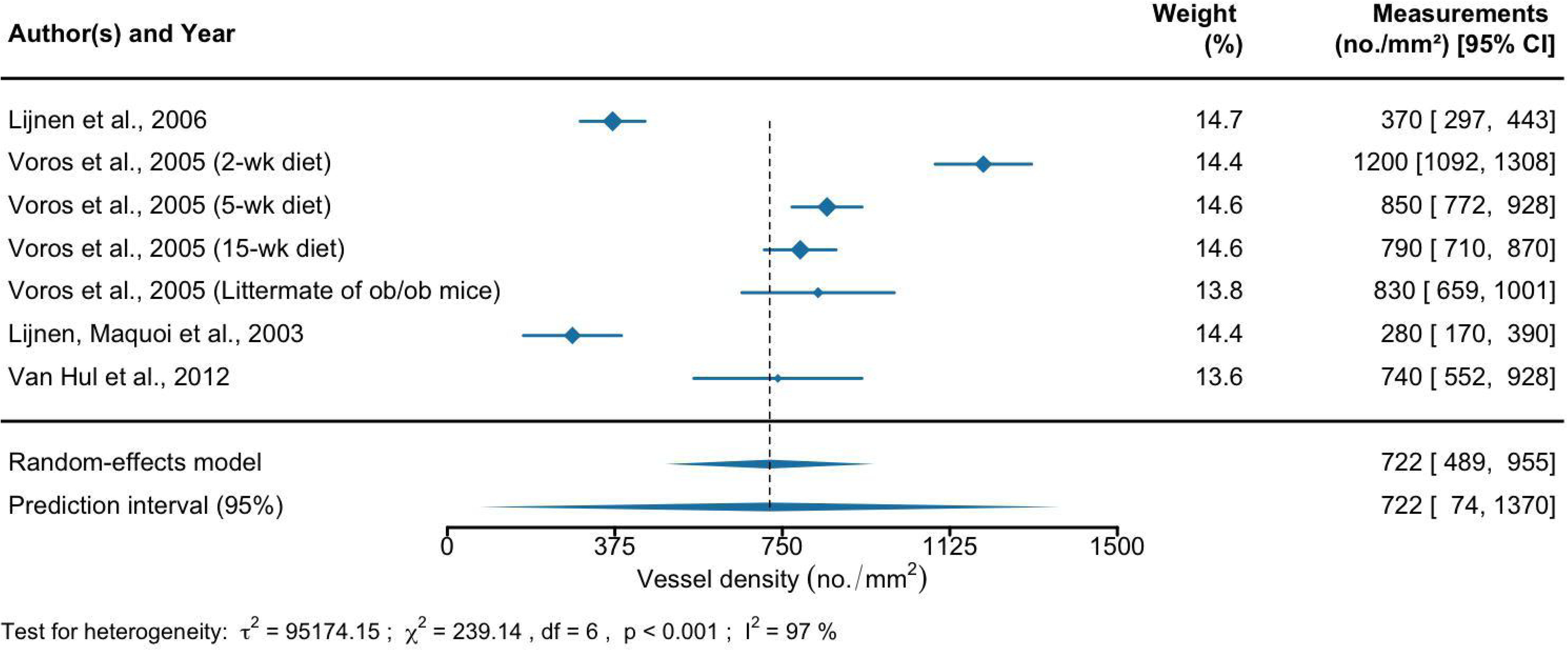
Forest plots of the analysis of vessel density (no./mm^2^) measured in adipose tissue of lean mice across studies. The first, third, and last columns represent the list of references, their weights in the analysis, and measurements with a 95% confidence interval, respectively. The size of the blue diamond for each study represents its statistical weight in the analysis. The two diamonds below the table and the black dashed line show the combined measurements across studies. The upper diamond represents the mean and 95% confidence interval, and the lower diamond represents the mean and 95% prediction interval. The results of the heterogeneity test are represented by the estimate of between-studies variance (*τ*^2^), Cochran’s *Q*-test statistic for heterogeneity (*χ*^2^), degrees of freedom (df), p-value (p), and proportion of heterogeneity-induced variability among the total variability (I^2^). Abbreviation: week (wk).

**Figure S5.**
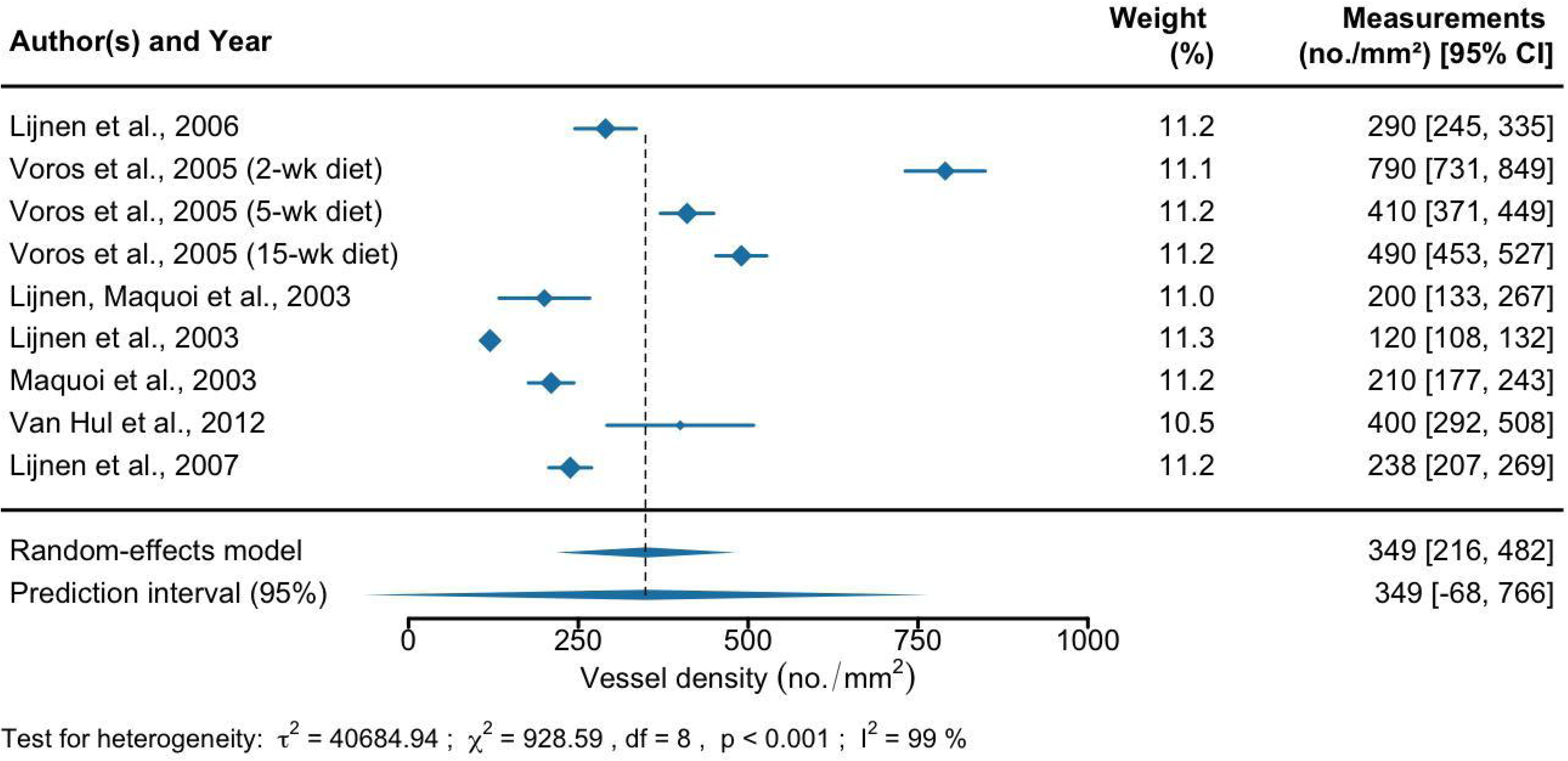
Forest plots of the analysis of vessel density (no./mm^2^) measured in adipose tissue of diet-induced obese mice across studies. The first, third, and last columns represent the list of references, their weights in the analysis, and measurements with a 95% confidence interval, respectively. The size of the blue diamond for each study represents its statistical weight in the analysis. The two diamonds below the table and the black dashed line show the combined measurements across studies. The upper diamond represents the mean and 95% confidence interval, and the lower diamond represents the mean and 95% prediction interval. The results of the heterogeneity test are represented by the estimate of between-studies variance (*τ*^2^), Cochran’s *Q*-test statistic for heterogeneity (*χ*^2^), degrees of freedom (df), p-value (p), and proportion of heterogeneity-induced variability among the total variability (I^2^). Abbreviation: week (wk).

**Figure S6.**
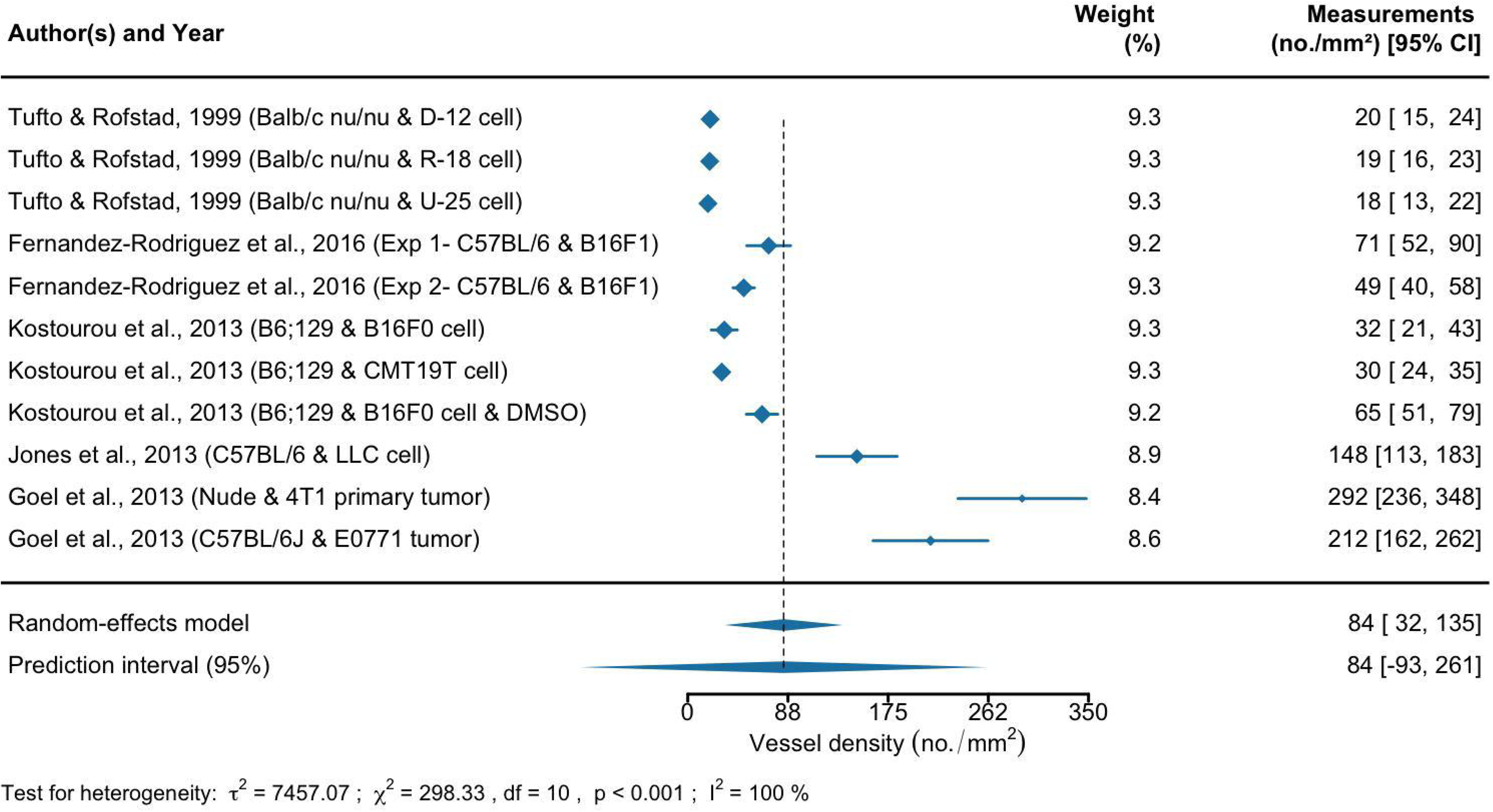
Forest plots of the analysis of vessel density (no./mm^2^) measured in mouse tumors across studies. The first, third, and last columns represent the list of references, their weights in the analysis, and measurements with a 95% confidence interval, respectively. The size of the blue diamond for each study represents its statistical weight in the analysis. The two diamonds below the table and the black dashed line show the combined measurements across studies. The upper diamond represents the mean and 95% confidence interval, and the lower diamond represents the mean and 95% prediction interval. The results of the heterogeneity test are represented by the estimate of between-studies variance (*τ*^2^), Cochran’s *Q*-test statistic for heterogeneity (*χ*^2^), degrees of freedom (df), p-value (p), and proportion of heterogeneity-induced variability among the total variability (I^2^). The reference names include mouse strain and tumor lines. Abbreviation: experiment (exp) and dimethyl sulfoxide (DMSO).

**Figure S7.**
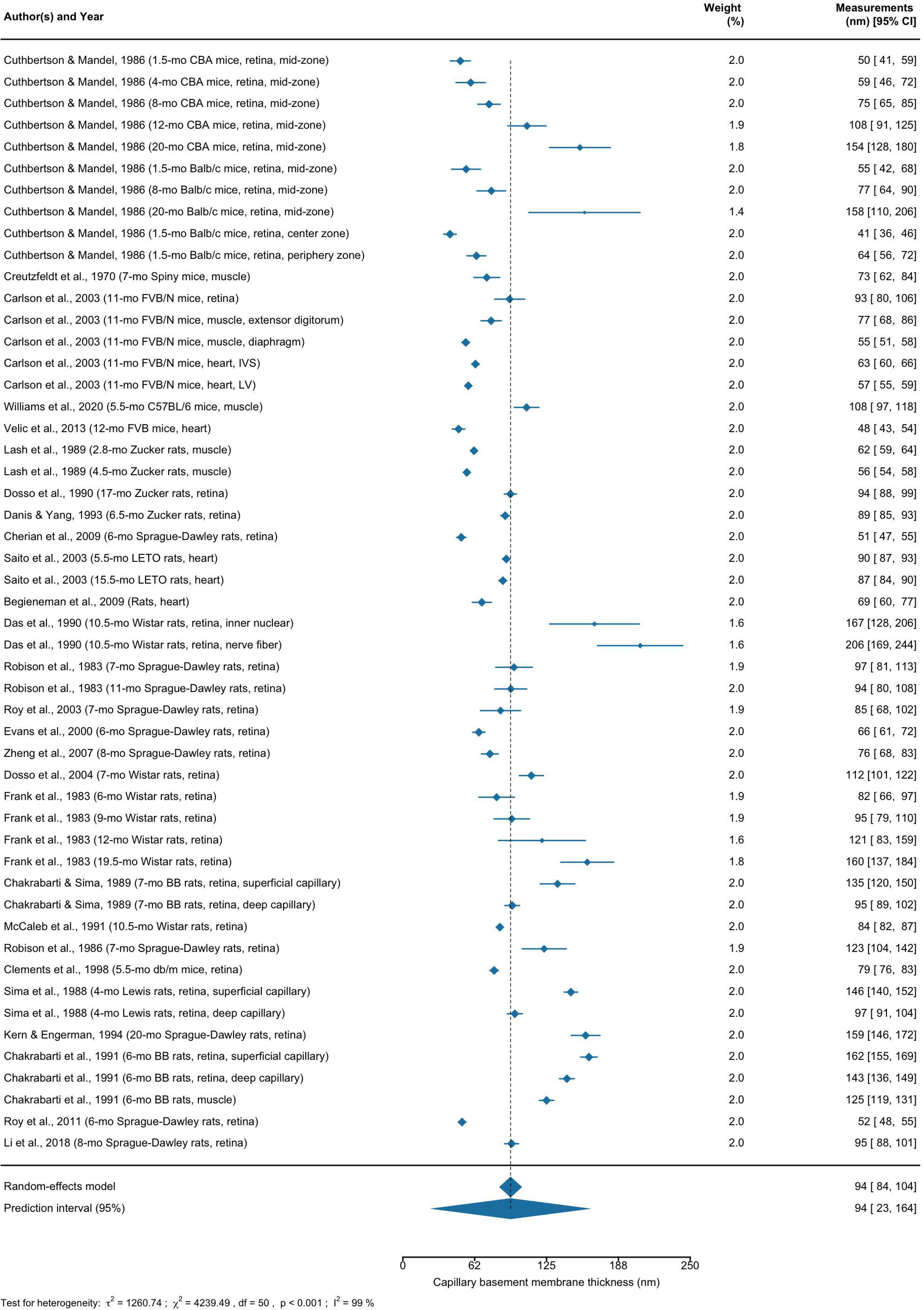
Forest plots of the analysis of capillary basement membrane thickness (nm) measured in tissues of lean mice and rats across studies. The first, third, and last columns represent the list of references, their weights in the analysis, and measurements with a 95% confidence interval, respectively. The size of the blue diamond for each study represents its statistical weight in the analysis. The two diamonds below the table and the black dashed line show the combined measurements across studies. The upper diamond represents the mean and 95% confidence interval, and the lower diamond represents the mean and 95% prediction interval. The results of the heterogeneity test are represented by the estimate of between-studies variance (*τ*^2^), Cochran’s *Q*-test statistic for heterogeneity (*χ*^2^), degrees of freedom (df), p-value (p), and proportion of heterogeneity-induced variability among the total variability (I^2^). Abbreviation: month (mo), interventricular septal (IVS), and left ventricular (LV).

**Figure S8.**
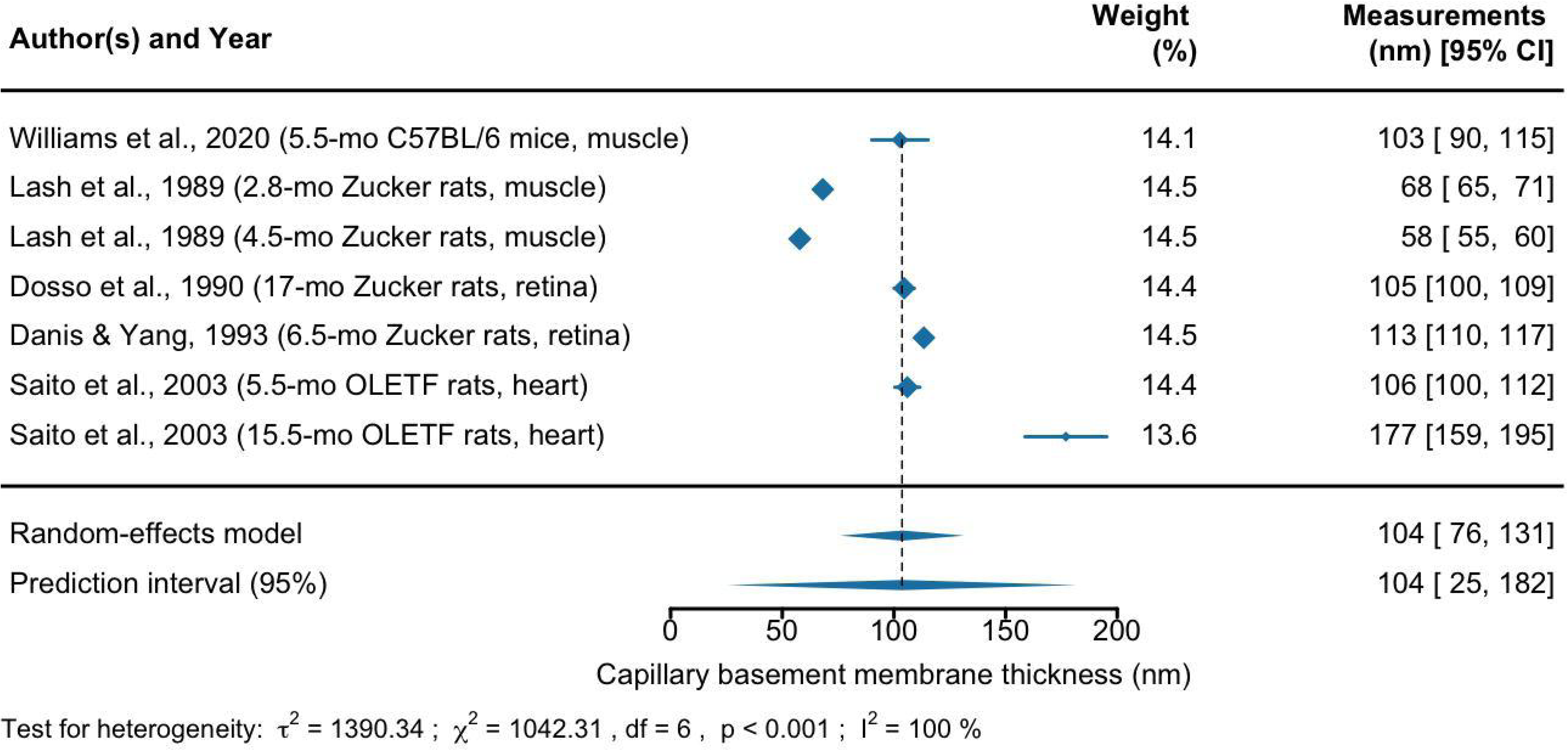
Forest plots of the analysis of capillary basement membrane thickness (nm) measured in tissues of obese mice and rats across studies. The first, third, and last columns represent the list of references, their weights in the analysis, and measurements with a 95% confidence interval, respectively. The size of the blue diamond for each study represents its statistical weight in the analysis. The two diamonds below the table and the black dashed line show the combined measurements across studies. The upper diamond represents the mean and 95% confidence interval, and the lower diamond represents the mean and 95% prediction interval. The results of the heterogeneity test are represented by the estimate of between-studies variance (*τ*^2^), Cochran’s *Q*-test statistic for heterogeneity (*χ*^2^), degrees of freedom (df), p-value (p), and proportion of heterogeneity-induced variability among the total variability (I^2^). Abbreviation: month (mo).

**Figure S9.**
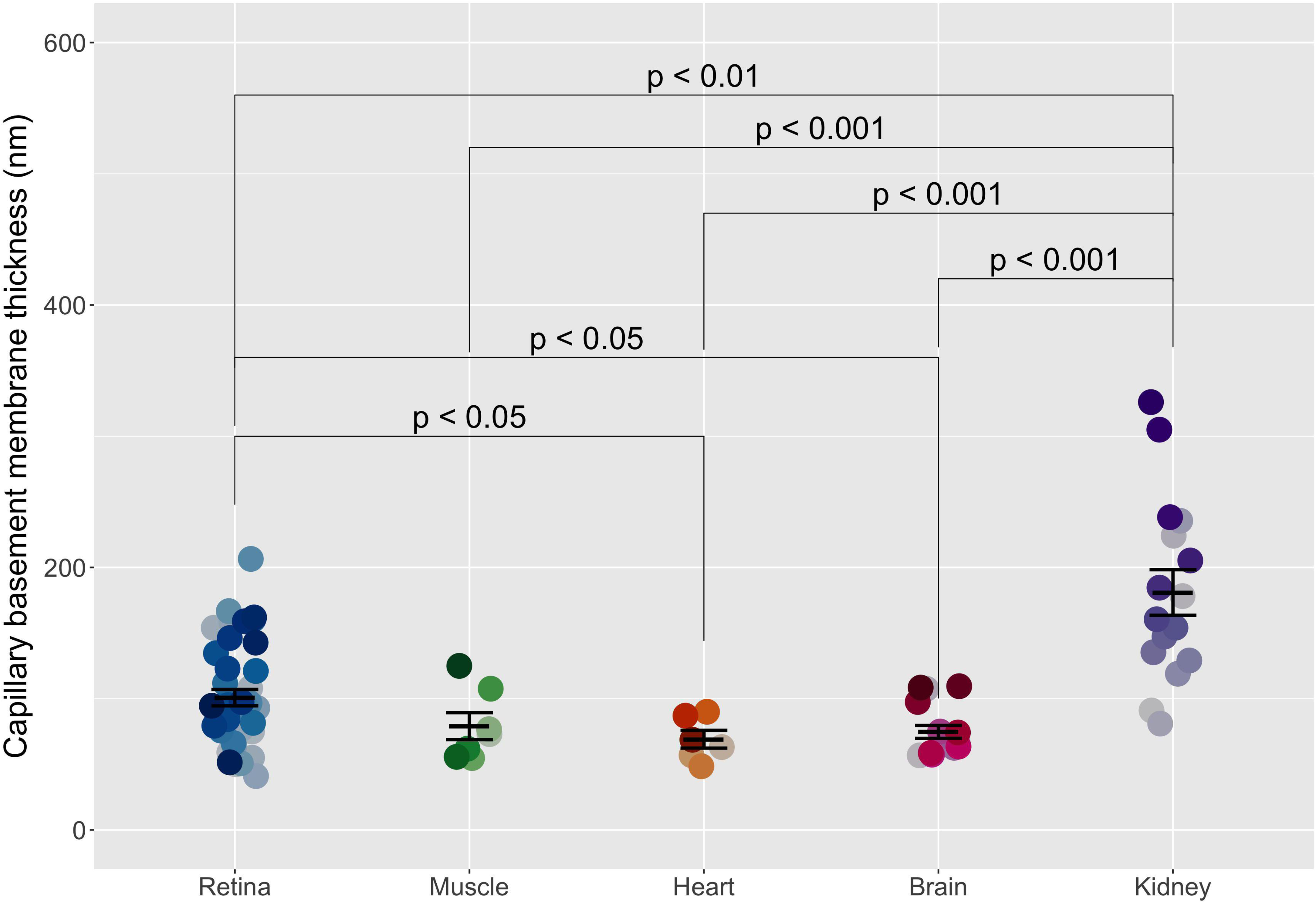
Capillary basement membrane thickness (nm) in tissues of lean mice and rats. The capillary basement membrane thickness in retina, muscle, heart, brain, and kidney in lean mice and rats were compared. Welch’s ANOVA followed by Dunnett’s T3 test was used to compare a pair of tissues. The error bar represents the weighted mean ± standard error of each group.

**Figure S10.**
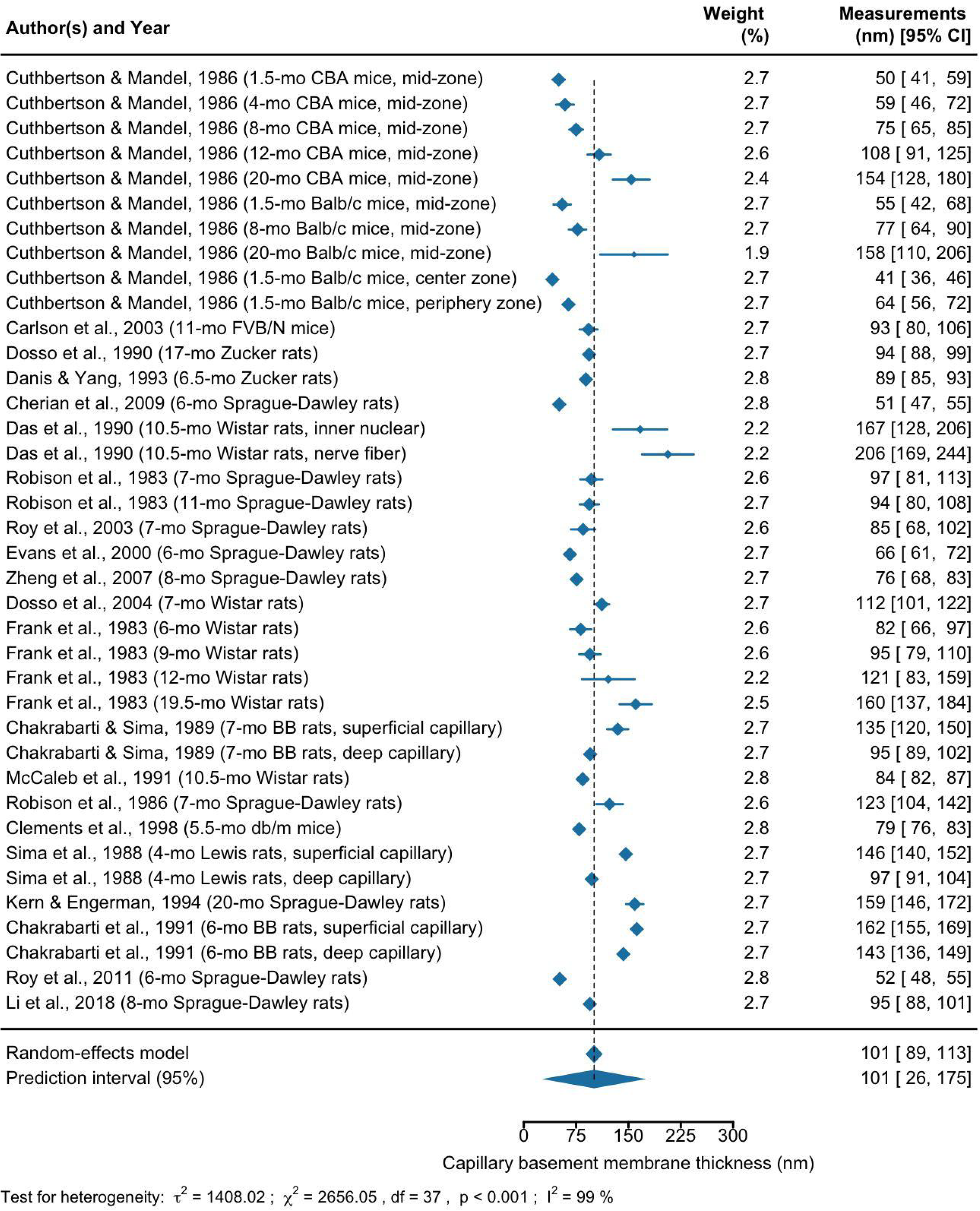
Forest plots of the analysis of capillary basement membrane thickness (nm) measured in retina of lean mice and rats across studies. The first, third, and last columns represent the list of references, their weights in the analysis, and measurements with a 95% confidence interval, respectively. The size of the blue diamond for each study represents its statistical weight in the analysis. The two diamonds below the table and the black dashed line show the combined measurements across studies. The upper diamond represents the mean and 95% confidence interval, and the lower diamond represents the mean and 95% prediction interval. The results of the heterogeneity test are represented by the estimate of between-studies variance (*τ*^2^), Cochran’s *Q*-test statistic for heterogeneity (*χ*^2^), degrees of freedom (df), p-value (p), and proportion of heterogeneity-induced variability among the total variability (I^2^). Abbreviation: month (mo).

**Figure S11.**
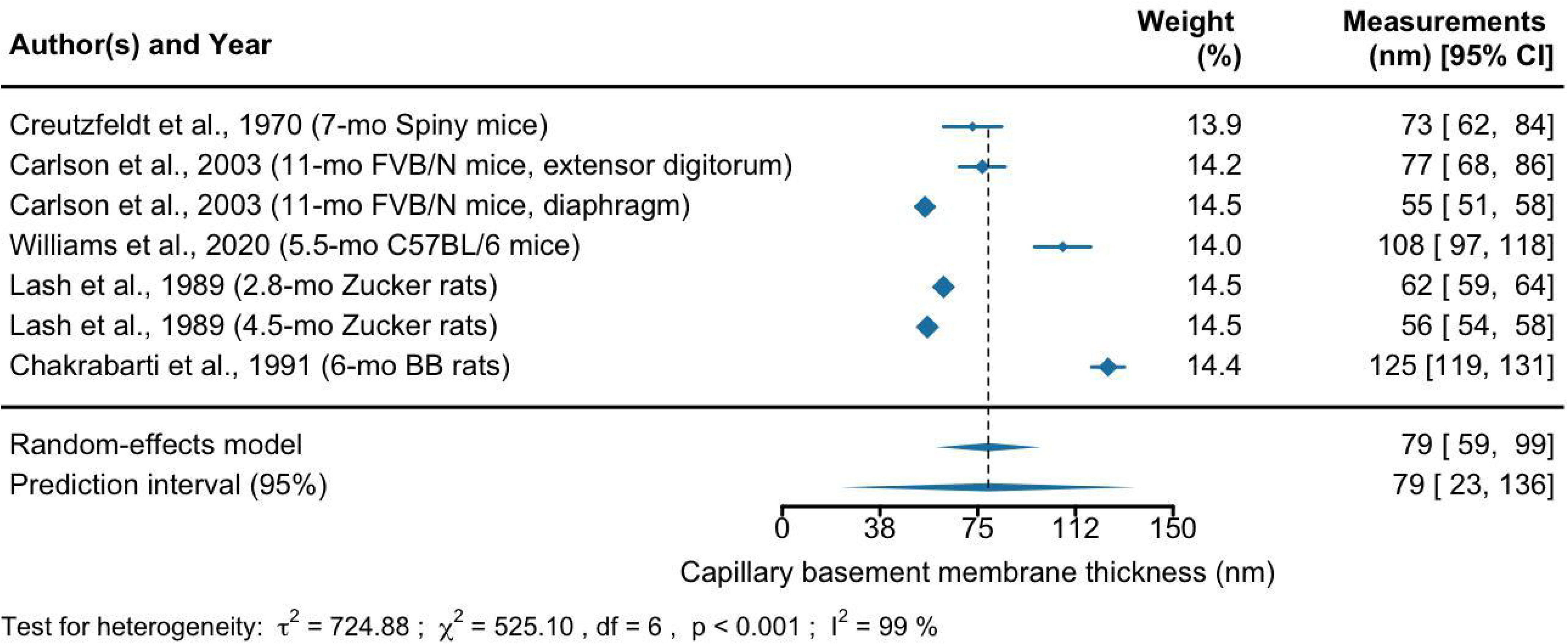
Forest plots of the analysis of capillary basement membrane thickness (nm) measured in muscle of lean mice and rats across studies. The first, third, and last columns represent the list of references, their weights in the analysis, and measurements with a 95% confidence interval, respectively. The size of the blue diamond for each study represents its statistical weight in the analysis. The two diamonds below the table and the black dashed line show the combined measurements across studies. The upper diamond represents the mean and 95% confidence interval, and the lower diamond represents the mean and 95% prediction interval. The results of the heterogeneity test are represented by the estimate of between-studies variance (*τ*^2^), Cochran’s *Q*-test statistic for heterogeneity (*χ*^2^), degrees of freedom (df), p-value (p), and proportion of heterogeneity-induced variability among the total variability (I^2^). Abbreviation: month (mo).

**Figure S12.**
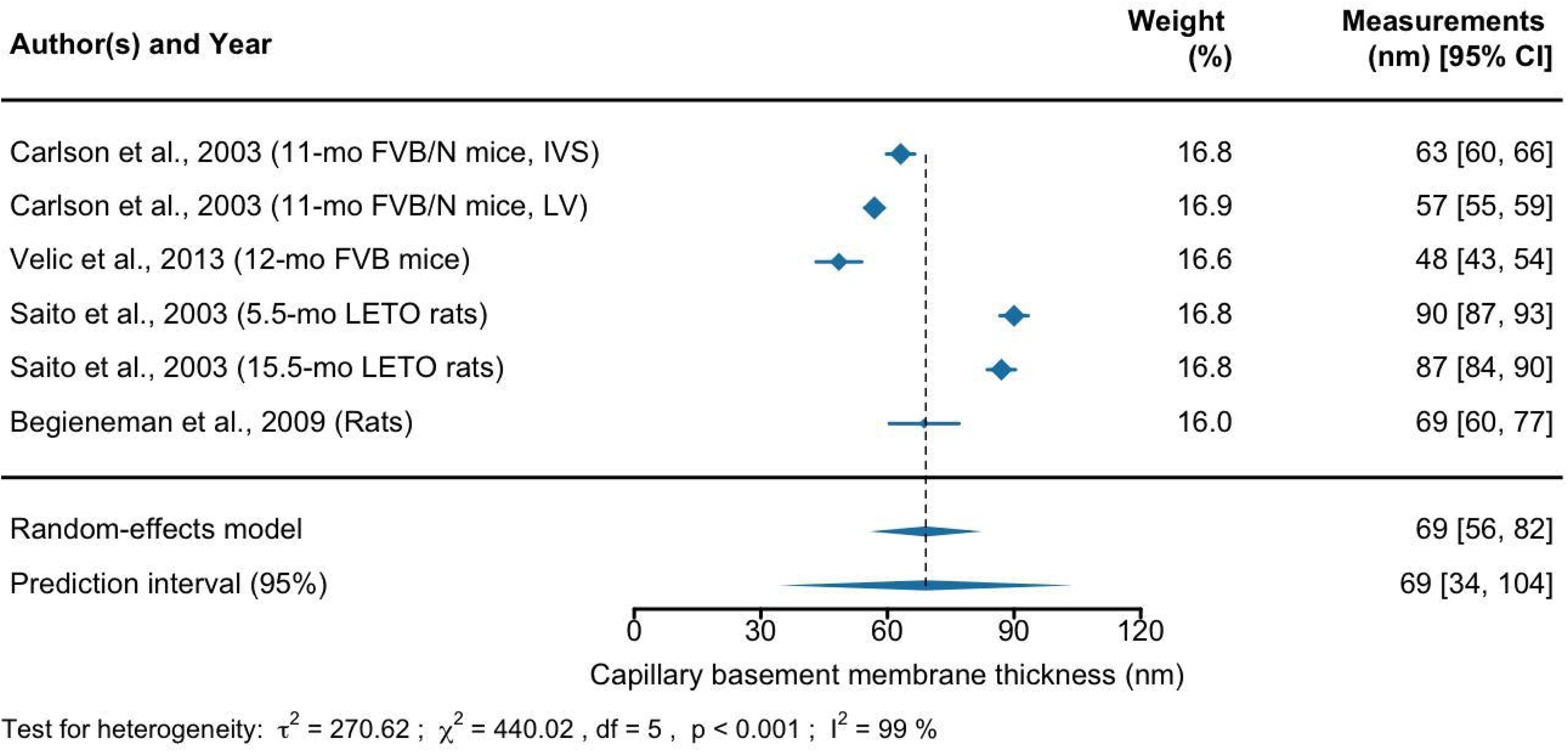
Forest plots of the analysis of capillary basement membrane thickness (nm) measured in heart of lean mice and rats across studies. The first, third, and last columns represent the list of references, their weights in the analysis, and measurements with 95% confidence interval, respectively. The size of the blue diamond for each study represents its statistical weight in the analysis. The two diamonds below the table and the black dashed line show the combined measurements across studies. The upper diamond represents the mean and 95% confidence interval, and the lower diamond represents the mean and 95% prediction interval. The results of the heterogeneity test are represented by the estimate of between-studies variance (*τ*^2^), Cochran’s *Q*-test statistic for heterogeneity (*χ*^2^), degrees of freedom (df), p-value (p), and proportion of heterogeneity-induced variability among the total variability (I^2^). Abbreviation: month (mo), interventricular septal (IVS), and left ventricular (LV).

**Figure S13.**
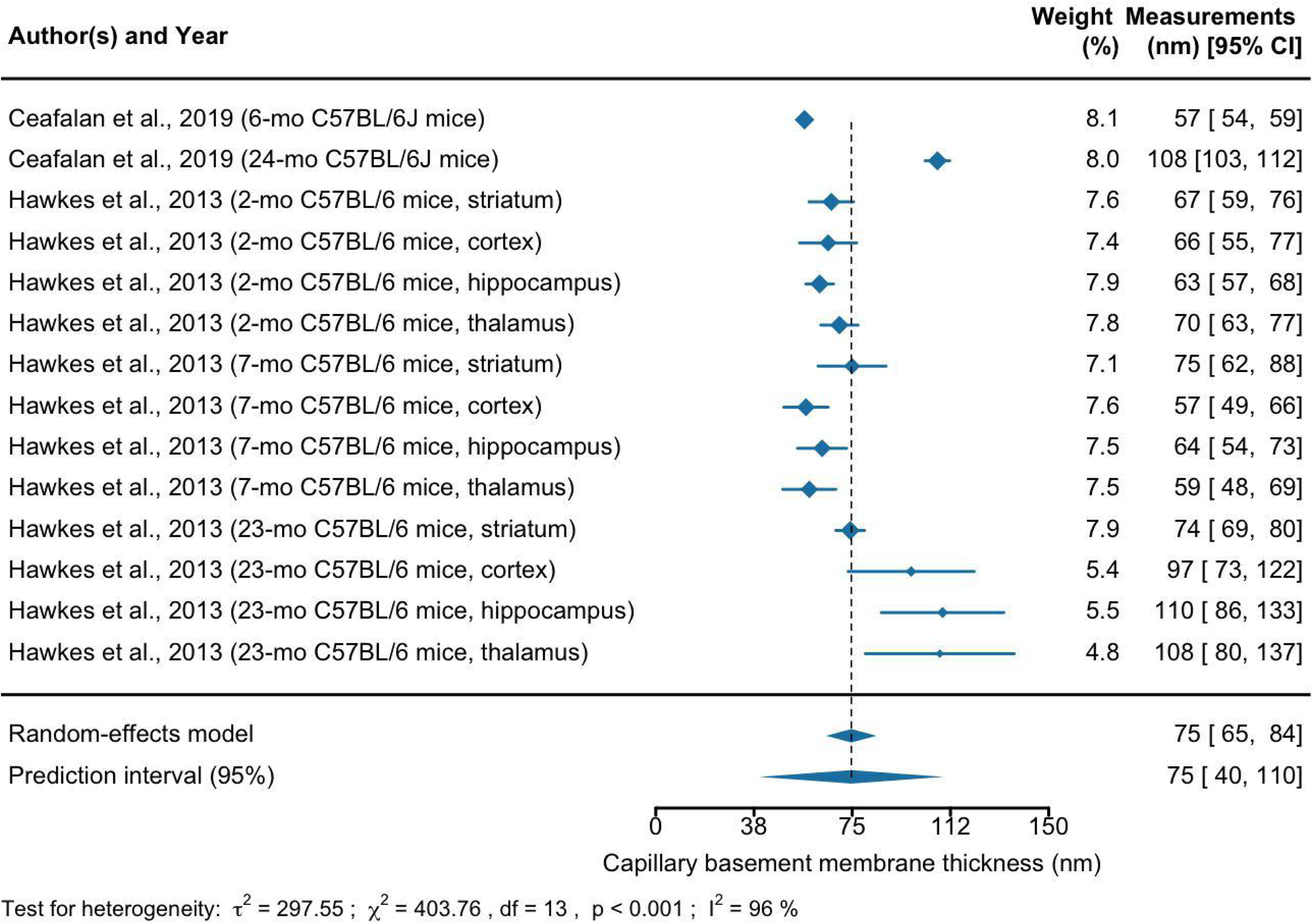
Forest plots of the analysis of capillary basement membrane thickness (nm) measured in brain of lean mice and rats across studies. The first, third, and last columns represent the list of references, their weights in the analysis, and measurements with a 95% confidence interval, respectively. The size of the blue diamond for each study represents its statistical weight in the analysis. The two diamonds below the table and the black dashed line show the combined measurements across studies. The upper diamond represents the mean and 95% confidence interval, and the lower diamond represents the mean and 95% prediction interval. The results of the heterogeneity test are represented by the estimate of between-studies variance (*τ*^2^), Cochran’s *Q*-test statistic for heterogeneity (*χ*^2^), degrees of freedom (df), p-value (p), and proportion of heterogeneity-induced variability among the total variability (I^2^). Abbreviation: month (mo).

**Figure S14.**
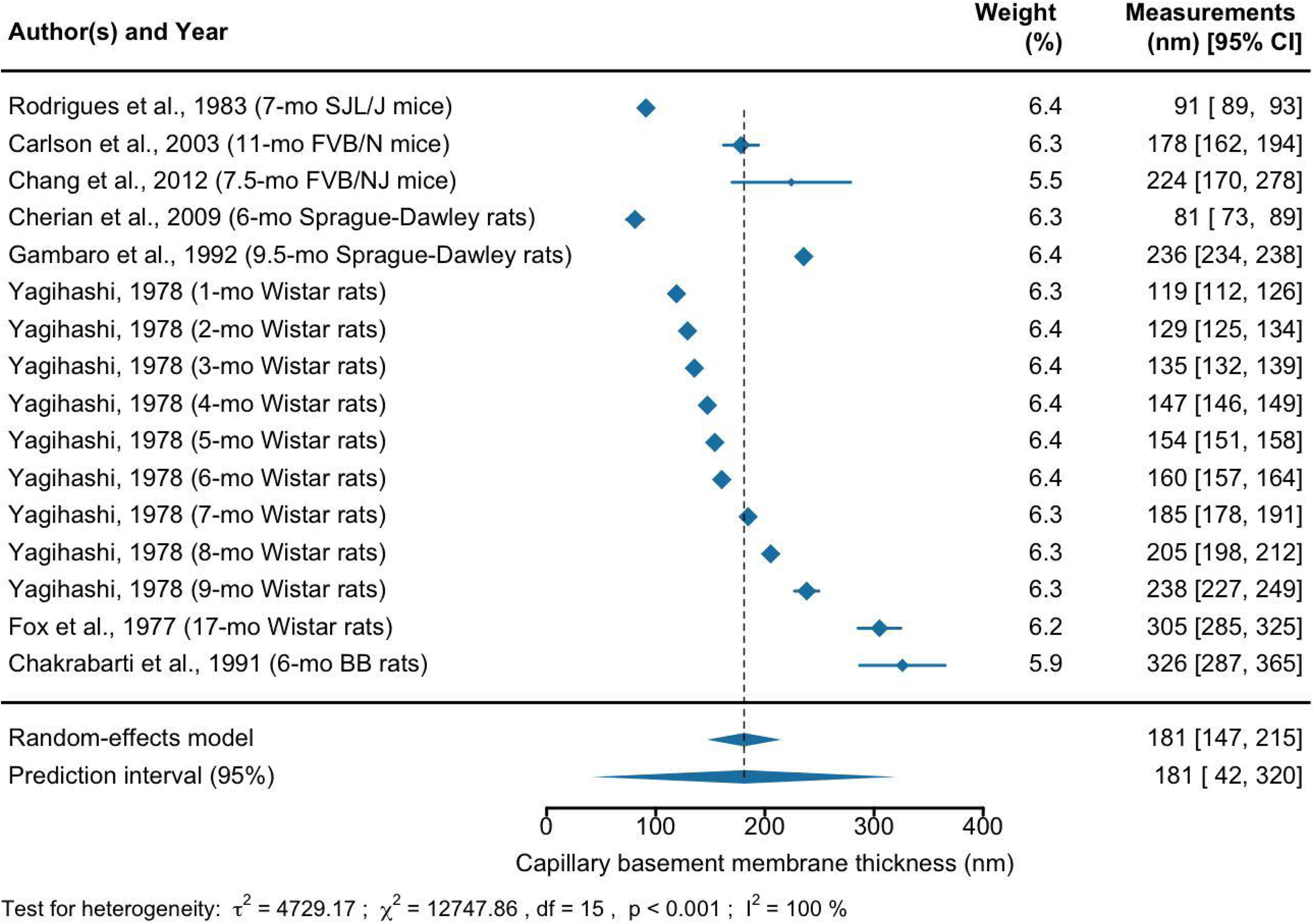
Forest plots of the analysis of capillary basement membrane thickness (nm) measured in kidney of lean mice and rats across studies. The first, third, and last columns represent the list of references, their weights in the analysis, and measurements with a 95% confidence interval, respectively. The size of the blue diamond for each study represents its statistical weight in the analysis. The two diamonds below the table and the black dashed line show the combined measurements across studies. The upper diamond represents the mean and 95% confidence interval, and the lower diamond represents the mean and 95% prediction interval. The results of the heterogeneity test are represented by the estimate of between-studies variance (*τ*^2^), Cochran’s *Q*-test statistic for heterogeneity (*χ*^2^), degrees of freedom (df), p-value (p), and proportion of heterogeneity-induced variability among the total variability (I^2^). Abbreviation: month (mo).

**Figure S15.**
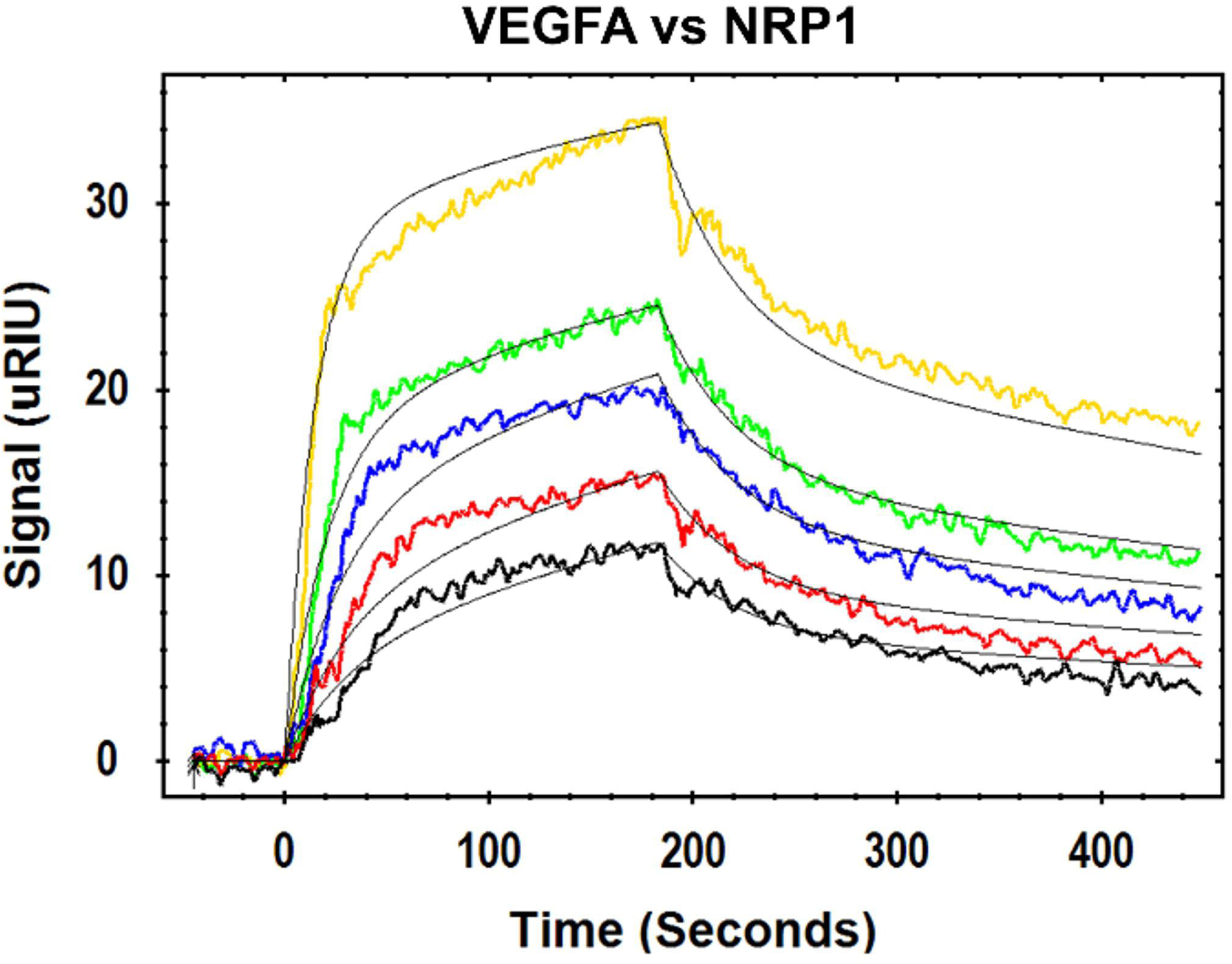
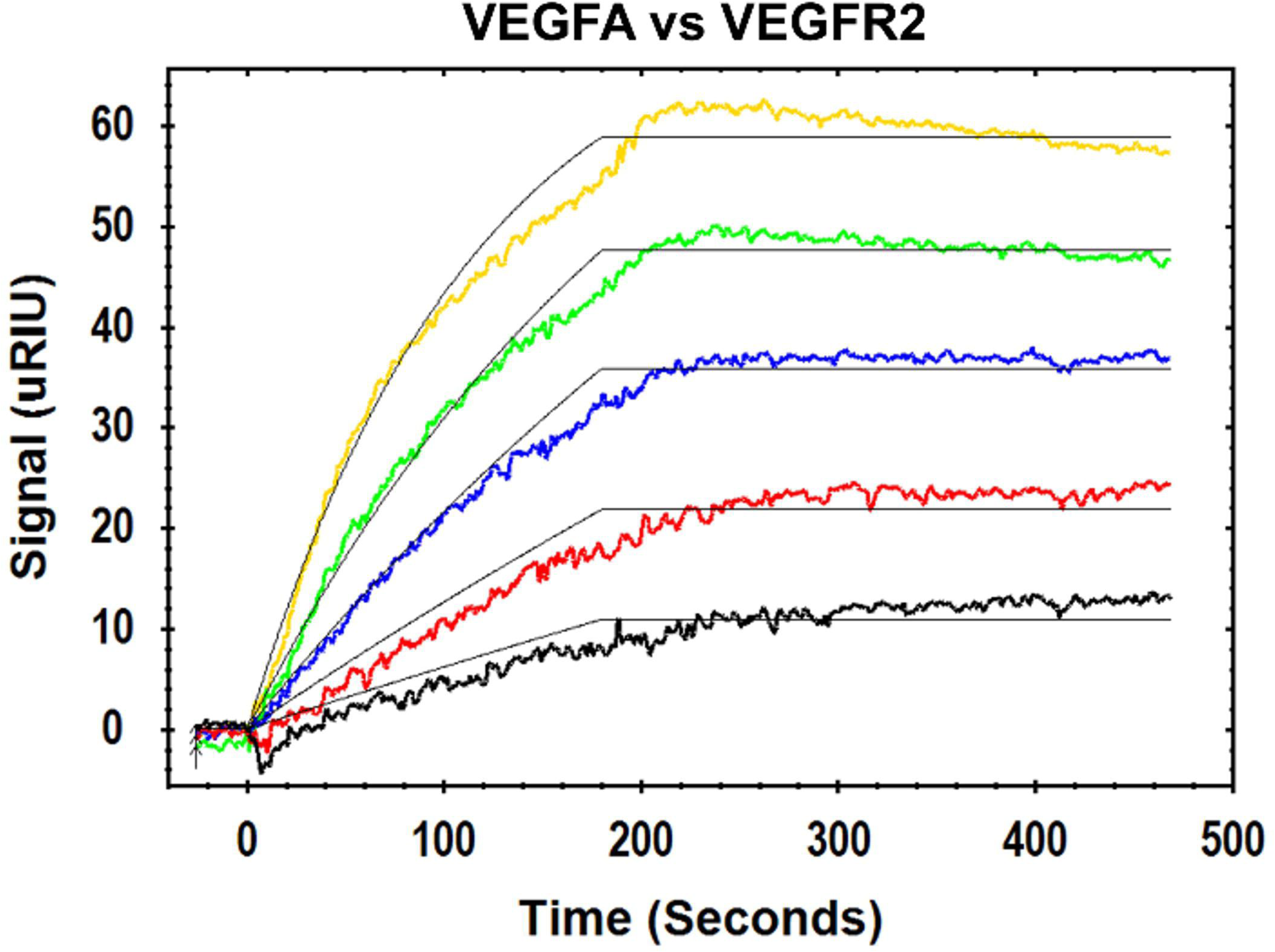
Interaction kinetics of VEGF-A:NRP1 and VEGF-A:VEGFR2. NRP1 (A) and VEGFR2 (B) were immobilized and VEGF-A as analyte was passed over them at different concentrations: 50 nM (yellow), 25 nM (green), 12.5 nM (blue), 6.25 nM (red), and 3.125 nM (black). Note: the thin black overlapping lines are fitted curves of a 1:1 Langmuir model drawn with TraceDrawer ver. 1.8.1 software.

**Figure S16.**
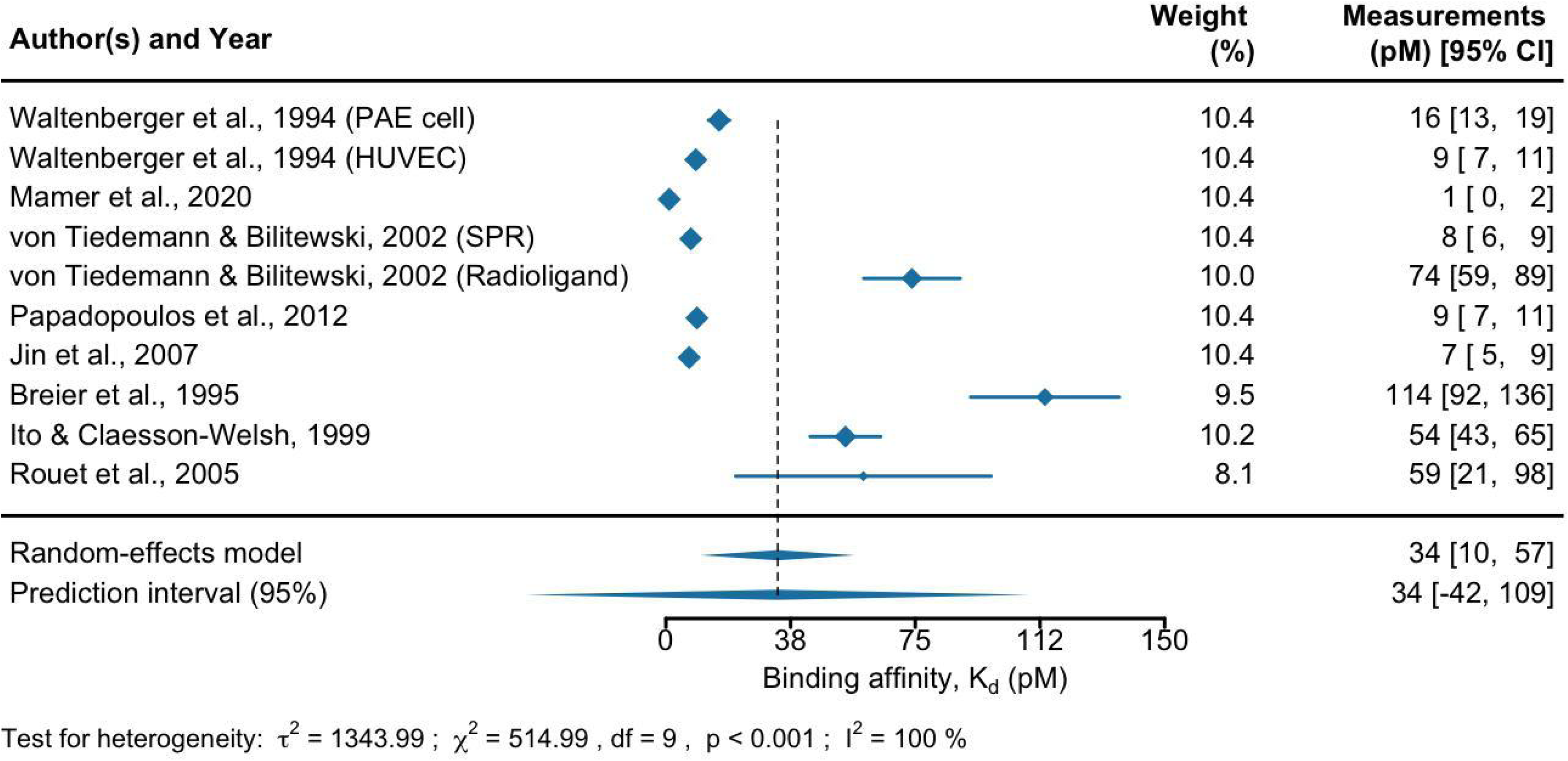
Forest plots of the analysis of binding affinity (pM) of VEGF to VEGFR1 across studies. The first, third, and last columns represent the list of references, their weights in the analysis, and measurements with a 95% confidence interval, respectively. The size of the blue diamond for each study represents its statistical weight in the analysis. The two diamonds below the table and the black dashed line show the combined measurements across studies. The upper diamond represents the mean and 95% confidence interval, and the lower diamond represents the mean and 95% prediction interval. The results of the heterogeneity test are represented by the estimate of between-studies variance (*τ*^2^), Cochran’s *Q*-test statistic for heterogeneity (*χ*^2^), degrees of freedom (df), p-value (p), and proportion of heterogeneity-induced variability among the total variability (I^2^). Abbreviation: porcine aortic endothelial (PAE), human umbilical vein endothelial cell (HUVEC), and surface plasmon resonance (SPR).

**Figure S17.**
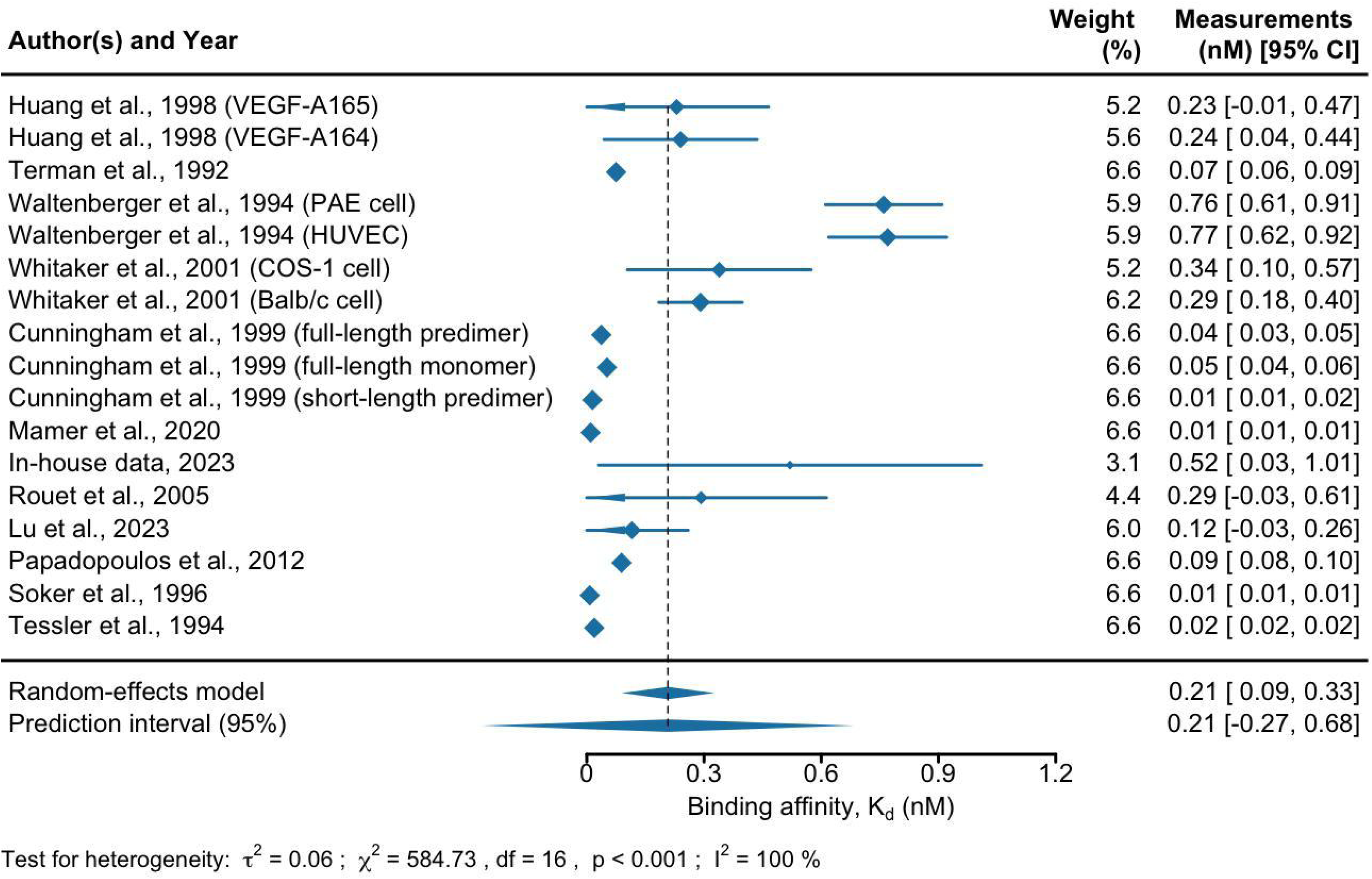
Forest plots of the analysis of binding affinity (nM) of VEGF to VEGFR2 across studies. The first, third, and last columns represent the list of references, their weights in the analysis, and measurements with a 95% confidence interval, respectively. The size of the blue diamond for each study represents its statistical weight in the analysis. The two diamonds below the table and the black dashed line show the combined measurements across studies. The upper diamond represents the mean and 95% confidence interval, and the lower diamond represents the mean and 95% prediction interval. The results of the heterogeneity test are represented by the estimate of between-studies variance (*τ*^2^), Cochran’s *Q*-test statistic for heterogeneity (*χ*^2^), degrees of freedom (df), p-value (p), and proportion of heterogeneity-induced variability among the total variability (I^2^). Abbreviation: porcine aortic endothelial (PAE) and human umbilical vein endothelial cell (HUVEC).

**Figure S18.**
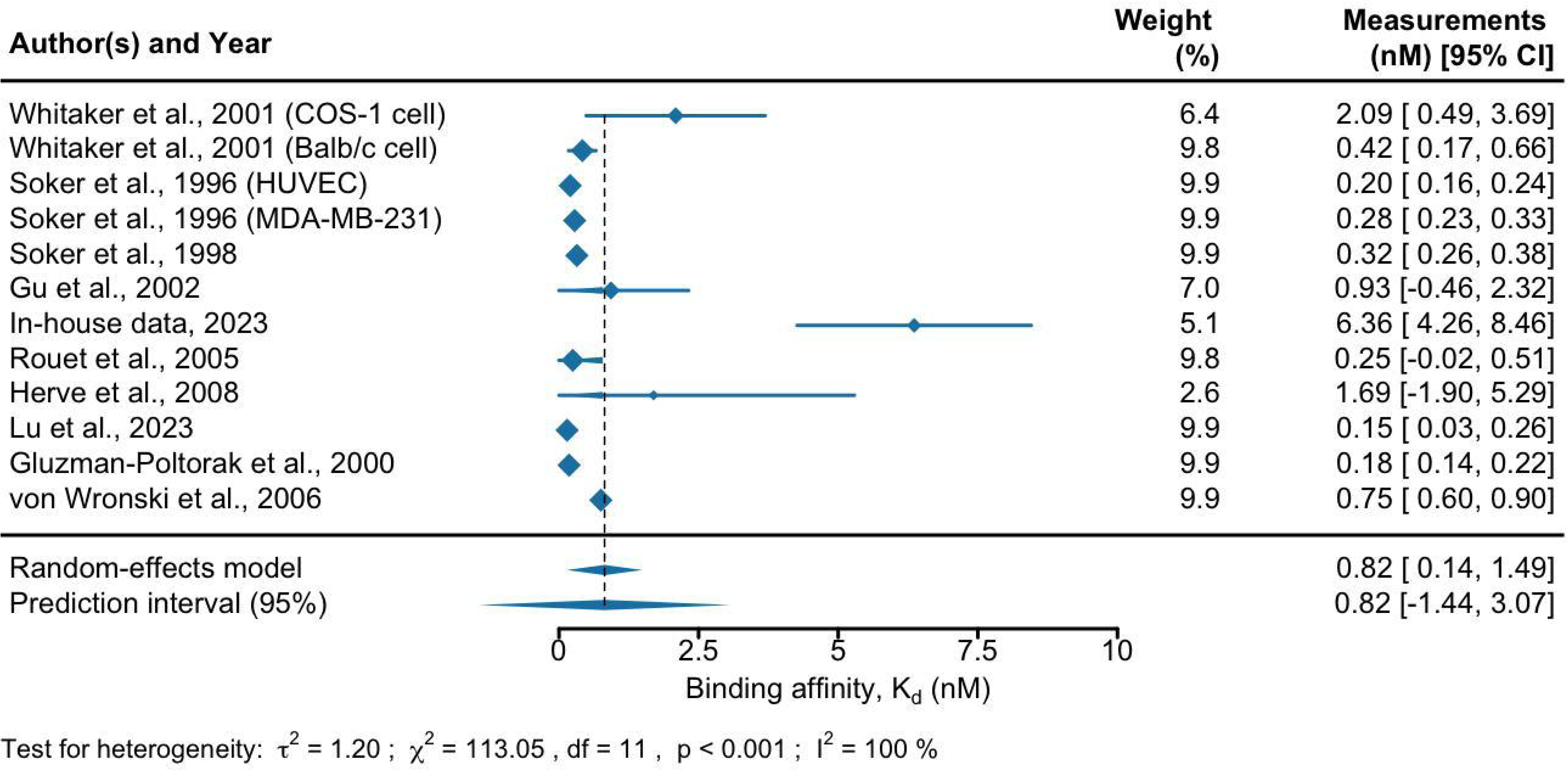
Forest plots of the analysis of binding affinity (nM) of VEGF to NRP1 across studies. The first, third, and last columns represent the list of references, their weights in the analysis, and measurements with a 95% confidence interval, respectively. The size of the blue diamond for each study represents its statistical weight in the analysis. The two diamonds below the table and the black dashed line show the combined measurements across studies. The upper diamond represents the mean and 95% confidence interval, and the lower diamond represents the mean and 95% prediction interval. The results of the heterogeneity test are represented by the estimate of between-studies variance (*τ*^2^), Cochran’s *Q*-test statistic for heterogeneity (*χ*^2^), degrees of freedom (df), p-value (p), and proportion of heterogeneity-induced variability among the total variability (I^2^). Abbreviation: human umbilical vein endothelial cell (HUVEC).

**Figure S19.**
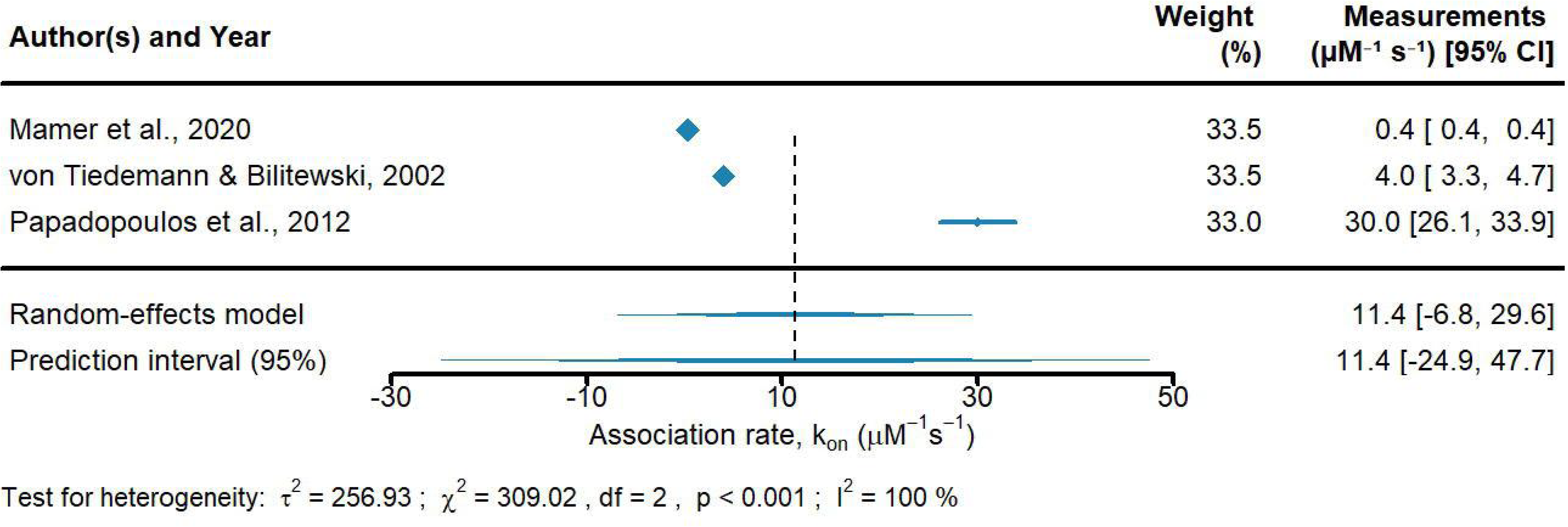
Forest plots of the analysis of association rates, k_on_ (µM^−1^s^−1^) of VEGF to VEGFR1 across studies. The first, third, and last columns represent the list of references, their weights in the analysis, and measurements with a 95% confidence interval, respectively. The size of the blue diamond for each study represents its statistical weight in the analysis. The two diamonds below the table and the black dashed line show the combined measurements across studies. The upper diamond represents the mean and 95% confidence interval, and the lower diamond represents the mean and 95% prediction interval. The results of the heterogeneity test are represented by the estimate of between-studies variance (*τ*^2^), Cochran’s *Q*-test statistic for heterogeneity (*χ*^2^), degrees of freedom (df), p-value (p), and proportion of heterogeneity-induced variability among the total variability (I^2^).

**Figure S20.**
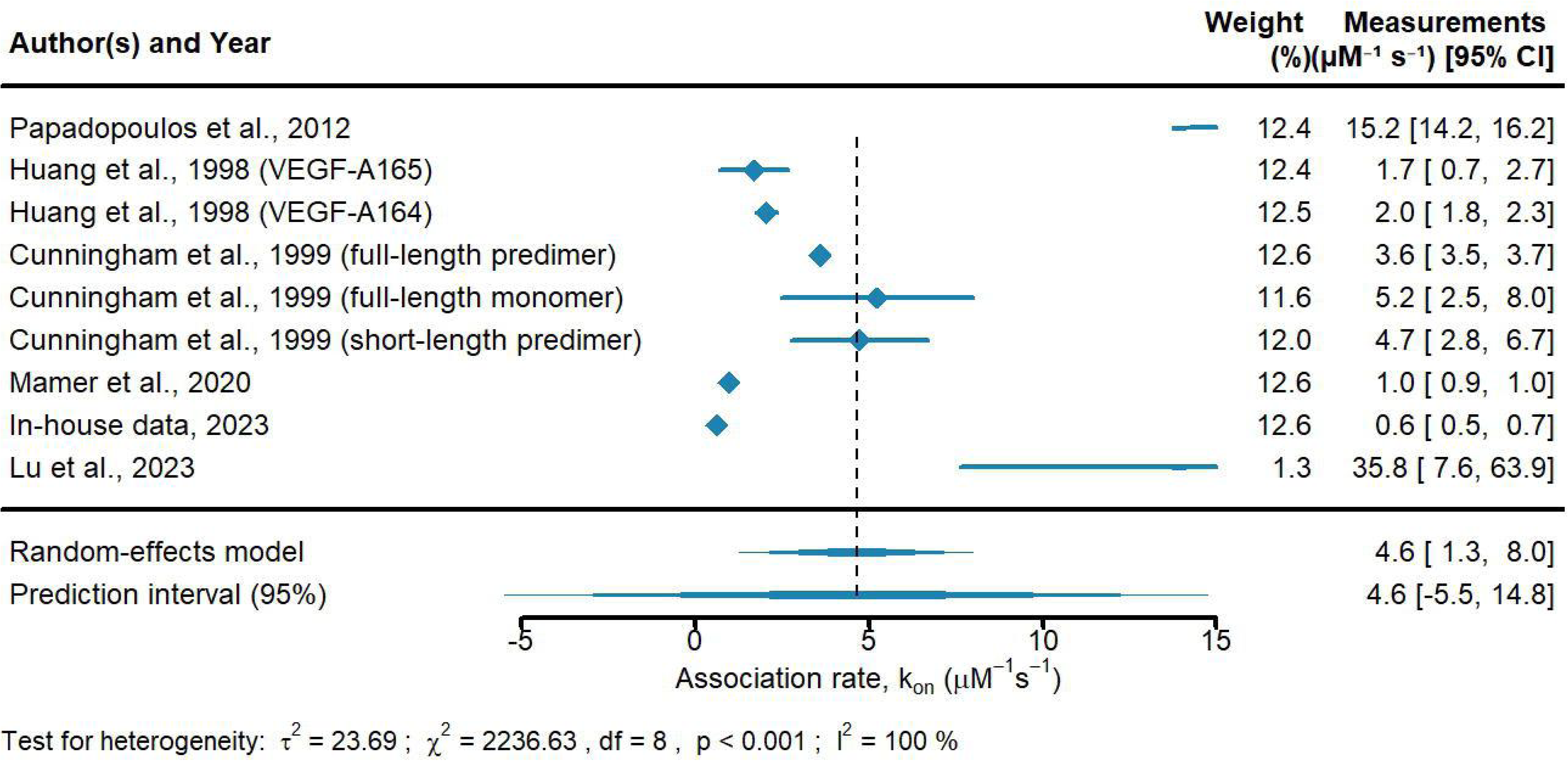
Forest plots of the analysis of association rates, k_on_ (µM^−1^s^−1^) of VEGF to VEGFR2 across studies. The first, third, and last columns represent the list of references, their weights in the analysis, and measurements with a 95% confidence interval, respectively. The size of the blue diamond for each study represents its statistical weight in the analysis. The two diamonds below the table and the black dashed line show the combined measurements across studies. The upper diamond represents the mean and 95% confidence interval, and the lower diamond represents the mean and 95% prediction interval. The results of the heterogeneity test are represented by the estimate of between-studies variance (*τ*^2^), Cochran’s *Q*-test statistic for heterogeneity (*χ*^2^), degrees of freedom (df), p-value (p), and proportion of heterogeneity-induced variability among the total variability (I^2^).

**Figure S21.**
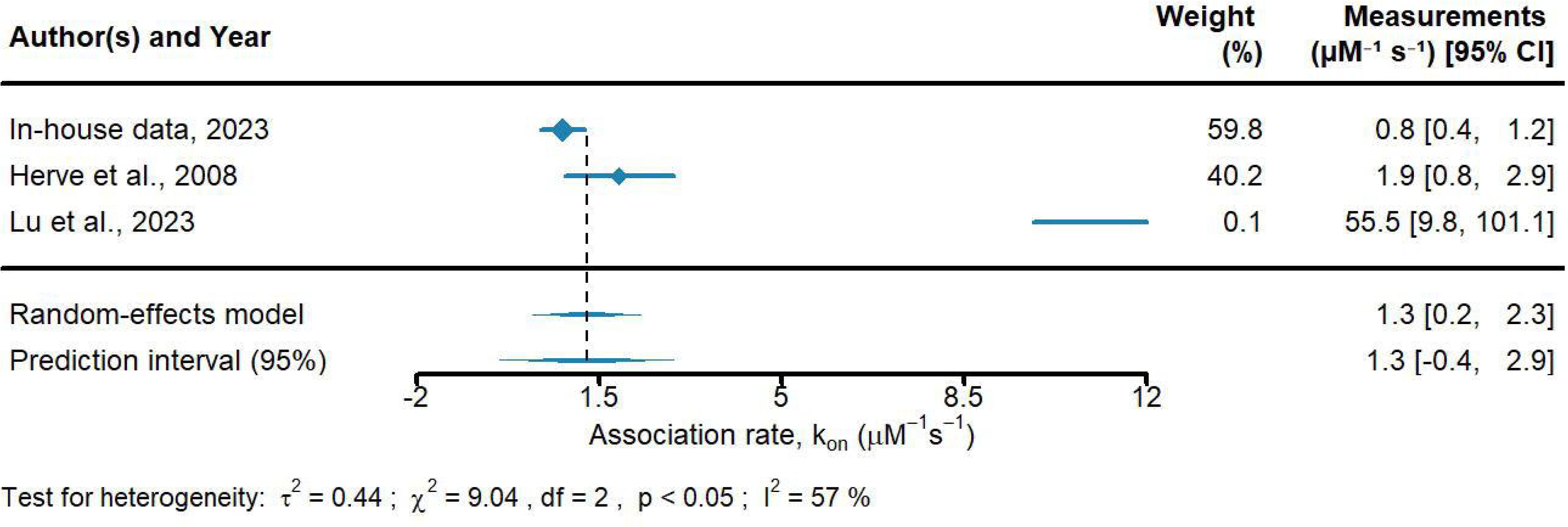
Forest plots of the analysis of association rates, k_on_ (µM^−1^s^−1^) of VEGF to NRP1 across studies. The first, third, and last columns represent the list of references, their weights in the analysis, and measurements with a 95% confidence interval, respectively. The size of the blue diamond for each study represents its statistical weight in the analysis. The two diamonds below the table and the black dashed line show the combined measurements across studies. The upper diamond represents the mean and 95% confidence interval, and the lower diamond represents the mean and 95% prediction interval. The results of the heterogeneity test are represented by the estimate of between-studies variance (*τ*^2^), Cochran’s *Q*-test statistic for heterogeneity (*χ*^2^), degrees of freedom (df), p-value (p), and proportion of heterogeneity-induced variability among the total variability (I^2^).

**Figure S22.**
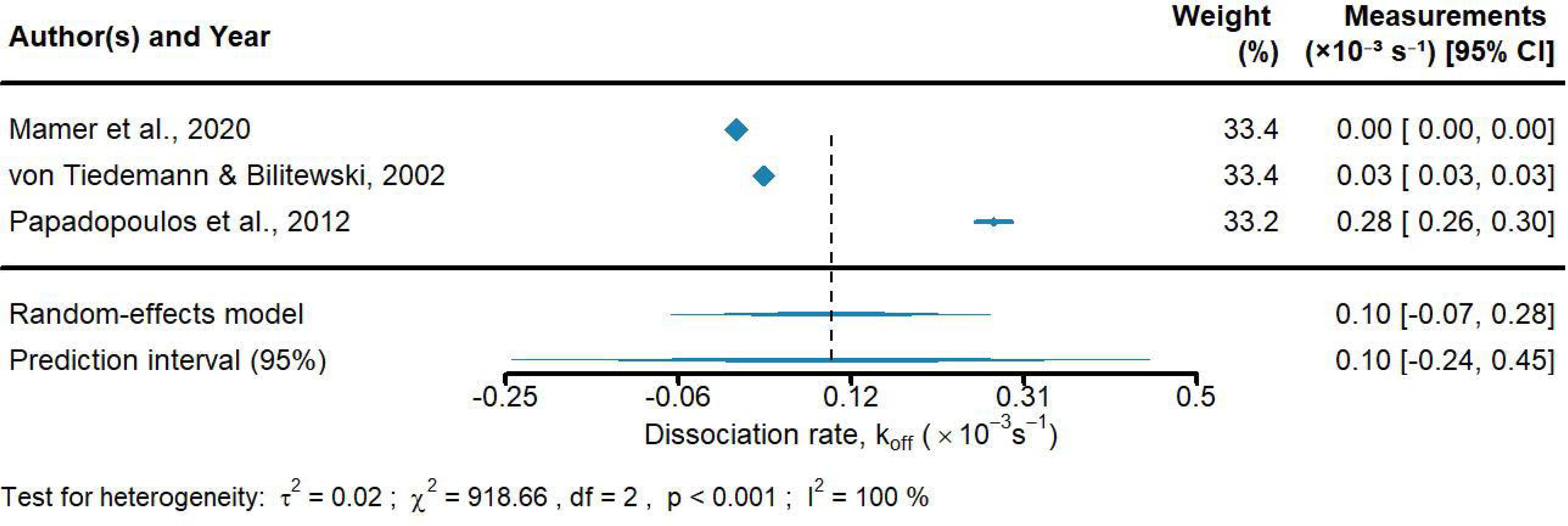
Forest plots of the analysis of dissociation rates, k_off_ (s^−1^) of VEGF to VEGFR1 across studies. The first, third, and last columns represent the list of references, their weights in the analysis, and measurements with a 95% confidence interval, respectively. The size of the blue diamond for each study represents its statistical weight in the analysis. The two diamonds below the table and the black dashed line show the combined measurements across studies. The upper diamond represents the mean and 95% confidence interval, and the lower diamond represents the mean and 95% prediction interval. The results of the heterogeneity test are represented by the estimate of between-studies variance (*τ*^2^), Cochran’s *Q*-test statistic for heterogeneity (*χ*^2^), degrees of freedom (df), p-value (p), and proportion of heterogeneity-induced variability among the total variability (I^2^).

**Figure S23.**
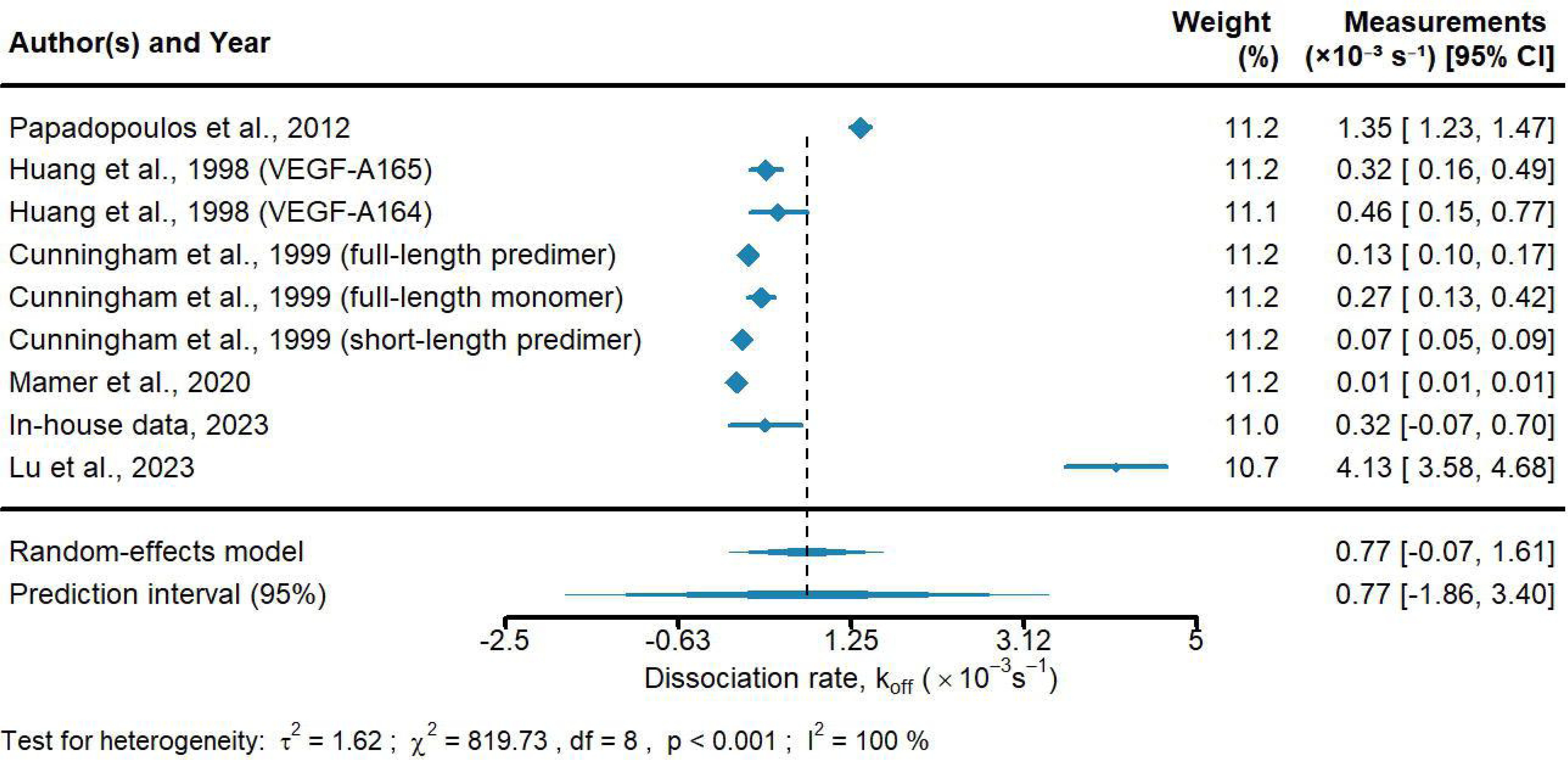
Forest plots of the analysis of dissociation rates, k_off_ (s^−1^) of VEGF to VEGFR2 across studies. The first, third, and last columns represent the list of references, their weights in the analysis, and measurements with a 95% confidence interval, respectively. The size of the blue diamond for each study represents its statistical weight in the analysis. The two diamonds below the table and the black dashed line show the combined measurements across studies. The upper diamond represents the mean and 95% confidence interval, and the lower diamond represents the mean and 95% prediction interval. The results of the heterogeneity test are represented by the estimate of between-studies variance (*τ*^2^), Cochran’s *Q*-test statistic for heterogeneity (*χ*^2^), degrees of freedom (df), p-value (p), and proportion of heterogeneity-induced variability among the total variability (I^2^).

**Figure S24.**
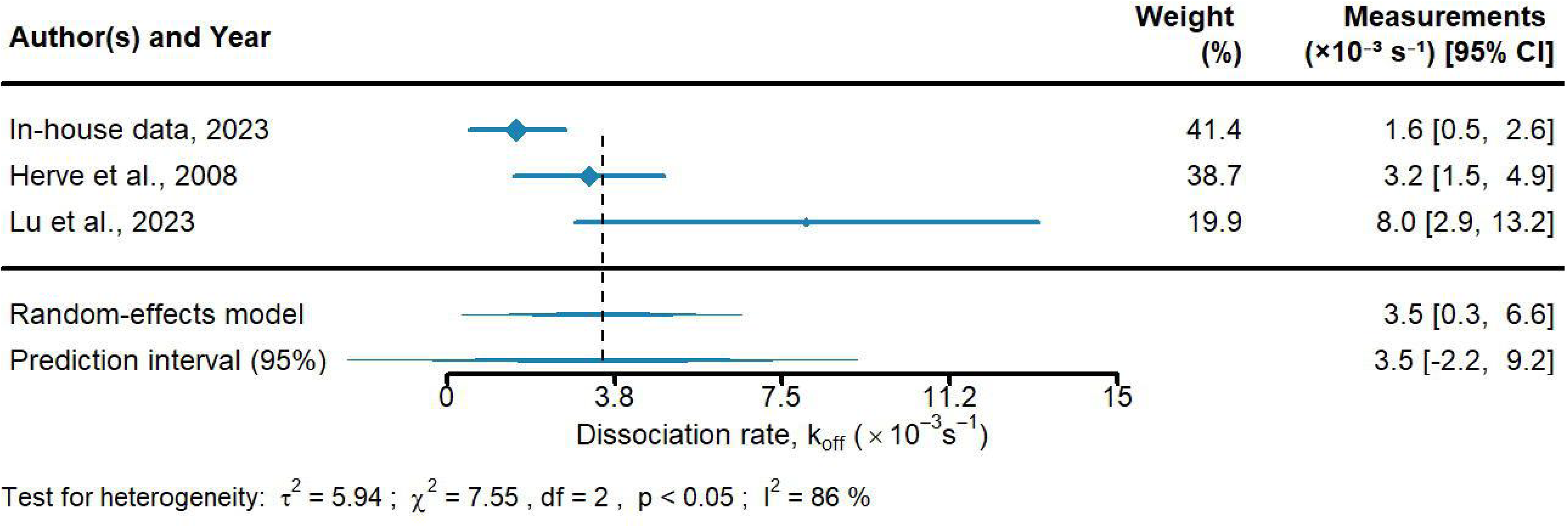
Forest plots of the analysis of dissociation rates, k_off_ (s^−1^) of VEGF to NRP1 across studies. The first, third, and last columns represent the list of references, their weights in the analysis, and measurements with a 95% confidence interval, respectively. The size of the blue diamond for each study represents its statistical weight in the analysis. The two diamonds below the table and the black dashed line show the combined measurements across studies. The upper diamond represents the mean and 95% confidence interval, and the lower diamond represents the mean and 95% prediction interval. The results of the heterogeneity test are represented by the estimate of between-studies variance (*τ*^2^), Cochran’s *Q*-test statistic for heterogeneity (*χ*^2^), degrees of freedom (df), p-value (p), and proportion of heterogeneity-induced variability among the total variability (I^2^).

**Figure S25.**
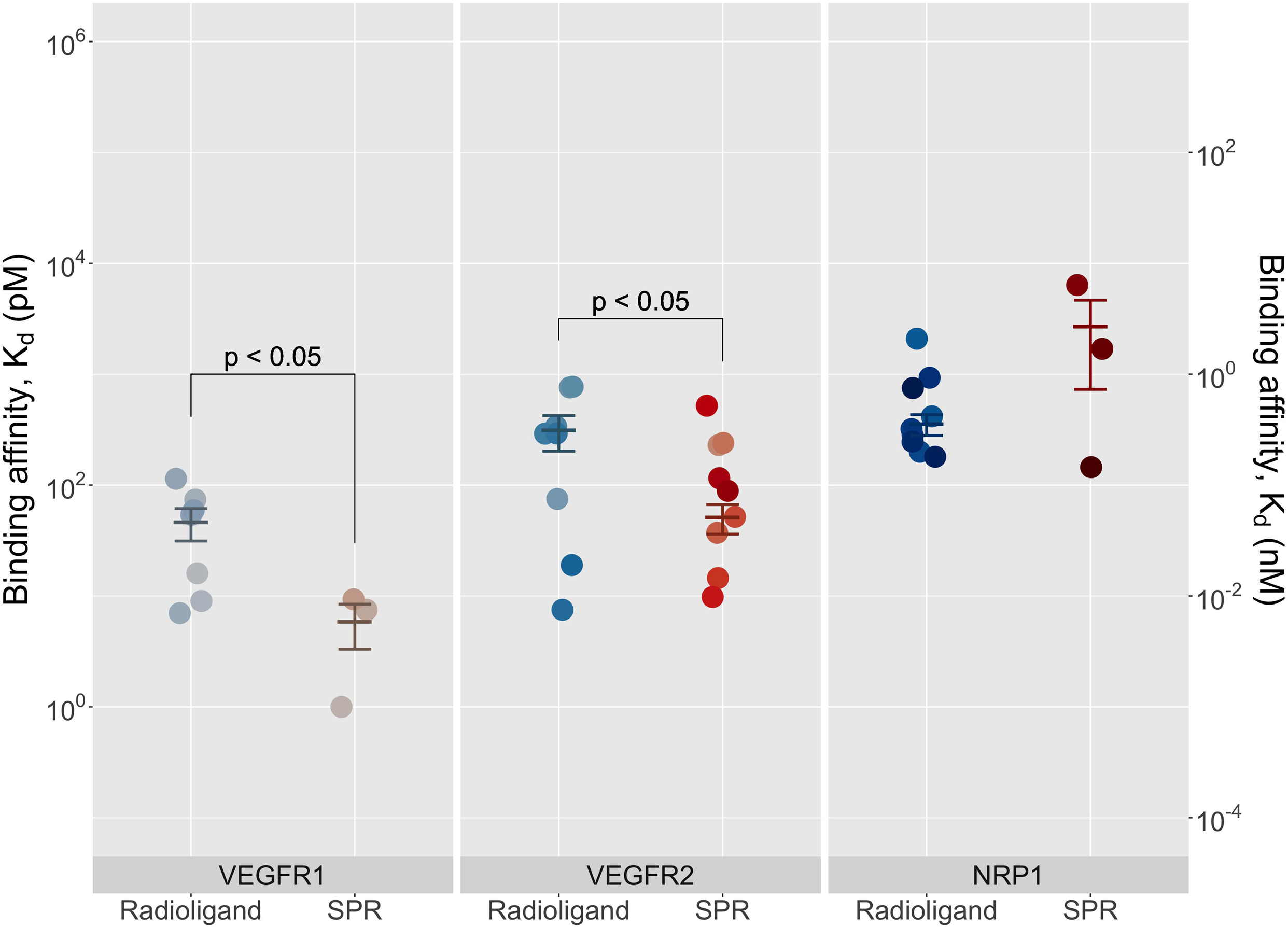
Binding affinities of VEGF to VEGFR1, VEGFR2, and NRP1 measured by cell-based (radioligand) assay and chip-based (surface plasmon resonance) assay. The error bar represents the weighted mean ± standard error of VEGF binding affinities for VEGFR1, VEGFR2, and NRP1. A one-tailed Welch’s t-test (p-value < 0.05) was used to compare a pair of groups.

**Figure S26.**
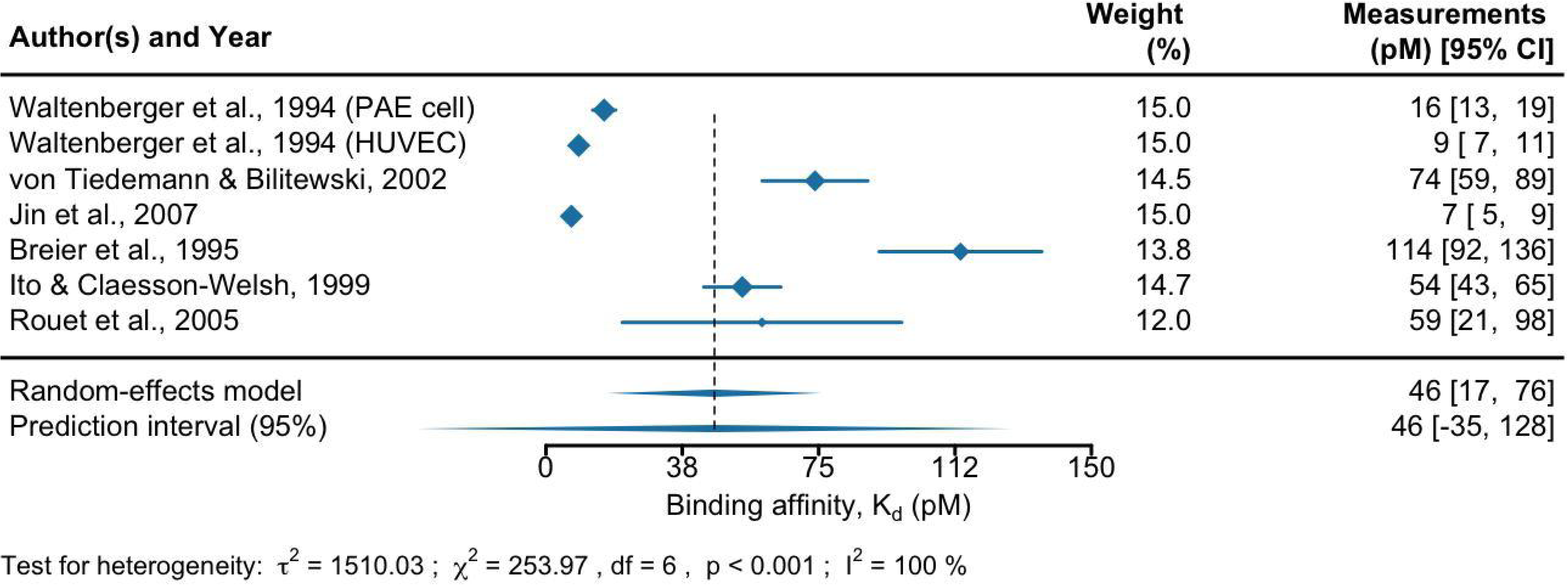
Forest plots of the analysis of binding affinity (pM) of VEGF to VEGFR1 measured by radioligand assay across studies. The first, third, and last columns represent the list of references, their weights in the analysis, and measurements with a 95% confidence interval, respectively. The size of the blue diamond for each study represents its statistical weight in the analysis. The two diamonds below the table and the black dashed line show the combined measurements across studies. The upper diamond represents the mean and 95% confidence interval, and the lower diamond represents the mean and 95% prediction interval. The results of the heterogeneity test are represented by the estimate of between-studies variance (*τ*^2^), Cochran’s *Q*-test statistic for heterogeneity (*χ*^2^), degrees of freedom (df), p-value (p), and proportion of heterogeneity-induced variability among the total variability (I^2^). Abbreviation: porcine aortic endothelial (PAE) and human umbilical vein endothelial cell (HUVEC).

**Figure S27.**
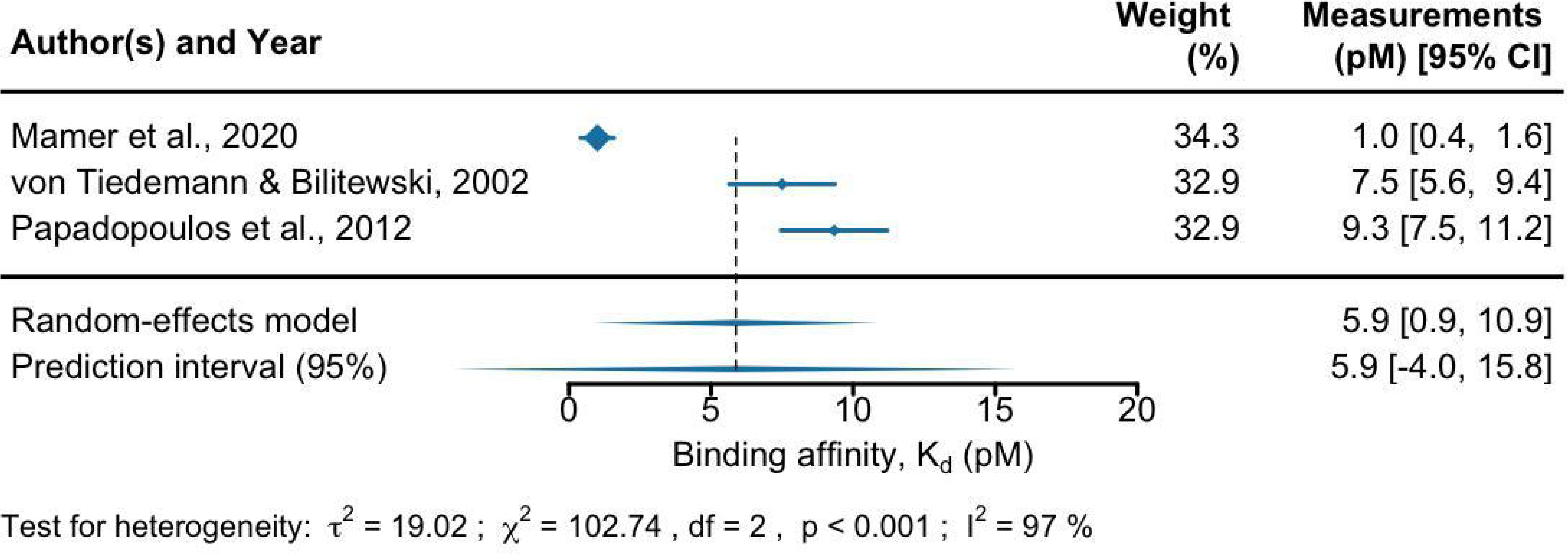
Forest plots of the analysis of binding affinity (pM) of VEGF to VEGFR1 measured by surface plasmon resonance across studies. The first, third, and last columns represent the list of references, their weights in the analysis, and measurements with a 95% confidence interval, respectively. The size of the blue diamond for each study represents its statistical weight in the analysis. The two diamonds below the table and the black dashed line show the combined measurements across studies. The upper diamond represents the mean and 95% confidence interval, and the lower diamond represents the mean and 95% prediction interval. The results of the heterogeneity test are represented by the estimate of between-studies variance (*τ*^2^), Cochran’s *Q*-test statistic for heterogeneity (*χ*^2^), degrees of freedom (df), p-value (p), and proportion of heterogeneity-induced variability among the total variability (I^2^).

**Figure S28.**
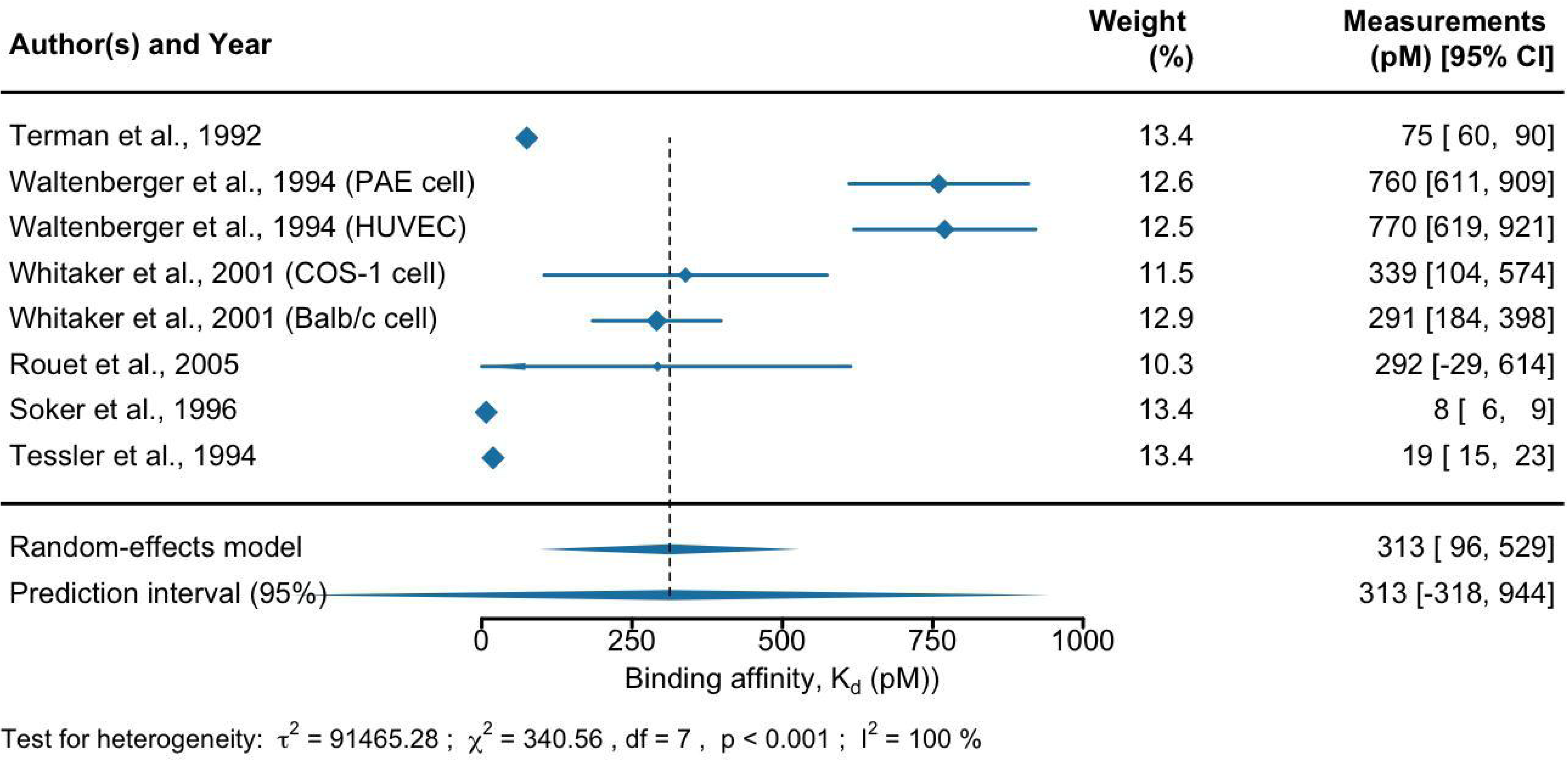
Forest plots of the analysis of binding affinity (pM) of VEGF to VEGFR2 measured by radioligand assay across studies. The first, third, and last columns represent the list of references, their weights in the analysis, and measurements with a 95% confidence interval, respectively. The size of the blue diamond for each study represents its statistical weight in the analysis. The two diamonds below the table and the black dashed line show the combined measurements across studies. The upper diamond represents the mean and 95% confidence interval, and the lower diamond represents the mean and 95% prediction interval. The results of the heterogeneity test are represented by the estimate of between-studies variance (*τ*^2^), Cochran’s *Q*-test statistic for heterogeneity (*χ*^2^), degrees of freedom (df), p-value (p), and proportion of heterogeneity-induced variability among the total variability (I^2^). Abbreviation: porcine aortic endothelial (PAE) and human umbilical vein endothelial cell (HUVEC).

**Figure S29.**
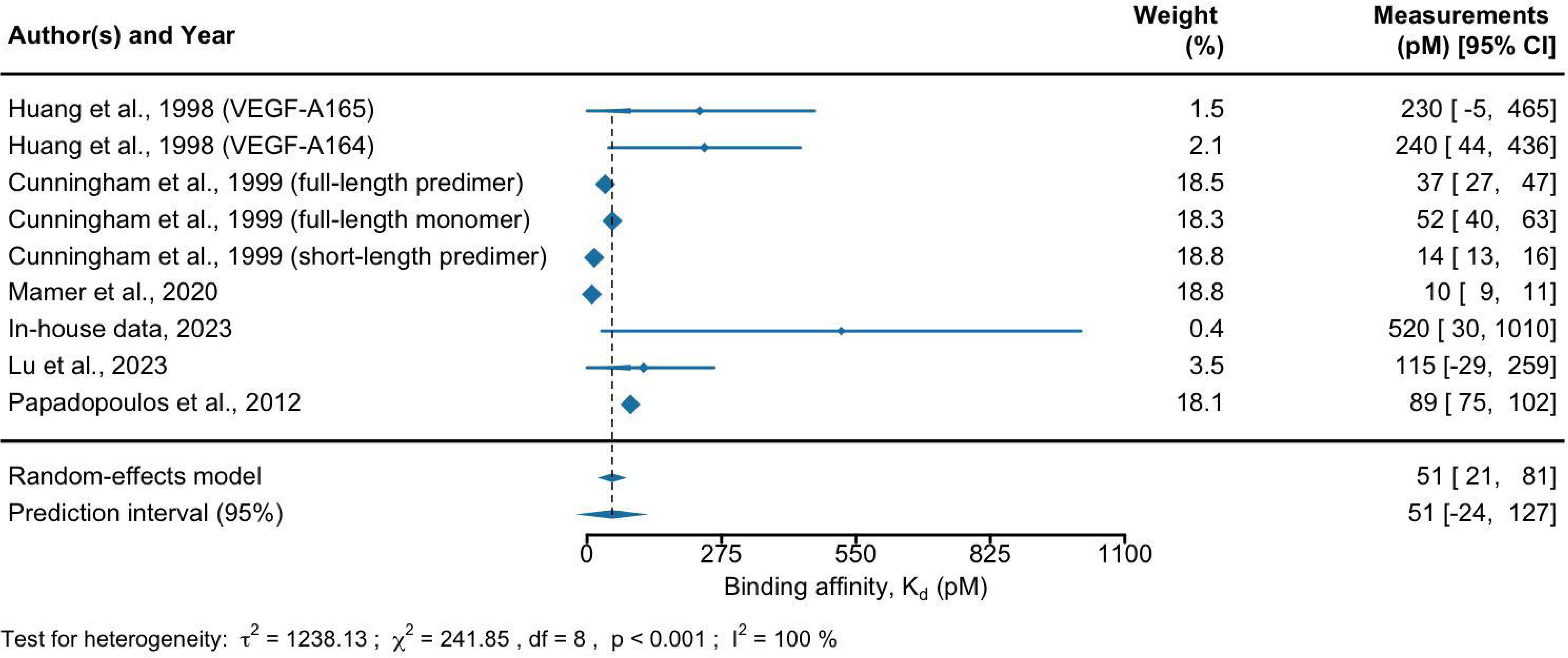
Forest plots of the analysis of binding affinity (pM) of VEGF to VEGFR2 measured by surface plasmon resonance across studies. The first, third, and last columns represent the list of references, their weights in the analysis, and measurements with a 95% confidence interval, respectively. The size of the blue diamond for each study represents its statistical weight in the analysis. The two diamonds below the table and the black dashed line show the combined measurements across studies. The upper diamond represents the mean and 95% confidence interval, and the lower diamond represents the mean and 95% prediction interval. The results of the heterogeneity test are represented by the estimate of between-studies variance (*τ*^2^), Cochran’s *Q*-test statistic for heterogeneity (*χ*^2^), degrees of freedom (df), p-value (p), and proportion of heterogeneity-induced variability among the total variability (I^2^).

**Figure S30.**
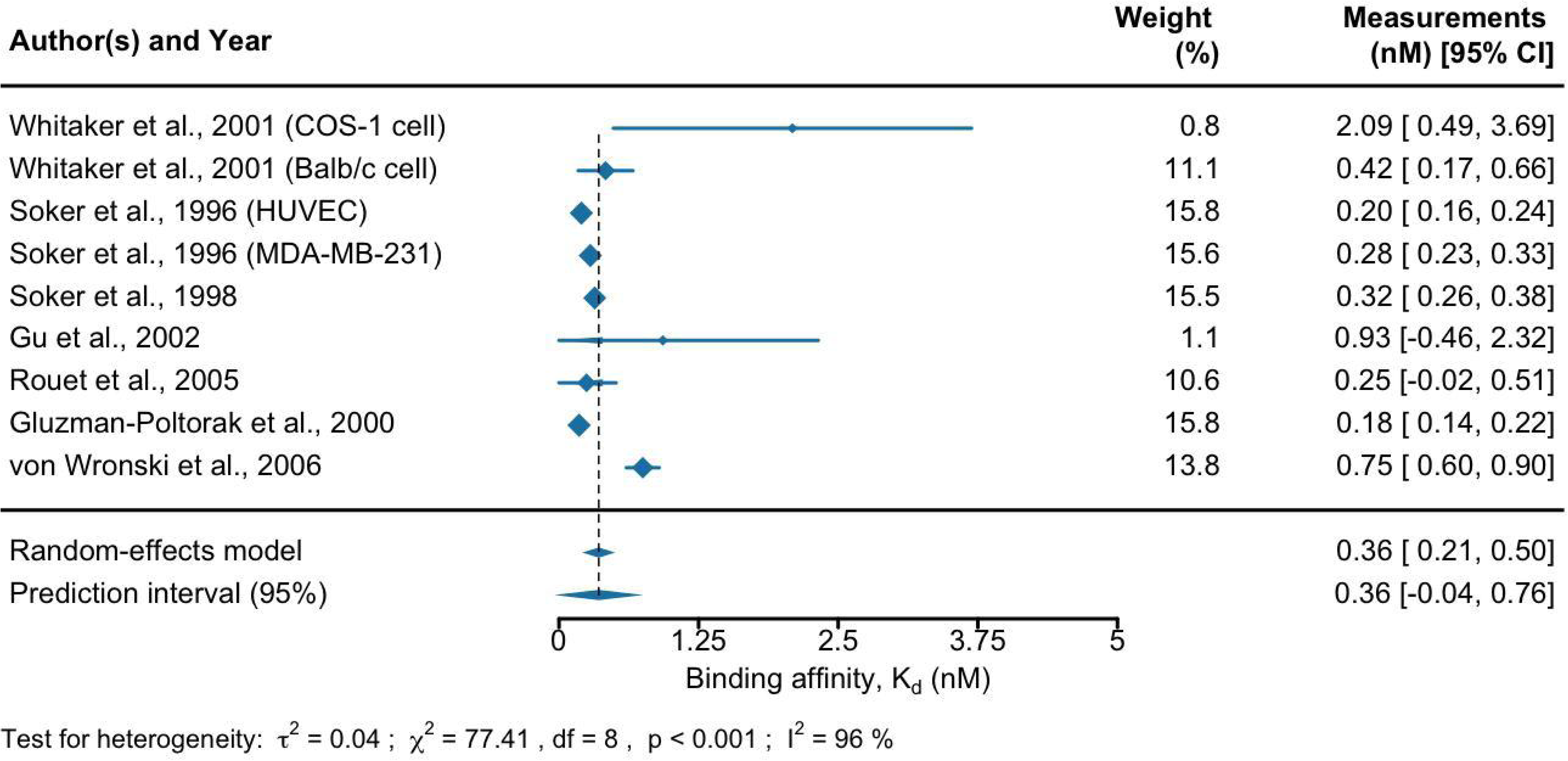
Forest plots of the analysis of binding affinity (nM) of VEGF to NRP1 measured by radioligand assay across studies. The first, third, and last columns represent the list of references, their weights in the analysis, and measurements with a 95% confidence interval, respectively. The size of the blue diamond for each study represents its statistical weight in the analysis. The two diamonds below the table and the black dashed line show the combined measurements across studies. The upper diamond represents the mean and 95% confidence interval, and the lower diamond represents the mean and 95% prediction interval. The results of the heterogeneity test are represented by the estimate of between-studies variance (*τ*^2^), Cochran’s *Q*-test statistic for heterogeneity (*χ*^2^), degrees of freedom (df), p-value (p), and proportion of heterogeneity-induced variability among the total variability (I^2^). Inside the parentheses of reference names, the cell lines used in the assay are included. Abbreviation: human umbilical vein endothelial cell (HUVEC).

**Figure S31.**
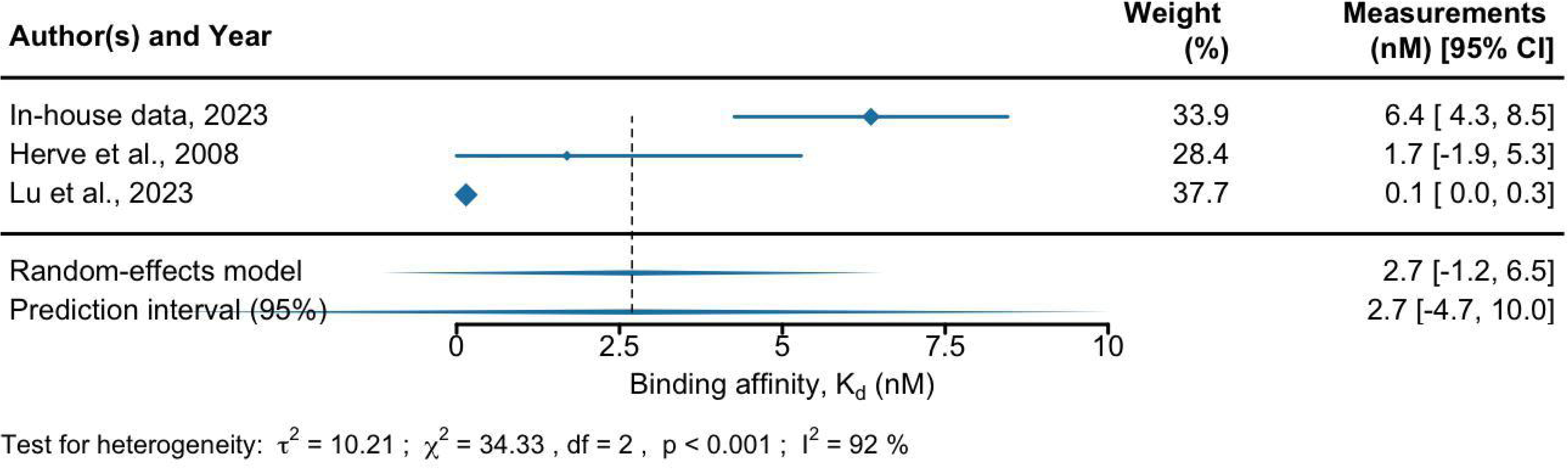
Forest plots of the analysis of binding affinity (nM) of VEGF to NRP1 measured by surface plasmon resonance across studies. The first, third, and last columns represent the list of references, their weights in the analysis, and measurements with a 95% confidence interval, respectively. The size of the blue diamond for each study represents its statistical weight in the analysis. The two diamonds below the table and the black dashed line show the combined measurements across studies. The upper diamond represents the mean and 95% confidence interval, and the lower diamond represents the mean and 95% prediction interval. The results of the heterogeneity test are represented by the estimate of between-studies variance (*τ*^2^), Cochran’s *Q*-test statistic for heterogeneity (*χ*^2^), degrees of freedom (df), p-value (p), and proportion of heterogeneity-induced variability among the total variability (I^2^).

