## Supplementary information for "Integrative analysis of angiogenic signaling in obesity: capillary features and VEGF binding kinetics"

### Data extraction

We used the following data extraction techniques from the main text or supplementary materials:

1. If the authors provided an average with the minimum and maximum values and the sample size, without standard error or deviation, then we estimated the standard error using the formula that depends on the sample size[1].
2. If the median, sample size, and 1st and 3rd quartiles were provided (e.g., in box plots), then we estimated the average and standard error using a method proposed by Wan et al., 2014[2].
3. If the standard error was reported in a transformed value (e.g., log-transformed value or adipocyte area reported instead of diameter), we estimated the normal standard error using the delta method[3]. To explain in more detail, let $Y$ be a function of $X$, $f(X)$, where $f$ is a transformation mapping $X$ to $Y$. Then, the variance of $Y$, $Var(Y)$, can be represented as follows:
$$Var(Y) = Var(X) \times [f'(\mu_x)]^2$$
where $f'(X)$ is the derivative of $f(X)$ and $\mu_x$ is the mean of $X$. Since $Var(X)$, $f(X)$, and $\mu_x$ are known, $Var(Y)$ can be calculated.
4. When the average and error bars were provided via a bar graph without reporting numbers, we used an image processing program, ImageJ V1.53k (https://imagej.net/), to measure them.
5. To extract capillary basement membrane thickness data from Fraselle-Jacobs et al.[4], we used ImageJ to measure the thickness at multiple points from Figure 3C and calculated the average and standard error.
6. If the papers reported VEGF binding affinity to its receptors via radioligand binding assays and did not provide the standard error, we assumed the standard error as 10% of the binding affinity. This assumption is based on the two studies: (1) the standard error of a binding affinity is usually less than 10% when the saturation assay is repeatedly performed[5], and (2) if the labeled and unlabeled ligands have the same binding affinities and the ligand depletion is less than 10%, then the expected error of binding affinities of unlabeled ligands is around 5%[6]. In our analysis, the studies employing radioligand assays used radiolabeled VEGF and unlabeled VEGF. Besides, researchers usually perform the assay with less than 10% ligand depletion. Thus, it is reasonable to assume the standard error to be 10% of the measurement value.
7. If a paper using SPR reported multiple binding affinities without standard errors for different ligand concentrations (e.g., Huang et al.[7]), we calculated the mean value and standard error across the various concentrations of analytes. This approach was grounded in the theory because, theoretically, the affinity constant does not depend on the concentration of the analyte in SPR due to the constant concentration of the immobilized receptor.
8. If a paper reported binding affinities obtained from the non-linear regression curve fitting and the Scatchard analysis, then we chose the one obtained from the Scatchard analysis. This is because, except for Whitaker et al.[8] and Gu et al.[9], all studies included in our analysis measured binding affinities using the Scatchard method.
9. When a paper reported multiple binding affinities under treatments with several heparin conditions, we included only measurements with no heparin.
10. When studies that included sensorgram graphs from SPR experiments did not provide association and dissociation rate constant data, the values of the graph were extracted using ImageJ. We assumed that VEGF-A binding to NRP1 follows 1:1 Langmuir model since other SPR studies used the model when they measured VEGF-A:NRP1 kinetic data[10–12]. Using the 1:1 Langmuir binding model, data were fitted with the following equations[13] to estimate the association and dissociation parameters:

$$R_t = \frac{R_{max}[A]}{\frac{k_{off}}{k_{on}}+[A]}\left[1 - e^{-(k_{on}[A]+k_{off})t}\right], \text{ association equation}$$

$$R_t = R_0 e^{-k_{off}t}, \text{ dissociation equation}$$

where $R_t$ is the response at time $t$, $R_{max}$ is the maximum response, $R_0$ is the response level at the beginning of the dissociation, *[A]* is the concentration of analyte (e.g., VEGF), and $k_{on}$, $k_{off}$ are the association and dissociation rates, respectively.

11. To extract the standard error from Papadopoulos et al.[14], we used the following equation derived from the Taylor expansion[15] and $|\mathrm{Cov}(X,Y)| \le \sigma_X\sigma_Y$:

$$\mathrm{Var}\left(\frac{X}{Y}\right) = \frac{\mathrm{Var}(X)}{\mu_Y^2} + \frac{\mu_X^2 Var(Y)}{\mu_Y^4} - \frac{2\mu_X \mathrm{Cov}(X,Y)}{\mu_Y^3} \le \frac{\mathrm{Var}(X)}{\mu_Y^2} + \frac{\mu_X^2 \mathrm{Var}(Y)}{\mu_Y^4} + \frac{2\mu_X\sigma_X\sigma_Y}{\mu_Y^3}$$

where $\mu_x$ and $\mu_Y$ are the mean values of random variables *X* and *Y*, $\sigma_x$ and $\sigma_y$ are the standard deviation of *X* and *Y*, respectively, and *Cov(X, Y)* is the covariance of *X* and *Y*.

12. To extract VEGF-A binding affinity to NRP1 from von Wronski et al.[16], we used the concentration of radiolabeled VEGF-A ([Radioligand] = 250 pM), the reported $IC_{50}$, and the following Cheng and Prussoff equation for the homologous competitive binding experiment[17]: $K_d = IC_{50} – [Radioligand]$.
