## Supplementary tables for "Integrative analysis of angiogenic signaling in obesity: capillary features and VEGF binding kinetics"

**Table S1. Studies that measured adipocyte diameter in mouse gonadal adipose tissue**

| Reference | Animal | Strain | Sex | n | Age | Diet | Duration of diet | Body weight | Location | Method | Mean ± SE[†] |
|---|---|---|---|---|---|---|---|---|---|---|---|
| 1 | Mouse | Swiss × 129Sv | Male | 15 | 20 wk | SFD | 15 wk | 38 ± 1.4 g | Gonadal fat | CIA[‡] | 45 ± 2.55 μm |
| | | | Male | 10 | | HFD | | 57 ± 1.4 g | | | 66 ± 1.8 μm |
| 2 | Mouse | C57BL/6 × 129SvJ | Both | ? | 22 wk | SFD | 17 wk | 27 ± 2.2 g | Gonadal fat | CIA | 49 ± 4.2 μm |
| | | | Both | ? | | HFD | | 39 ± 3.1 g | | | 80 ± 5.3 μm |
| | | | Both | ? | 37 wk | HFD | 32 wk | 39 ± 2.2 g | | | 86 ± 1.6 μm |
| 3 | Mouse | C57BL/6 × 129SvJ | Male | 2 | 20 wk | SFD | 15 wk | 28 ± 1.2 g | Gonadal fat | CIA | 42 ± ? μm[§] |
| | | | Male | 6 | | HFD | | 40 ± 1.4 g | | | 83 ± 3 μm |
| 4 | Mouse | C57BL/6 × 129Sv | Both | 7–11 | 21 wk | SFD | 17 wk | 28 ± 1.4 g | Gonadal fat | CIA | 49 ± 4.3 μm |
| | | | Both | 7–11 | | HFD | | 42 ± 2 g | | | 82 ± 3.5 μm |
| 5 | Mouse | C57BL/6 | Male | ? | 7 wk | SFD | 2 wk | 21.1 ± 0.7 g | Gonadal fat | CIA | 21 ± 0.83 μm |
| | | | Male | ? | | HFD | | 26.4 ± 0.6 g | | | 35 ± 0.72 μm |
| | | | Male | ? | 10 wk | SFD | 5 wk | 25.4 ± 0.4 g | | | 27 ± 0.47 μm |
| | | | Male | ? | | HFD | | 32.9 ± 0.9 g | | | 41 ± 2 μm |
| | | | Male | ? | 20 wk | SFD | 15 wk | 30.0 ± 0.62 g | | | 29 ± 1.1 μm |
| | | | Male | ? | | HFD | | 46.3 ± 1.77 g | | | 53 ± 0.46 μm |
| 6 | Mouse | C57BL/6 | Male | 12–20 | 20 wk | SFD | 15 wk | 33 ± 0.91 g | Gonadal fat | CIA | 62 ± 4.1 μm |
| | | | Male | 12–20 | | HFD | | 45 ± 1.4 g | | | 85 ± 2.3 μm |
| 7 | Mouse | C57BL/6 × 129 SvJae | Male | 4 | 20 wk | SFD | 15 wk | 29.4 ± 0.6 g | Gonadal fat | CIA | 40.1 ± 0.763 μm |
| | | | Male | 10 | | HFD | | 41 ± 1.8 g | | | 94.6 ± 4.58 μm |
| 8 | Mouse | B10.RIII | Male | 11 | 20 wk | HFD | 15 wk | 37 ± 1.5 g | Gonadal fat | CIA | 76.4 ± 2.25 μm |
| 9 | Mouse | C57BL/6 | Male | 8 | 20 wk | HFD | 15 wk | 42 ± 1.4 g | Gonadal fat | CIA | 89.1 ± 1.46 μm |
| 10 | Mouse | C57BL6/129 SvJ/EMS + Ter | Male | 10–14 | 20 wk | SFD | 15 wk | 23 ± 0.46 g | Gonadal fat | CIA | 42.4 ± 1.952 μm |
| | | | | 10–14 | | HFD | | 27 ± 0.72 g | | | 58.37 ± 2.225 μm |

[†]SE: Standard error
[‡]CIA: Computer-assisted image analysis
[§]This data was not included in the analysis because of the missing standard error.

**Table S2. Studies that measured vessel size in mouse gonadal adipose tissue**

| Reference | Animal | Strain | Sex | n | Age | Diet | Duration of diet | Body weight | Location | Method | Mean ± SE |
|---|---|---|---|---|---|---|---|---|---|---|---|
| 1 | Mouse | Swiss × 129Sv | Male | 15 | 20 wk | SFD | 15 wk | 38 ± 1.4 g | Gonadal fat | CIA | 27 ± 1.7 μm² |
| | | | Male | 10 | | HFD | | 57 ± 1.4 g | | | 41 ± 3.1 μm² |
| 5 | Mouse | C57BL/6 | Male | 5 | 7 wk | SFD | 2 wk | 21.1 ± 0.7 g | Gonadal fat | CIA | 52 ± 1.9 μm² |
| | | | Male | 5 | | HFD | | 26.4 ± 0.6 g | | | 48 ± 2.2 μm² |
| | | | Male | 5 | 10 wk | SFD | 5 wk | 25.4 ± 0.4 g | | | 50 ± 2.6 μm² |
| | | | Male | 5 | | HFD | | 32.9 ± 0.9 g | | | 41 ± 1.8 μm² |
| | | | Male | 5 | 20 wk | SFD | 15 wk | 30.0 ± 0.62 g | | | 49 ± 3.4 μm² |
| | | | Male | 5 | | HFD | | 46.3 ± 1.77 g | | | 54 ± 3.3 μm² |
| 6 | Mouse | C57BL/6 | Male | 5–10 | 20 wk | SFD | 15 wk | 33 ± 0.91 g | Gonadal fat | CIA | 74 ± 4.8 μm² |
| | | | Male | 5–10 | | HFD | | 45 ± 1.4 g | | | 140 ± 19 μm² |
| 7 | Mouse | C57BL/6 × 129 SvJae | Male | 4 | 20 wk | SFD | 15 wk | 29.4 ± 0.6 g | Gonadal fat | CIA | ? |
| | | | Male | 10 | | HFD | | 41 ± 1.8 g | | | 47 ± 2.6 μm² |
| 8 | Mouse | B10.RIII | Male | 7–11 | 20 wk | HFD | 15 wk | 37 ± 1.5 g | Gonadal fat | CIA | 76 ± 3.9 μm² |
| 9 | Mouse | C57BL/6 | Male | 8 | 20 wk | HFD | 15 wk | 42 ± 1.4 g | Gonadal fat | CIA | 108 ± 7.7 μm² |
| 10 | Mouse | C57BL6/129 SvJ/EMS + Ter | Male | 10–14 | 20 wk | SFD | 15 wk | 23 ± 0.46 g | Gonadal fat | CIA | 59 ± 5.1 μm² |
| | | | | 10–14 | | HFD | | 27 ± 0.72 g | | | 49 ± 2.8 μm² |

**Table S3. Studies that measured vessel size in mouse tumors**

| Reference | Animal | Strain | Sex | n | Age | Body weight | Tumor cell source | Injected location | Staining antibody | Measurement method | Mean ± SE |
|---|---|---|---|---|---|---|---|---|---|---|---|
| 11 | Mouse | C3H/HeNCr | Female | 4 | 9–11 wk | ? | Radiation-induced fibrosarcoma (RIF) | Shoulder | CD31 | Immunohisto-chemistry (fluorescence microscopy) | 53 ± 9.5 μm$^2$† |
| | | Nude | | 5 | | | | | | | 130 ± 15 μm$^2$† |
| 12 | Mouse | Nude | Female | 6 | 6–8 wk | ? | 4T1 primary tumor + 5% dextrose (control) | Left third mammary fat pad | CD31 | Confocal microscopy | 86.7 ± 2.76 μm$^2$† |
| | | | | 6 | | | 4T1 primary tumor + AKB-9778 | | | | 110 ± 3.71 μm$^2$ |
| | | C57BL/6J | | 6 | | | E0771 tumors + 5% dextrose (control) | | | | 88.9 ± 1.11 μm$^2$† |
| | | | | 6 | | | E0771 tumors + AKB-9778 | | | | 107 ± 3.67 μm$^2$ |
| | | Nos3-/- | | 5 | | | E0771 tumors + 5% dextrose (control) | | | | 75.2 ± 1.23 μm$^2$ |
| | | | | 5 | | | E0771 tumors + AKB-9778 | | | | 71.4 ± 2.39 μm$^2$ |
| 13 | Mouse | Balb/c nu/nu | ? | 9 | 8–12 wk | ? | Human melanoma xenograft D-12 | Flank | Hemato-xylin and eosin | CIA | 135 ± 8.32 μm$^2$† |
| | | | | 9 | | | R-18 | | | | 93.3 ± 4.23 μm$^2$† |
| | | | | 9 | | | U-25 | | | | 113 ± 5.51 μm$^2$† |
| 14 | Mouse | C57BL/6 | ? | ? | ? | ? | B16F1 murine melanoma cells | Right flank | Endo-mucin | CIA | 216.46 ± 19.31 μm$^2$† |
| | | | | 17 | | | | | | | 209.62 ± 15.46 μm$^2$† |
| 15 | Mouse | Balb/c nu/nu | ? | ? | 10 wk | 24–27 g | C51 murine colon carcinoma | Left flank | - | SRμCT‡ in phase contrast mode | 45 ± 36 μm$^2$ (mean ± SD) |
| | | | | ? | | | | | | SRμCT in absorption contrast mode | 12 ± 6.7 μm$^2$ (mean ± SD) |
| 16 | Mouse | NOD/SCID | Female | ? | 7–10 wk | 20–25 g | MDA-MB-231 human breast cancer cells | Fat pad of mammary crest in the groin area | CD31 | CIA | 42 ± 19 μm$^2$ (mean ± SD) |
| | | | | ? | | | BT-474 human breast cancer cells | | | | 113 ± 115 μm$^2$ (mean ± SD) |
| 17 | Mouse | C57BL/6 | Female | 3 | 8–12 wk | ? | Lewis lung carcinoma (LLC) | Right flank | CD31 | CIA | 1128 ± 46.73 μm$^2$ |

†Data were included in the analysis.
‡Synchrotron radiation–based micro computed tomography (SRμCT)

**Table S4. Studies that measured vessel density in mouse gonadal adipose tissue**

| Reference | Animal | Strain | Sex | n | Age | Diet | Duration of diet | Body weight | Location | Method | Mean ± SE |
|---|---|---|---|---|---|---|---|---|---|---|---|
| [1] | Mouse | Swiss × 129Sv | Male | 15 | 20 wk | SFD | 15 wk | 38 ± 1.4 g | Gonadal fat | CIA | 370 ± 37/mm$^2$ |
| | | | Male | 10 | | HFD | | 57 ± 1.4 g | | | 290 ± 23/mm$^2$ |
| [5] | Mouse | C57BL/6 | Male | 5 | 7 wk | SFD | 2 wk | 21.1 ± 0.7 g | Gonadal fat | CIA | 1200 ± 55/mm$^2$ |
| | | | Male | 5 | | HFD | | 26.4 ± 0.6 g | | | 790 ± 30/mm$^2$ |
| | | | Male | 5 | 10 wk | SFD | 5 wk | 25.4 ± 0.4 g | | | 850 ± 40/mm$^2$ |
| | | | Male | 5 | | HFD | | 32.9 ± 0.9 g | | | 410 ± 20/mm$^2$ |
| | | | Male | 5 | 20 wk | SFD | 15 wk | 30.0 ± 0.62 g | | | 790 ± 41/mm$^2$ |
| | | | Male | 5 | | HFD | | 46.3 ± 1.77 g | | | 490 ± 19/mm$^2$ |
| | | Wild-type littermate of ob/ob | Male | 5 | ? | SFD | ? | 27 ± 0.62 g | | | 830 ± 87/mm$^2$ |
| | | ob/ob | Male | 5 | 9 wk | SFD | ? | 41 ± 1.2 g | | | 390 ± 21/mm$^2$† |
| [6] | Mouse | C57BL/6 | Male | 5–10 | 20 wk | SFD | 15 wk | 33 ± 0.91 g | Gonadal fat | CIA | 280 ± 56/mm$^2$ |
| | | | Male | 5–10 | | HFD | | 45 ± 1.4 g | | | 200 ± 34/mm$^2$ |
| [7] | Mouse | C57BL/6 × 129 SvJae | Male | 4 | 20 wk | SFD | 15 wk | 29.4 ± 0.6 g | Gonadal fat | CIA | ? |
| | | | Male | 10 | | HFD | | 41 ± 1.8 g | | | 120 ± 6.2/mm$^2$ |
| [8] | Mouse | B10.RIII | Male | 7–11 | 20 wk | HFD | 15 wk | 37 ± 1.5 g | Gonadal fat | CIA | 210 ± 17/mm$^2$ |
| [9] | Mouse | C57BL/6 | Male | 8 | 20 wk | HFD | 15 wk | 42 ± 1.4 g | Gonadal fat | CIA | 238 ± 16/mm$^2$ |
| [10] | Mouse | C57BL6/129SvJ/EMS + Ter | Male | 10–14 | 20 wk | SFD | 15 wk | 23 ± 0.46 g | Gonadal fat | CIA | 740 ± 96/mm$^2$ |
| | | | | 10–14 | | HFD | | 27 ± 0.72 g | | | 400 ± 55/mm$^2$ |

†Data was excluded because the mouse strain is not a wild type.

**Table S5. Studies that measured vessel density in mouse tumors**

| Reference | Animal | Strain | Sex | n | Age | Body weight | Tumor cell source | Injected Location | Staining antibody | Measurement Method | Mean ± SE |
|---|---|---|---|---|---|---|---|---|---|---|---|
| 17 | Mouse | C57BL/6 | Female | 3 | 8–12 wk | ? | Lewis lung carcinoma (LLC) | Right flank | CD31 | CIA | $125.8 \pm 5.2/mm^{2\dagger}$ |
| 12 | Mouse | Nude | Female | 6 | 6–8 wk | ? | 4T1 primary tumor + 5% dextrose (control) | Left third mammary fat pad | CD31 | Confocal microscopy | $292 \pm 28.6/mm^{2\ddagger}$ |
| | | | | 6 | | | 4T1 primary tumor + AKB-9778 | | | | $400 \pm 55.5/mm^2$ |
| | | C57BL/6J | | 6 | | | E0771 tumors + 5% dextrose (control) | | | | $212 \pm 25.6/mm^{2\ddagger}$ |
| | | | | 6 | | | E0771 tumors + AKB-9778 | | | | $269 \pm 15.4/mm^2$ |
| | | Nos3-/- | | 5 | | | E0771 tumors + 5% dextrose (control) | | | | $95.8 \pm 19.1/mm^2$ |
| | | | | 5 | | | E0771 tumors + AKB-9778 | | | | $82.7 \pm 10.4/mm^2$ |
| 13 | Mouse | Balb/c nu/nu | ? | 5 | 8–12 wk | ? | Human melanoma xenograft: D-12 | Flank | Hema-toxylin and eosin | CIA | $19.5 \pm 2.1/mm^{2\ddagger}$ |
| | | | | 5 | | | R-18 | | | | $19.1 \pm 1.8/mm^{2\ddagger}$ |
| | | | | 5 | | | U-25 | | | | $17.6 \pm 2.4/mm^{2\ddagger}$ |
| 14 | Mouse | C57BL/6 | ? | ? | ? | ? | B16F1 murine melanoma cells | Right flank | Endo-mucin | CIA | $70.62 \pm 9.74/mm^{2\ddagger}$ |
| | | | | 17 | | | | | | | $48.87 \pm 4.75/mm^{2\ddagger}$ |
| 16 | Mouse | NOD/SCID | Female | ? | 7–10 wk | 20–25 g | MDA-MB-231 human breast cancer cells | Fat pad of mammary crest in the groin area | CD31 | CIA | $13.5 \pm 1.6/mm^2$ (mean ± SD) |
| | | | | ? | | | BT-474 human breast cancer cells | | | | $11.1 \pm 2.2/mm^2$ (mean ± SD) |
| 18 | Mouse | Athymic nude mice | Female | ? | 4–5 wk | ? | U87MG human glioblastoma cells: Small tumor | Front leg | CD31 | CIA | $96 \pm 19/mm^2$ (mean ± SD) |
| | | | | ? | | | Large tumor | | | | $20 \pm 9/mm^2$ (mean ± SD) |
| 19 | Mouse | B6;129 | ? | 30 | ? | ? | B16F0 melanoma cells | Flank | Endo-mucin | Immunohisto-chemical analysis | $31.87 \pm 5.8/mm^{2\ddagger}$ |
| | | | | 20 | | | CMT19T lung carcinoma cells | | | | $29.65 \pm 2.81/mm^{2\ddagger}$ |
| | | | | 9 | | | B16F0 melanoma cells (injected DMSO as a control of FAK inhibitor) | | | | $64.85 \pm 7/mm^{2\ddagger}$ |
| 20 | Mouse | C57BL/6 | Male | 4 | 7–9 wk | ? | Luciferase-tagged Lewis lung carcinoma cells | ? | CD31 | CIA | $147.74 \pm 17.9/mm^{2\ddagger}$ |
| | | C57BL/6-based Rpl29- | | 4 | | | | | | | $94.33 \pm 7.75/mm^2$ |

| | | |
|---|---|---|
| | | heterozygote mice |

†This paper was excluded; because of the too large vessel size, we considered it an outlier.
‡Data were included in the analysis.

**Table S6. Studies that measured capillary basement membrane thickness in murine tissues**

| Reference | Animal | Age | n | Status | Body weight | Location | Method | Mean ± SD† |
|---|---|---|---|---|---|---|---|---|
| 21 | Male CBA mouse | 1.5 mo | 4 | Normal | ? | Retina (mid-zone) | TEM‡ | 50 ± 9 nm§ |
| | | 4 mo | 4 | | | | | 59 ± 13 nm§ |
| | | 8 mo | 4 | | | | | 75 ± 10 nm§ |
| | | 12 mo | 4 | | | | | 108 ± 17 nm§ |
| | | 20 mo | 4 | | | | | 154 ± 27 nm§ |
| | Female Balb/c | 1.5 mo | 2 | | | | | 55 ± 9 nm§ |
| | | 8 mo | 2 | | | | | 77 ± 9 nm§ |
| | | 20 mo | 2 | | | | | 158 ± 35 nm§ |
| | | 1.5 mo | 2 | | | Retina (center) | | 41 ± 4 nm§ |
| | | 1.5 mo | 2 | | | Retina (periphery) | | 64 ± 6 nm§ |
| 22 | Male SJL/J mouse | 5-6 wk+ 6 mo (7 mo) | ? | Normal (Uninfected) | ? | Retina | TEM | 69 ± 5.5 nm |
| | | | 8 | Diabetes (EMC virus infected) | | | | 91 ± 10.2 nm |
| | | | 12 | Normal (Uninfected) | | Kidney | | 91 ± 3 nm§ |
| | | | 12 | Diabetes (EMC virus infected) | | | | 382 ± 6 nm |
| 23 | C57BL/6J mouse | 6 mo | 100 | Normal | ? | Brain | TEM | 56.78 ± 12.50 nm§ |
| | | 24 mo | 100 | Normal | ? | | | 107.53 ± 23.76 nm§ |
| 24 | Spiny mouse | 220±146 days (7±5 mo) | 8 | Normal | 45 ± 10 g | Gastrocnemius (muscle) | EM¶ | 73 ± 16 nm§ |
| | | 254±190 days (8.5±6.3 mo) | 6 | Moderately impaired glucose tolerance | 49 ± 7 g | | | 75 ± 18 nm |
| | | 207±125 days (7±4 mo) | 6 | Severely impaired glucose tolerance | 50 ± 11 g | | | 80 ± 18 nm |
| | | 462±174 days (15.4±6 mo) | 5 | Spontaneous severe diabetes | ? | | | 105 ± 9 nm |

| | | | | | | | | |
|---|---|---|---|---|---|---|---|---|
| 25 | Control (FVB/N) vs. OVE26 mouse | 300–350 days (10–12 mo) | 8 | Normal | 39.40 ± 3.11 g | Retina | TEM | 92.87 ± 18.90 nm§ |
| | | | 14 | Genetic diabetes | 40.52 ± 3.16 g | | | 113.09 ± 9.57 nm |
| | | | 10 | Normal | 39.40 ± 3.11 g | Extensor digitorum (muscle) | | 76.75 ± 14.17 nm§ |
| | | | 8 | Genetic diabetes | 40.52 ± 3.16 g | | | 72.10 ± 16.85 nm |
| | | | 19 | Normal | 39.40 ± 3.11 g | Kidney | | 178.16 ± 35.61 nm§ |
| | | | 12 | Genetic diabetes | 40.52 ± 3.16 g | | | 333.19 ± 66.07 nm |
| | | | 8 | Normal | 39.40 ± 3.11 g | Pulmonary alveolus | | 67.99 ± 6.95 nm |
| | | | 8 | Genetic diabetes | 40.52 ± 3.16 g | | | 77.02 ± 8.97 nm |
| | | | 8 | Normal | 39.40 ± 3.11 g | Diaphragm (muscle) | | 54.75 ± 5.00 nm§ |
| | | | 8 | Genetic diabetes | 40.52 ± 3.16 g | | | 67.98 ± 2.81 nm |
| | | | 8 | Normal | 39.40 ± 3.11 g | Pancreas | | 103.54 ± 16.33 nm |
| | | | 6 | Genetic diabetes | 40.52 ± 3.16 g | | | 120.77 ± 28.05 nm |
| | | | 3 | Normal | 39.40 ± 3.11 g | Choroid | | 78.58 ± 7.06 nm |
| | | | 5 | Genetic diabetes | 40.52 ± 3.16 g | | | 81.73 ± 12.71 nm |
| | | | 12 | Normal | 39.40 ± 3.11 g | Heart IVS# | | 63.13 ± 5.91 nm§ |
| | | | 13 | Genetic diabetes | 40.52 ± 3.16 g | | | 69.85 ± 12.34 nm |
| | | | 12 | Normal | 39.40 ± 3.11 g | Heart LV†† | | 56.92 ± 4.02 nm§ |
| | | | 12 | Genetic diabetes | 40.52 ± 3.16 g | | | 62.97 ± 9.64 nm |
| | | | 15 | Normal | 39.40 ± 3.11 g | Peripheral nerve | | 58.75 ± 9.86 nm |
| | | | 5 | Genetic diabetes | 40.52 ± 3.16 g | | | 58.64 ± 4.55 nm |
| 26 | Male C57BL/6 mouse | 22 wk (5.5 mo) | 31 | Lean | 28.676 g | Skeletal muscle (gastrocnemius) | EM | 107.6 ± 5.482 nm§ (mean ± SE) |
| | | | 31 | Obese | 43.588 g | | | 102.7 ± 6.44 nm§ (mean ± SE) |
| 27 | FVB/NJ vs. Akita FVB/NJ mouse | 30 wk (7.5 mo) | 4 | Normal | ? | Kidney | TEM | 224.2 ± 27.7 nm§ (mean ± SE) |
| | | | 4 | Genetic diabetes (Akita) | ? | | | 240.8 ± 77.5 nm (mean ± SE) |
| 28 | FVB mouse | 350 days (12 mo) | 7 | Normal | 23.90 ± 0.52 g | Heart LV | TEM | 48.50 ± 7.31 nm§ |
| | Mt (overexpressing metallothionein) | | 4 | Normal & overexpressing MT | 25.60 ± 1.82 g | | | 44.56 ± 3.62 nm |
| | OVE26 | | 8 | Genetic diabetes | 23.60 ± 0.95 g | | | 62.18 ± 10.34 nm |
| | OVEMt (OVE26 + Mt) | | 7 | Genetic diabetes & overexpressing MT | 23.30 ± 1.14 g | | | 47.28 ± 7.07 nm |
| 29 | Male and female C57BL/6 | 2 mo | 3 | Normal | ? | Brain (Striatum) | EM | 67.03 ± 4.34 nm§ (mean ± SE) |
| | | | 3 | | | Brain (Cortex) | | 65.79 ± 5.56 nm§ (mean ± SE) |

| | | | | | | | | |
|---|---|---|---|---|---|---|---|---|
| | | | 3 | | | Brain (Hippocampus) | | 62.58 ± 2.70 nm§ (mean ± SE) |
| | | | 3 | | | Brain (Thalamus) | | 70.11 ± 3.63 nm§ (mean ± SE) |
| | | 7 mo | 3 | | | Brain (Striatum) | | 74.94 ± 6.62 nm§ (mean ± SE) |
| | | | 3 | | | Brain (Cortex) | | 57.35 ± 4.33 nm§ (mean ± SE) |
| | | | 3 | | | Brain (Hippocampus) | | 63.50 ± 4.78 nm§ (mean ± SE) |
| | | | 3 | | | Brain (Thalamus) | | 58.62 ± 5.17 nm§ (mean ± SE) |
| | | 23 mo | 3 | | | Brain (Striatum) | | 74.23 ± 2.80 nm§ (mean ± SE) |
| | | | 3 | | | Brain (Cortex) | | 97.49 ± 12.27 nm§ (mean ± SE) |
| | | | 3 | | | Brain (Hippocampus) | | 109.50 ± 11.93 nm§ (mean ± SE) |
| | | | 3 | | | Brain (Thalamus) | | 108.43 ± 14.51 nm§ (mean ± SE) |
| 30 | Male db/m mouse | 22 wk (5.5 mo) | ? | Normal | ? | Retina (outer plexiform layer) | EM | 79.4 ± 1.9 nm§ (mean ± SE) |
| | Male db/db mouse | | ? | Diabetes | ? | | | 87.3 ± 3.4 nm (mean ± SE) |
| | | | ? | Diabetes + A717 | ? | | | 83.8 ± 1.8 nm (mean ± SE) |
| 31 | Zucker rat | 11 wk (2.75 mo) | 6 | Lean (FA/fa) | ? | Plantar muscle | EM | 61.87 ± 1.33 nm§ (Mean ± SE) |
| | | | 6 | Genetic obesity (fa/fa) | | | | 68.13 ± 1.66 nm§ (Mean ± SE) |
| | | | 6 | Genetic obesity + exercise (6–11 wk) | | | | 65.10 ± 1.43 nm (Mean ± SE) |
| | | 18 wk (4.5 mo) | 6 | Lean (Fa/fa) | | | | 55.67 ± 1.04 nm§ (Mean ± SE) |
| | | | 7 | Genetic obesity (fa/fa) | | | | 57.82 ± 1.24 nm§ (Mean ± SE) |
| | | | 6 | Genetic obesity + exercise (11–18 wk) | | | | 63.70 ± 1.29 nm (Mean ± SE) |
| | | | 6 | Genetic obesity + exercise (6–18 wk) | | | | 65.57 ± 1.48 nm (Mean ± SE) |
| 32 | Male Zucker rat | 68 wk (17 mo) | 5 | Lean (FA/FA) + standard chow | ? | Retina | EM | 93.6 ± 6.12 nm§ |

| | | | | | | | | | |
|---|---|---|---|---|---|---|---|---|---|
| | | | 4 | Obese (fa/fa) + standard chow | | | | | 104.6 ± 4.58 nm§ |
| | | | 6 | Lean (FA/FA) + Sucrose | | | | | 97.4 ± 2.82 nm |
| | | | 3 | Obese (fa/fa) + Sucrose | | | | | 99.6 ± 9.78 nm |
| 33 | Male albino Wistar rat | 6 mo | 1 | Normal | 334.5 ± 16.8 g | Epididymal adipose tissue | Basement membrane | EM + Image J by Yunjeong Lee, this study | 109±11 nm (mean ± SE) |
| | | | 5 | | | | Lamina lucida | EM | 21.7 ± 2.5 nm |
| | | | | | | | Basal lamina (Lamina densa) | | 43.1 ± 3.0 nm |
| | | 24 mo | 5 | | 358.5 ± 14.7 g | Epididymal adipose tissue | Lamina lucida | EM | 21.9 ± 2.1 nm |
| | | | | | | | Basal lamina (Lamina densa) | | 26.9 ± 2.6 nm |
| 34 | Male Zucker rat | 6–7 mo | 4 | Lean (Fa/fa) | ? | Retina | | TEM | 89.0 ± 1.958 nm§ (mean ± SE) |
| | | | 4 | Genetic obesity & diabetes (fa/fa) | | | | | 113.4 ± 1.78 nm§ (mean ± SE) |
| 35 | Sprague-Dawley rat | > 6 mo | 6 | Normal | ? | Retina | | EM | 50.8 ± 5.1 nm§ |
| | | | | | | Kidney | | | 81 ± 9.9 nm§ |
| | | | 6 | Diabetes | | Retina | | | 69.2 ± 15.9 nm |
| | | | | | | Kidney | | | 111.4 ± 25.2 nm |
| | | | 6 | Tightly controlled diabetes | | Retina | | | 53.6 ± 3.3 nm |
| | | | | | | Kidney | | | 86.7 ± 6.3 nm |
| 36 | Male Sprague-Dawley rat | 8 mo + 6–7 wk (9–10 mo) | 3 | Normal | 694.5 ± 17.2 g | Kidney | | EM | 235.57 ± 1.05 nm§ (mean ± SE) |
| | | | 3 | STZ-induced diabetes | 380.2 ± 11.9 g | | | | 346.19 ± 1.11 nm (mean ± SE) |
| | | | 3 | STZ-induced diabetes + low molecular weight heparin | 378.8 ± 12.7 g | | | | 221.85 ± 1.08 nm (mean ± SE) |
| | | | 3 | STZ-induced diabetes + Dermatan sulphate | 363.4 ± 18.5 g | | | | 260.08 ± 1.06 nm (mean ± SE) |
| 37 | Wistar rat | 4 wk (1 mo) | 4 | Normal | 109 ± 14 g | Kidney | | EM | 119.0 ± 6.8 nm§ |
| | | 8 wk | 4 | Normal | 224 ± 17 g | | | | 129.1 ± 4.6 nm§ |

| | | | | | | | | |
|---|---|---|---|---|---|---|---|---|
| | | (2 mo) | | | | | | |
| | | 12 wk (3 mo) | 4 | Normal | 278 ± 13 g | | | 135.4 ± 3.5 nm§ |
| | | 16 wk (4 mo) | 4 | Normal | 313 ± 10 g | | | 147.3 ± 1.3 nm§ |
| | | 20 wk (5 mo) | 4 | Normal | 345 ± 18 g | | | 154.1 ± 3.5 nm§ |
| | | 24 wk (6 mo) | 4 | Normal | 404 ± 48 g | | | 160.5 ± 3.8 nm§ |
| | | 28 wk (7 mo) | 4 | Normal | 488 ± 41 g | | | 184.6 ± 6.5 nm§ |
| | | 32 wk (8 mo) | 4 | Normal | 563 ± 48 g | | | 205.3 ± 7.2 nm§ |
| | | 36 wk (9 mo) | 4 | Normal | 624 ± 40 g | | | 238.3 ± 11.4 nm§ |
| 38 | Male Wistar rat | >14 mo+11 wk (total 17 mo) | 6 | STZ-induced diabetes + normal diet | 436 ± 14 g | Kidney | EM | 431 ± 18 nm (mean ± SE) |
| | | | 6 | STZ-induced diabetes + low carbohydrate diet | 444 ± 29 g | | | 389 ± 15 nm (mean ± SE) |
| | | | 6 | STZ-induced diabetes + insulin + normal diet | 459 ± 10 g | | | 414 + 8 nm (mean ± SE) |
| | | | 5 | STZ-induced diabetes + insulin + low carbohydrate diet | 604 ± 50 g | | | 400 ± 26 nm (mean ± SE) |
| | | | 6 | Normal + normal diet | 598 ± 24 g | | | 305 ± 10 nm§ (mean ± SE) |
| | | | 6 | Normal + low carbohydrate diet | 655 ± 35 g | | | 349 ± 22 nm (mean ± SE) |
| 39 | Male OLEFT rat vs. LETO rat | 22 wk (5.5 mo) | 50 (5 rats) | Normal | 506 ± 22 g | Heart | EM | 90 ± 12 nm§ |
| | | 62 wk (15.5 mo) | 50 (5 rats) | | 528 ± 18 g | | | 87 ± 12 nm§ |
| | | 22 wk (5.5 mo) | 50 (5 rats) | Obese with diabetes | 644 ± 32 g | | | 106 ± 20 nm§ |
| | | 62 wk (15.5 mo) | 50 (5 rats) | | 523 ± 90 g | | | 177 ± 66 nm§ |

| | | | | | | | | | |
|---|---|---|---|---|---|---|---|---|---|
| 40 | Rat | ? | 6 | Normal | ? | Heart (left ventricular) | | EM | 68.7 ± 4.22 nm§ (mean ± SE) |
| | | ? | 5 | acute myocardial infarction | ? | | | | 76 ± 9.85 nm (mean ± SE) |
| 41 | Female Wistar-Kyoto albino rat | 10.5 mo (age 6 wk, diet 9 mo) | 5 | Normal | ? | Retina | Inner nuclear layer | EM | 166.7 ± 44.5 nm§ |
| | | | 5 | | ? | | Nerve fiber layer | | 206.5 ± 42.3 nm§ |
| | | | 5 | Galactose fed | ? | | Inner nuclear layer | | 280.6 ± 69.7 nm |
| | | | 5 | | ? | | Nerve fiber layer | | 316.4 ± 80.3 nm |
| | | | 5 | Galactose + sorbinil fed | ? | | Inner nuclear layer | | 164.0 ± 31.2 nm |
| | | | 5 | | ? | | Nerve fiber layer | | 214.6 ± 50.2 nm |
| 42 | Male Sprague-Dawley rat | >28 wk (>7 mo) | 3 | Control | ? | Retina (outer plexiform layer) | | EM | 96.7 ± 14.1 nm§ |
| | | | 3 | Galactose fed | ? | | | | 151.9 ± 18.7 nm |
| | | | 3 | Galactose + sorbinil fed | ? | | | | 99.1 ± 14.1 nm |
| | | >44 wk (>11 mo) | 4 | Control | ? | | | | 93.9 ± 12.3 nm§ |
| | | | 4 | Galactose fed | ? | | | | 194.4 ± 40.4 nm |
| | | | 4 | Galactose + sorbinil fed | ? | | | | 105.8 ± 17.2 nm |
| 43 | Male Sprague-Dawley rat | >7 mo | 5 | Control | ~200 g (beginning of the experiment) | Retina (outer plexiform layer) | | EM | 85 ± 20 nm§ |
| | | | 5 | Galactose fed | | | | | 139 ± 16 nm |
| | | | 5 | Galactose fed + antisense oligos | | | | | 105 ± 24 nm |
| 44 | Male Sprague-Dawley rat | >6 mo | 8 | Control | 631.8 ± 23.51 g | Retina (outer plexiform and inner nuclear layer) | | EM | 66.142 ± 2.756 nm§ (mean ± SE) |
| | | | 8 | Diabetes | 495.2 ± 15.2 g | | | | 113.268 ± 5.236 nm (mean ± SE) |
| | | | 8 | Diabetes with bosentan‡‡ | 484.9 ± 9.8 g | | | | 84.606 ± 3.307 nm (mean ± SE) |
| | | | 8 | Galactose-fed | 542.2 ± 17.2 g | | | | 119.331 ± 4.685 nm (mean ± SE) |
| | | | 8 | Galactose with bosentan | 529.2 ± 9.2 g | | | | 84.055 ± 4.409 nm (mean ± SE) |

| | | | | | | | | | |
|---|---|---|---|---|---|---|---|---|---|
| 45 | Male Sprague-Dawley rat | >32 wk (>8 mo) | 8 | Control | 250–300 g | Retina (outer plexiform layer) | | TEM | 75.7 ± 10.5 nm[§] |
| | | | 8 | Control treated with propranolol | | | | | 73.4 ± 7.90 nm |
| | | | 8 | Control treated with fosenopril sodium | | | | | 73.7 ± 9.87 nm |
| | | | 8 | Diabetes | | | | | 171 ± 19.4 nm |
| | | | 8 | Diabetes treated with propranolol | | | | | 164 ± 15.8 nm |
| | | | 8 | Diabetes treated with fosenopril sodium | | | | | 93 ± 11.2 nm |
| 46 | Male Wistar Kyoto rat | 27 wk (<7 mo) | 7 | Control (normotensive non-diabetic) | 429 ± 21 g | Retina (outer plexiform layer) | | EM | 111.8 ± 14.2 nm[§] |
| | | | 7 | normotensive diabetes | 165 ± 11 g | | | | 132.8 ± 19.4 nm |
| | | | 7 | normotensive diabetes + cilazapril | 160 ± 22 g | | | | 131.9 ± 17.3 nm |
| | Male spontaneously hypertensive rat | | 7 | hypertensive diabetic | 175 ± 15 g | | | | 150.3 ± 20.2 nm |
| | | | 7 | hypertensive diabetes + cilazapril | 190 ± 28 g | | | | 116.7 ± 11.0 nm |
| 47 | Female normotensive Wistar Kyoto | 6 mo | 4 | Control | ? | Retina (inner nuclear and inner plexiform layers) | | Light microscopy | 81.58 ± 7.89 nm[§] (mean ± SE) |
| | | 9 mo | 4 | | ? | | | | 94.74 ± 7.89 nm[§] (mean ± SE) |
| | | 12 mo | 4 | | ? | | | | 121.1 ± 19.3 nm[§] (mean + SE) |
| | | 6 wk+ 15–21 mo (16.5–22.5 mo) | 7 | | 331 ± 4 g | | | | 160.4 ± 31.2 nm[§] |
| | | | 7 | 30% galactose-fed | 285 ± 28 g | | | | 203.3 ± 37.6 nm |
| | | | 9 | 30% galactose-fed + sorbinil | 282 ± 11 g | | | | 159.4 ± 35.2 nm |
| | Female spontaneously hypertensive rat | 6 wk + 15–21 mo (16.5–22.5 mo) | 7 | Control | 30–40 g lighter than WKY rats | | | | 159.4 ± 36.1 nm |
| | | | 6 | 30% galactose-fed | | | | | 230.7 ± 62.3 nm |
| | | | 8 | 30% galactose-fed + sorbinil | | | | | 154.3 ± 43.5 nm |
| 48 | Male prediabetic diabetes-prone BB rat | >3 wk+6 mo (>7 mo) | 5 | Diabetes | 306 g | Retina | Superficial | EM | 192.2 ± 5.8 nm (mean ± SE) |
| | | | 5 | | | | Deep | | 139.9 ± 5.8 nm (mean ± SE) |
| | | | 5 | Diabetes + ponalrestat | 275 g | | Superficial | | 175.7 ± 4.5 nm (mean ± SE) |
| | | | 5 | | | | Deep | | 99.9 ± 9.2 nm |

| | | | | | | | | | |
|---|---|---|---|---|---|---|---|---|---|
| | | | | | | | | | (mean ± SE) |
| | | | 5 | Diabetes + insulin | 366 g | | Superficial | | 134.0 ± 6.9 nm (mean ± SE) |
| | | | 5 | | | | Deep | | 97.7 ± 7.0 nm (mean ± SE) |
| | Male non-diabetes- prone male BB rat | | 5 | Control | 495 g | | Superficial | | 134.6 ± 7.7 nm (mean ± SE) § |
| | | | 5 | | | | Deep | | 95.1 ± 3.3 nm (mean ± SE) § |
| | | | 5 | Control + ponalrestat | 456 g | | Superficial | | 127.9 ± 6.9 nm (mean ± SE) |
| | | | 5 | | | | Deep | | 82.8 ± 8.5 nm (mean ± SE) |
| 49 | Female Wistar rat | >10 wk+8 mo (>10.5 mo) | 6 | Control | 324 ± 7 g | Retina (outer plexiform layer) | | EM | 84.4 ± 1.16 nm§ (mean ± SE) |
| | | | 12 | Diabetes | ? | | | | 124 ± 5.42 nm (mean ± SE) |
| | | | 12 | Diabetes + tolrestat | ? | | | | 98.3 ± 4.65 nm (mean ± SE) |
| 50 | Male Sprague-Dawley rat | >6 mo | ? | Control | Approx. 550 g | Inner ear | | EM | 102 ± 9 nm (mean ± SE) |
| | | | ? | Diabetes | Approx. 350 g | | | | 177 ± 10 nm (mean ± SE) |
| | | | ? | Noise-exposed control | Approx. 470 g | | | | 97 ± 1 nm (mean ± SE) |
| | | | ? | Noise-exposed diabetes | Approx. 330 g | | | | 175 ± 5 nm (mean ± SE) |
| 51 | Male CRL:COBS-CD (SD) Sprague-Dawley rat | >7 mo | 4 | Control | ? | Retina (outer plexiform layer) | | EM | 122.9 ± 19.6 nm§ |
| | | | 4 | 50% galactose-fed | ? | | | | 200.1 ± 39.9 nm |
| | | | 4 | 50% galactose-fed + 0.03% tolrestat | ? | | | | 130.1 ± 19.6 nm |
| | | | 4 | 50% galactose-fed + 0.04% tolrestat | ? | | | | 120.3 ± 15.7 nm |
| 52 | Male Lewis rat of the AC1 (AgB4/4) strain | >4 mo | 5 | Control | 329 ± 29 g | Retina Superficial capillary bed | | EM | 146.2 ± 6.7 nm§ |
| | | | | | | Retina Deep capillary bed | | | 97.4 ± 7.5 nm§ |
| 53 | Male Sprague-Dawley rat | >20 mo | 13 | Control | ? | Retina (outer plexiform layer or inner nuclear layer) | | EM | 159 ± 23 nm§ |
| | | | 18 | Diabetes | ? | | | | 216 ± 36 nm |
| | | | 6 | 30% galactose | ? | | | | 227 ± 37 nm |
| | | | 6 | 50% galactose | ? | | | | 276 ± 26 nm |
| 54 | BB-rat | >6 mo | ? | Control | 539.0 ± 21.6 g | Superficial retinal capillary | | EM | 161.9 ± 3.7 nm§ (mean ± SE) |

| | | | | | | | | |
|---|---|---|---|---|---|---|---|---|
| | | | | | | Deep retinal capillary | | 142.7 ± 3.3 nm§ (mean ± SE) |
| | | | | | | Muscle | | 125.0 ± 3.2 nm§ (mean ± SE) |
| | | | | | | Endoneurial capillary | | 142.8 ± 4.5 nm (mean ± SE) |
| | | | | | | Kidney | | 326.0 ± 20.0 nm§ (mean ± SE) |
| 55 | Male Sprague Dawley rat | >6 mo | 8 | Control | ? | Retina (outer plexiform layer) | EM | 51.5 ± 4.8 nm§ |
| | | | 8 | Diabetes | ? | | | 72.5 ± 5.0 nm |
| | | | 8 | Diabetes + FN-siRNA§§ | ? | | | 56.4 ± 2.8 nm |
| | | | 8 | Diabetes + scrambled siRNA | ? | | | 73.7 ± 3.9 nm |
| 56 | Male Sprague-Dawley rat | 33 wk (8 mo) | 6 | Control | 577.4 ± 36.1 g | Retina (outer plexiform and ganglion cell layers) | TEM | 94.6 ± 7.7 nm§ |
| | | | 6 | Diabetic | 343.5 ± 31.3 g | | | 212.4 ± 18.1 nm |
| | | | 6 | Diabetic + fenofibrate | 381.2 ± 28.7 g | | | 129.3 ± 9.4 nm |
| 57 | Wistar rat | 6 mo | ? | Control | ? | Heart LV (basal lamina was measured) | EM | 44.9 ± 3.2 nm |
| | | | ? | Diabetic | ? | | | 49.9 ± 3.8 nm |
| | | | ? | Diabetic+EGb 761 treatment | ? | | | 48.2 ± 1.5 nm |
| 58 | Human | 48±12 y | 6 | Lean | ? | Visceral adipose tissue | TEM | 103.38 nm [67.23, 194.26] (Median, 95% percentile) |
| | | 41±9 y | 5 | Obese without diabetes | | | | 108.78 nm [60.47, 181.76] (Median, 95% percentile) |
| | | 45±10 y | 5 | Obese with prediabetes | | | | 116.56 nm [68.79, 208.28] (Median, 95% percentile) |
| | | 52±9 y | 5 | Obese with type 2 diabetes | | | | 139.87 nm [68.24, 209.12] (Median, 95% percentile) |
| 59 | Human | ? | 15 | Normal | ? | Muscle | EM | 158 ± 10.3 nm (mean ± SE) |
| | | | 9 | Prediabetes | | | | 163 ± 5.5 nm |

| | | | | | | | | |
|---|---|---|---|---|---|---|---|---|
| | | | | | | | | (mean ± SE) |
| | Dog | ? | 21 | Normal | ? | Abdominal muscle | | 86 ± 14 nm (mean ± SE) |
| | | | 12 | | | Thigh muscle | | 117.5 ± 10.5 nm (mean ± SE) |
| | | 5 y | 3 | Alloxan-induced diabetic | ? | Abdominal muscle | | 340 ± 51.8 nm (mean ± SE) |
| | | 5 y | 2 | Metasomatotrophin-induced diabetic | ? | Abdominal muscle | | 215 ± 35 nm (mean ± SE) |
| | | 3 y | 2 | Alloxan-induced diabetic | ? | Abdominal muscle | | 285 ± 15 nm (mean ± SE) |
| | | | 6 | | | Thigh muscle | | 187 ± 14 nm (mean ± SE) |
| | | 0–16 mo | ? | Normal | ? | Kidney | | 113 nm, 594 nm (min, max) |
| | | | ? | | | Retina | | 60.4 nm, 441 nm (min, max) |
| | | 0–17 mo | ? | | ? | Fat | | 33.8 nm, 74.1 nm (min, max) |
| | Rat | 0–19 mo | ? | Normal | ? | Kidney | | 102 nm, 377 nm (min, max) |
| | | | ? | | | Retina | | 37.5 nm, 198 nm (min, max) |

†SD: Standard deviation

‡TEM: Transmission electron microscopy

§Data were included in the analysis

¶EM: Electron microscopy

#Heart IVS: Interventricular septal sample

††Heart LV: Left ventricular sample

‡‡Endothelin receptor blocker

§§FN: Fibronectin

**Table S7. Studies that measured binding affinities of VEGF-A165 to VEGFR1**

| Reference | Ligand | Receptor | Method | Ligand/receptor source | $k_{on}$ ($M^{-1}s^{-1}$) | $k_{off}$ ($s^{-1}$) | $K_d$ (Mean ± SE) |
|---|---|---|---|---|---|---|---|
| 60 | Recombinant $^{125}$I-VEGF-A165 expressed in baculovirus system | Human VEGFR1 on Porcine aortic endothelial (PAE) cells or HUVECs | Radioligand (Competitive binding) | PAE cells transfected with a VEGFR1-expressing vector | ? | ? | 16 pM† |
| | | | | HUVECs | | | 9 pM† |
| 61 | Recombinant VEGF-A165 | Immobilized recombinant human VEGFR1 protein | SPR | Obtained from R&D Systems | $(4.0 \pm 0.04) \times 10^{5}$† | $(4.0 \pm 0.1) \times 10^{-7}$† | 1 ± 0.3 pM† |
| 62 | Recombinant VEGF-A165 | Immobilized human VEGFR1 Fc chimera‡ | SPR | Obtained from R&D Systems | $(2.91 \pm 0.04) \times 10^{6}$ | $(5.69 \pm 0.07) \times 10^{-4}$ | 196 ± 4 pM |
| 63 | VEGF-A165 | Immobilized sVEGFR1§ | SPR | VEGF-A165 & VEGFR1: Sf158 insect cells infected with a baculovirus-based vector (both ligand and receptor) | $(4.0 \pm 0.38) \times 10^{6}$† | $(3.0 \pm 0.25) \times 10^{-5}$† | 7.5 ± 0.95 pM† |
| | | sVEGFR1§ on microtiter plates | Radioligand (Saturation analysis; Scatchard analysis) | | ? | ? | 74 ± 7.4 pM† |
| 64 | VEGF-A165 | Human VEGFR1-hFc¶ | SPR | VEGF-A165: Made at Regeneron Pharmaceuticals<br><br>Human VEGFR1-hFc: Obtained from R&D systems | $(3.0 \pm 0.2) \times 10^{7}$† | $(2.8 \pm 0.1) \times 10^{-4}$† | 9.33 ± 0.956 pM† |
| 65 | $^{125}$I-VEGF-A165 | Rat VEGFR1 on adventitial fibroblasts | Radioligand (Saturation analysis; Scatchard analysis) | VEGF-A165: Obtained from R&D Systems<br><br>VEGFR1: Adventitial fibroblasts from thoracic aorta of 6–8-week-old male Sprague–Dawley rats | ? | ? | 7±1 pM† |

| | | | | | | | | |
|---|---|---|---|---|---|---|---|---|
| 66 | Recombinant $^{125}$I-VEGF-A165 | Murine VEGFR1 on COS cells | Radioligand (Saturation analysis) | Non-linear regression curve | VEGF-A165: Expressed in Sf9 insect cells<br><br>VEGFR1: COS cells transfected with murine full length VEGFR1 cDNA | ? | ? | 90 pM |
| | | | | Scatchard analysis | | | | 114 pM† |
| 67 | VEGF-A165 | Human VEGFR1 on PAE cells | Radioligand (Saturation analysis; Scatchard) | No heparin | VEGF-A165: From PeproTech. Inc<br><br>VEGFR1: PAE cells transfected with human VEGFR1 cDNA | ? | ? | 54 pM† |
| | | | | Heparin 0.5 μg/ml | | | | 77 pM |
| | | | | Heparin 5 μg/ml | | | | 118 pM |
| 68 | Recombinant $^{125}$I-VEGF-A165 | Human VEGFR1-Fc chimera | ELISA plate + saturation analysis (Scatchard) | | VEGF-A165: Expressed in Sf9 insect cells<br><br>VEGFR1: R&D Systems | ? | ? | 59.4±19.6 pM† |
| 69 | α5β1 integrin§ | Recombinant human VEGFR1 | SPR | | VEGFR1: R&D Systems<br><br>α5β1 integrin: Merck Millipore | $1.5 \times 10^3$ | $2.9 \times 10^{-4}$ | 195 ± 40 nM |
| 70 †† | Recombinant $^{125}$I-VEGF-A165 | Possibly Human VEGFR1 | Normal cell line 001 | Radioligand (Saturation analysis; Scatchard) | VEGF-A165: Purified from a baculovirus expression system<br><br>VEGFR1: Human retinal pigment epithelial (HRPE) line | ? | ? | 5 pM |
| | | | Normal cell line 002 | | | | | 6 pM |
| | | | Normal cell line 003 | | | | | 8 pM |
| | | | Immortalized cell line 004 | | | | | 9 pM |
| 71 | - | Human VEGFR1 | This study did not perform binding assays. | | | | | |

†Data was included in the analysis.
‡Fc chimera: pre-dimerized fusion proteins consisting of VEGFRs and the constant Fc region of human IgG1
§Species were not specified.
¶The extracellular domains of dimerized human VEGFR fused inline to hFc.
††This study was excluded because HRPE cells express VEGFR1, VEGFR2, and NRP1 and we could not differentiate the binding affinities of the receptors.

**Table S8. Studies that measured binding affinities of VEGF-A165 to VEGFR2**

| Reference | Ligand | Receptor | Method | Ligand/receptor source | $k_{on}$ ($M^{-1}s^{-1}$) | $k_{off}$ ($s^{-1}$) | $K_d$ (Mean ± SE) |
|---|---|---|---|---|---|---|---|
| 72 | Recombinant VEGF-A165 (50 nM) | Recombinant sVEGFR2 (Full length mouse Flk-1 cDNA) | SPR | VEGF-A165: R&D Systems<br><br>VEGFR2: Spodoptera frugiperda (Sf9) infected with sVEGFR2 recombinant baculovirus | $1.2 \times 10^{6\dagger}$ | $4.1 \times 10^{-4\dagger}$ | 340 pM$^{\dagger}$ |
| | Recombinant VEGF-A165 (5 nM) | | | | $2.2 \times 10^{6\dagger}$ | $2.4 \times 10^{-4\dagger}$ | 110 pM$^{\dagger}$ |
| | Recombinant VEGF-A164 (20 nM) | | | | $1.9 \times 10^{6\dagger}$ | $6.2 \times 10^{-4\dagger}$ | 330 pM$^{\dagger}$ |
| | Recombinant VEGF-A164 (5 nM) | | | | $2.2 \times 10^{6\dagger}$ | $3.0 \times 10^{-4\dagger}$ | 140 pM$^{\dagger}$ |
| 73 | Carrier-free recombinant VEGF-A165 | Human VEGFR2 | Radioligand (Saturation analysis; Nonlinear regression) | VEGF-A165: R&D Systems<br><br>VEGFR2: COS-1 cells transiently transfected with human VEGFR2 cDNA | ? | ? | $339 \pm 120$ pM$^{\dagger}$ |
| | | | | VEGF-A165: R&D Systems<br><br>NRP1: Balb/c cells transiently transfected with human VEGFR2 cDNA | ? | ? | $291 \pm 54.4$ pM$^{\dagger}$ |
| 60 | Recombinant $^{125}$I-VEGF-A165 expressed in baculovirus system | Human VEGFR2 on PAE cells or HUVECs | Radioligand (Scatchard analysis) | PAE cells transfected with a VEGFR2-expressing vector | ? | ? | 760 pM$^{\dagger}$ |
| | | | | HUVECs | | | 770 pM$^{\dagger}$ |
| 64 | VEGF-A165 | Human VEGFR2-hFc | SPR | VEGF-A165: Made at Regeneron Pharmaceuticals<br><br>Human VEGFR2-hFc: Obtained from R&D systems | $(1.52 \pm 0.05) \times 10^{7\dagger}$ | $(1.35 \pm 0.06) \times 10^{-3\dagger}$ | $88.8 \pm 6.87$ pM$^{\dagger}$ |
| 74 | VEGF-A165 | Recombinant VEGFR2 Fc$^{\ddagger\P}$ | SPR | VEGFR2: | $(3.6 \pm 0.07) \times 10^{6\dagger}$ | $(1.34 \pm 0.19) \times 10^{-4\dagger}$ | $37.1 \pm 4.9$ pM$^{\dagger}$ |

| | | | | | | | |
|---|---|---|---|---|---|---|---|
| | VEGF-A165 | Recombinant VEGFR2 cbu§¶ | | SF21 cells expressing VEGFR2 Fc or cbu | $(5.23 \pm 1.4) \times 10^{6\dagger}$ | $(2.74 \pm 0.76) \times 10^{-4\dagger}$ | $51.7 \pm 5.8$ pM† |
| | VEGF-A165 | First three immunoglobulin-like domains of VEGFR2¶ | | | $(4.72 \pm 1.0) \times 10^{6\dagger}$ | $(0.67 \pm 0.11) \times 10^{-4\dagger}$ | $14.5 \pm 1.0$ pM† |
| 61 | Recombinant VEGF-A165 | Immobilized recombinant human VEGFR2 protein | SPR | Obtained from R&D Systems | $(9.7 \pm 0.3) \times 10^{5\dagger}$ | $(9.5 \pm 0.2) \times 10^{-6\dagger}$ | $9.8 \pm 0.4$ pM† |
| 62 | Recombinant VEGF-A165 | Immobilized human VEGFR2 Fc chimera | SPR | Obtained from R&D Systems | $(1.76 \pm 0.04) \times 10^{6}$ | $(1.51 \pm 0.07) \times 10^{-2}$ | $8.6 \pm 0.5$ nM |
| 68 | Recombinant $^{125}$I-VEGF-A165 | Human VEGFR2-Fc chimera | ELISA plate + saturation analysis (Scatchard) | VEGF-A165: Expressed in Sf9 insect cells<br><br>VEGFR2: R&D Systems | ? | ? | $292.5 \pm 163.8$ pM† |
| 75 | Recombinant $^{125}$I-VEGF-A165 | Human VEGFR2 on CMT-3 cells | Radioligand (Saturation analysis; Scatchard) | VEGF-A165: Cells transfected with an expression vector containing the VEGF cDNA encoding the 165 amino acid form of VEGF<br><br>VEGFR2: CMT-3 monkey kidney cells transfected with KDR gene | ? | ? | 75 pM† |
| 76 | Recombinant VEGF-A165 | Recombinant human VEGFR2 | SPR | VEGF-A165: PeproTech<br><br>VEGFR2 R&D Systems | $(3.58 \pm 1.44) \times 10^{7\dagger}$ | $(4.13 \pm 0.28) \times 10^{-3\dagger}$ | $115 \pm 73.44$ pM† |
| In-house data, 2023 | Recombinant VEGF-A165 | Recombinant human VEGFR2 | SPR | VEGF-A165: R&D Systems<br><br>VEGFR2: R&D Systems | $(6.24 \pm 0.46) \times 10^{5\dagger}$ | $(3.18 \pm 1.98) \times 10^{-4\dagger}$ | $520 \pm 250$ pM† |
| 77 | Recombinant $^{125}$I-VEGF-A165 | VEGFR2 on HUVEC | Radioligand (Saturation analysis; Scatchard) | VEGF-A165: Sf-9 insect cells infected with a baculovirus-based | ? | ? | 7.5 pM† |

<table>
<tr><td></td><td></td><td colspan="2"></td><td></td><td>vector expressing VEGF-165 cDNA</td><td></td><td></td><td></td></tr>
<tr><td>78</td><td>$^{125}$I-VEGF-A165</td><td colspan="2">Mouse VEGFR2 on NIH-3T3 cells</td><td>Radioligand (Saturation analysis; Scatchard)</td><td>VEGF-A165:<br>Sf-9 insect cells infected with a baculovirus-based expression vector for VEGF165<br><br>VEGFR2:<br>The DNA encoding the entire chimera was then subcloned into the pMFG expression vector and expressed in NIH-3T3 cells.</td><td>?</td><td>?</td><td>19 pM†</td></tr>
<tr><td>79</td><td>VEGF-A121</td><td colspan="2">Recombinant human VEGFR2 extracellular domain</td><td>SPR</td><td>VEGF-A121:<br>R&D Systems<br><br>VEGFR2:<br>Calbiochem EMD Chemicals</td><td>$(3.3 \pm 0.2) \times 10^5$</td><td>$(1.0 \pm 0.1) \times 10^{-3}$</td><td>$3.1 \pm 0.4$ nM</td></tr>
<tr><td rowspan="4">70††</td><td rowspan="4">Recombinant $^{125}$I-VEGF-A165</td><td rowspan="4">Possibly Human VEGFR2</td><td>Normal cell line 001</td><td rowspan="4">Radioligand (Saturation analysis; Scatchard)</td><td rowspan="4">VEGF-A165:<br>Purified from a baculovirus expression system<br><br>VEGFR2:<br>HRPE line</td><td rowspan="4">?</td><td rowspan="4">?</td><td>68 pM</td></tr>
<tr><td>Normal cell line 002</td><td>95 pM</td></tr>
<tr><td>Normal cell line 003</td><td>105 pM</td></tr>
<tr><td>Immortalized cell line 004</td><td>210 pM</td></tr>
</table>

†Data were included in the analysis.
‡ Fc: pre-dimerized fusion protein
§ cbu: monomeric fusion protein
¶Species were not specified.
††This study was excluded because HRPE cells express VEGFR1, VEGFR2, and NRP1 and we could not differentiate the binding affinities of the receptors.

**Table S9. Studies that measured binding affinities of VEGF-A165 to NRP1**

| Reference | Ligand | Receptor | Method | Ligand/receptor source | $k_{on}$ ($M^{-1}s^{-1}$) | $k_{off}$ ($s^{-1}$) | $K_d$ (Mean ± SE) |
|---|---|---|---|---|---|---|---|
| 73 | Carrier-free recombinant VEGF-A165 | Human NRP1 | Radioligand (Saturation analysis; Nonlinear regression) | VEGF-A165: R&D Systems<br>NRP1: COS-1 cells transiently transfected with human NRP1 cDNA | ? | ? | 2.09 ±0.818 nM[†] |
| | | | | VEGF-A165: R&D Systems<br>NRP1: Balb/c cells transiently transfected with human NRP1 cDNA | | | 417 ± 125 pM[†] |
| 77 | Recombinant $^{125}$I-VEGF-A165 | NPR1 on HUVEC | Radio-ligand (Saturation analysis; Scatchard); No heparin | VEGF-A165: Sf-9 insect cells infected with a baculovirus-based vector expressing VEGF-165 cDNA | ? | ? | 200 pM[†] |
| | | NRP1 on breast cancer cell (MDA-MB-231) | | | | | 280 pM[†] |
| | | | + heparin | | | | 270 pM |
| 80 | Recombinant $^{125}$I-VEGF-A165 | Human NRP1 on PAE cells | Radioligand (Scatchard analysis) | VEGF-A165: Sf-21 insect cells infected with recombinant baculovirus vectors<br>NRP1: PAE cells transfected with NRP1 cDNA | ? | ? | 320 pM[†] |
| 81 | Biotinylated VEGF-A165 | First 600 amino acids of mouse NPR-1 extracellular domain, which lacked C-terminal MAM domain (immobilized) | SPR Low density (350 RU) | NRP1: Transfected D. melanogaster cells | < (0.1–1) × $10^6$ | < $10^{-2}$ | 2,000 nM |
| | | | SPR High density (1400 RU) | | | | 113 nM |
| | | | ELISA No heparin | | | | 120 nM ($IC_{50}$) |
| | | | ELISA Add heparin | | | | 25 nM ($IC_{50}$) |
| 82 | VEGF-A165 | Human sNRP1-Fc[‡] | SPR (Steady-state analysis) | VEGF-A165: | ? | ? | 120 nM |

| | | | | | | | |
|---|---|---|---|---|---|---|---|
| | | | | Purchased from R&D Systems<br><br>sNRP1:<br>Transfected Chinese hamster ovary cells | | | |
| [62] | Recombinant VEGF-A165 | Immobilized rat NRP-1 Fc chimera | SPR | Obtained from R&D Systems | $(3.6 \pm 0.2) \times 10^6$ | $(9.0 \pm 0.4) \times 10^{-2}$ | $25 \pm 2$ nM |
| | | Immobilized mouse sNRP-1 monomer (only ECD of the mouse sequence) | | | $(2.64 \pm 0.08) \times 10^6$ | $(6.5 \pm 0.2) \times 10^{-2}$ | $25 \pm 1$ nM |
| [83] | Recombinant $^{125}$I-VEGF-A165 | Rat NRP1 on COS-1 cells | Radioligand (Saturation analysis; Nonlinear regression) | NRP1:<br>COS-1 cells transiently expressing Npn-1 | ? | ? | $0.93 \pm 0.71$ nM$^{\dagger}$ |
| [68] | Recombinant $^{125}$I-VEGF-A165 | Human NRP1-Fc chimera$^{\S}$ | ELISA plate + saturation analysis (Scatchard) | VEGF-A165:<br>expressed in Sf9 insect cells<br><br>NRP1:<br>R&D Systems | ? | ? | $246.4 \pm 135.1$ pM$^{\dagger}$ |
| [84] | Recombinant VEGF-A165 | Recombinant NRP1¶ | SPR | NRP1:<br>R&D Systems | $5 \times 10^5$ | $6 \times 10^{-4}$ | 1.2 nM |
| [85] | Recombinant human VEGF-A165 | Recombinant NRP1¶ | SPR | VEGF-A165:<br>R&D Systems<br><br>NRP1:<br>Purchased from an unknown vendor | $(1.88 \pm 0.53) \times 10^{6\dagger}$ | $(3.18 \pm 0.85) \times 10^{-3\dagger}$ | $1.694 \pm 1.834$ nM$^{\dagger}$ |
| [76] | Recombinant VEGF-A165 | Recombinant human NRP1 | SPR | VEGF-A165:<br>PeproTech<br><br>NRP1:<br>R&D Systems | $(5.55 \pm 2.33) \times 10^{7\dagger}$ | $(8.05 \pm 2.65) \times 10^{-3\dagger}$ | $145.1 \pm 58.91$ pM$^{\dagger}$ |
| In-house data, 2023 | Recombinant VEGF-A165 | Recombinant human NRP1 | SPR | VEGF-A165:<br>R&D Systems<br><br>NRP1:<br>R&D Systems | $(7.96 \pm 2.15) \times 10^{5\dagger}$ | $(1.56 \pm 0.55) \times 10^{-3\dagger}$ | $6.36 \pm 1.07$ nM$^{\dagger}$ |
| [86] | VEGF-A165 | Human NRP1 | Radioligand (Saturation analysis; Scatchard) | VEGF-A165: | ? | ? | 180 pM$^{\dagger}$ |

| | | | | | | | |
|---|---|---|---|---|---|---|---|
| | | | | produced in sF9 cells using appropriate baculoviruses<br><br>NRP1:<br>Transfected PAE cells with the pcDNA3/np-1 expression vector | | | |
| 87 | $^{125}$I-VEGF-A165 | NRP1-Fc¶ | Radioligand (Competitive assay) | VEGF-A165:<br>Amersham Biosciences<br><br>NRP1:<br>R&D Systems | ? | ? | 750 pM† |
| 69 | Recombinant human VEGFR1 | Recombinant human NRP1 | SPR | VEGFR1 and NRP1:<br>R&D Systems<br><br>α5β1 integrin:<br>Merck Millipore | $1.2 \times 10^{4}$ | $3 \times 10^{-4}$ | 25 ± 4 nM |
| | α5β1 integrin¶ | | | | $2.1 \times 10^{4}$ | $2.9 \times 10^{-4}$ | 14 ± 4 nM |
| | Semaphorin 3A¶ | | | | $3.7 \times 10^{3}$ | $9 \times 10^{-5}$ | 24 ± 6 nM |
| 88 | $^{125}$I-VEGF-A165 | Recombinant human NRP2 | Radioligand (Saturation assay; nonlinear regression) | VEGF-A165:<br>R&D Systems or National Cancer Institute<br><br>NRP2:<br>PAE cells overexpressing NRP2 | ? | ? | 5.2 ± 3.7 nM |
| 89 | $^{125}$I-VEGF-A165 | Recombinant human NRP1 b1b2 domain | Radioligand (Scatchard) | NRP1 domains:<br>PAE cells transfected with pSecTag mammalian expression vector | ? | ? | 30–42 nM |
| 90 | Recombinant $^{125}$I-VEGF-A165 | Human NRP1 on PAE cells | Radioligand (Saturation assay) | VEGF-A165:<br>Sf-21 insect cells infected with recombinant baculovirus vectors<br><br>NRP1:<br>PAE cells transfected with NRP1 cDNA | ? | ? | ? |
| 91 | VEGF-A165 fused to AP | Mouse NRP1 | AP ligand-binding assay | NRP1:<br>Hindbrain of E12.5 mouse embryo | ? | ? | ? |

†Data were used in the analysis.

‡Soluble NRP1 constructs, containing the NRP1 extracellular domain but lacking the MAM domain, were cloned into the expression vector pRK5 fused to the Fc portion of human IgG1 to facilitate affinity purification.
§The extracellular domain of NRP1.
¶Species were not specified.

**Table S10. Pairwise comparison of capillary basement membrane thickness in various tissues analyzed by Dunnett's T3 test**

| Comparison | | t-value | P-value (Prob[>\|t\|]) | Significance |
|---|---|---|---|---|
| Retina | Muscle | −1.81 | 0.555 | Not significant ($p > 0.05$) |
| | Heart | −3.44 | **0.031** | **Significant ($p < 0.05$)** |
| | Brain | −3.29 | **0.019** | **Significant ($p < 0.05$)** |
| | Kidney | 4.35 | **0.003** | **Significant ($p < 0.01$)** |
| Muscle | Heart | −0.81 | 0.990 | Not significant ($p > 0.05$) |
| | Brain | −0.39 | 1.000 | Not significant ($p > 0.05$) |
| | Kidney | 5.05 | **0.0005** | **Significant ($p < 0.001$)** |
| Heart | Brain | 0.67 | 0.998 | Not significant ($p > 0.05$) |
| | Kidney | 6.01 | **$8.5\times10^{-5}$** | **Significant ($p < 0.001$)** |
| Brain | Kidney | 5.90 | **0.0002** | **Significant ($p < 0.001$)** |
